# Direct activation of the Ras-Akt network mediates polarity and organizes protrusions in human neutrophil migration

**DOI:** 10.1101/2022.10.27.513966

**Authors:** Dhiman Sankar Pal, Tatsat Banerjee, Yiyan Lin, Jane Borleis, Peter N. Devreotes

## Abstract

Functions of Ras oncogenes and their downstream effectors are typically associated with cell proliferation and growth control while their role in immune cell migration has been largely unexplored. Although Ras-mediated signaling cascades have been implicated in immune response, there is no conclusive evidence to show local activation of these pathways on the plasma membrane directly regulates cell motility or polarity. Using spatiotemporally precise, cryptochrome-based optogenetic systems in human neutrophils, we abruptly altered protrusive activity, bypassing the chemoattractant-sensing receptor/G-protein network. First, global recruitment of active KRas4B/HRas isoforms or the guanine nucleotide exchange factor, RasGRP4, immediately increased spreading and random motility in neutrophils. Second, creating Ras activity at the cell rear generated new protrusions at the site and reversed pre-existing polarity, similar to the effects of steep chemoattractant gradients. Third, recruiting GTPase activating protein, RASAL3, at cell fronts abrogated existing protrusions and changed the direction of motility whereas dynamically inhibiting nascent fronts stopped migration completely. Fourth, combining pharmacological inhibition studies with optogenetics revealed that mTorC2 is more important than PI3K for Ras-mediated polarity and migration. Finally, local recruitment of Ras-mTorC2 effector, Akt, also generated new protrusions, rearranged pre-existing polarity, and triggered migration, even in absence of PI3K signaling. We propose that actin assembly, cell shape, and migration modes in immune cells are promptly controlled by rapid, local activities of established components of classical growth-control pathways independently of receptor activation.

## Introduction

Leukocytes, namely monocytes, neutrophils, and effector T-lymphocytes, sense chemical cues and rapidly migrate towards sites of tissue damage and infection^1–3^. This directed migration is coordinated by rearrangements of signal transduction proteins and lipids to the leading or trailing edges of the cell. This dictates the location and dynamics of cellular protrusions and contractions that move the cell^4–11^. Misdirected leukocyte migration is responsible for a wide array of autoimmune and inflammatory diseases such as Alzheimer disease, rheumatoid arthritis, juvenile diabetes, and asthma^1, 12, 13^. Discovering the signal transduction events that direct cytoskeletal activity, cellular protrusions, and migratory behavior in cells of the immune system will facilitate development of next generation treatments^14, 15^.

The current model for migration in neutrophils and other leukocytes is that chemoattractant mediated receptor/G-protein network stimulation activates downstream PI3Kγ signaling which triggers Rac to initiate actin polymerization, protrusion formation, and migration^9, 10, 16–35^. Although local PI3K activation does trigger cellular protrusions, other studies suggest that PI3Kγ signaling is not singularly essential for directed migration^20, 22, 36–44^. It is also unclear whether Akt, which is brought to the cell’s leading edge by PI3K/PIP3, is necessary for immune cell migration since it has been reported that Akt1 isoform is a negative regulator of motility whereas Akt2 promotes it^43, 45–48^. G-protein coupled receptors also activate Ras, which in turn stimulates mTorC2 and Akt, and is important for cell growth, survival, and energy metabolism^49–58^. It is unclear whether the Ras/mTorC2/Akt axis is needed for migration signaling. Knockout or knockdown studies with modulators of Ras activity have not given a conclusive understanding of Ras proteins in chemotaxis^50, 59^. Clearly, there is much uncertainty about the roles of these signaling pathways in directed migration. A sufficiently precise approach to directly link these activities to actin organization and cell polarity is lacking.

In the established model organism, *Dictyostelium discoideum,* role for Ras in migration has been more clearly defined using biochemical and genetic tools. Ras is activated by G-protein coupled receptor stimulation but also spontaneously at protrusions, which has not been shown in migrating mammalian cells^5, 6, 60–69^. However, as multiple Ras isoforms are expressed in these cells it has been difficult to demonstrate that Ras activity is important for migration and has prevented assignment of specific roles for each isoform by gene deletions^68, 70–72^. To circumvent this problem, researchers conditionally expressed constitutively active Ras isoforms which did produce phenotypes over many hours, allowing sufficient time for signaling networks to re-adjust through protein rearrangement or differential gene expression^61, 68, 73–75^. Miao and colleagues globally recruited signaling proteins to the plasma membrane within minutes and observed global increases or decreases in protrusive activity and motility depending on the particular node in the signaling networks targeted^76, 77^. Although these studies provided stronger evidence for involvement of Ras in migration, they could not show that local activation of these pathways on the membrane could bring about localized protrusions and directed migration. Moreover, these results were observed in the free-living amoeboid cells, and may or may not apply to the migratory cells of the human immune system.

New approaches with improved spatiotemporal resolution are needed to address critical questions which could not be answered with previous methods^78^. First, is Ras and its downstream effectors important for mammalian migration? Can activation of individual signaling components on the cell membrane influence cytoskeletal arrangement and affect motility? Does activation of different signaling nodes, such as Ras or Akt, have similar or opposing effects on cellular protrusions and migration? Second, chemoattractant gradients can bias pre-existing front-back axis of the cell, but the occupied receptors activate a vast array of downstream signaling events^79–85^. Can localized activation of individual components at the back override pre-existing polarity? Conversely, can localized inhibition of Ras at the cell front convert it to a back? Using sub-cellular optogenetics, we abruptly and locally perturbed Ras and Akt activity, bypassing chemoattractant-sensing receptor/G-protein network. Our studies answer these questions and show that cell shape, actin assembly, and migration modes in immune cells are controlled by local, spontaneous activities of the Ras/mTorC2/Akt axis of classical growth-control pathways.

## Results

For optical manipulation of signaling in migratory neutrophils, we engineered a blue light-inducible, cryptochrome-based dimerization system in the human HL-60 cell line. This process was carried out in a stepwise manner. First, the plasma membrane component CIBN, fused to a C-terminal CAAX motif, was expressed from an integrated lentiviral vector in wildtype neutrophils. Second, the F-actin polymerization biosensor, LifeAct tagged with an infrared fluorophore, was stably co-expressed to generate a dual-expressing cell line. Third, the cytosolic recruitable component, CRY2PHR-mcherry2, was stably introduced to these dual-expressing cells by transposon-based integration (Figure S1A). Illuminating the entire periphery of these ‘triple expressors’ with 488 nm light resulted in a global recruitment of cytosolic CRY2PHR, fused with a protein of interest, to the plasma membrane (Figure S1B-D). The corresponding linescan clearly shows that the CRY2PHR-mcherry2 intensity peak shifts from the cytosol to the plasma membrane when blue light is switched on, demonstrating membrane recruitment (Figure S1E). Similarly, once blue light was selectively shined on the front or back of the cell, the CRY2PHR-fused protein of interest was locally recruited to the illuminated region of the membrane (Figure S1B and C). This system thus allowed us to spatio-temporally control activities of signaling components on the membrane in a tightly regulated fashion.

### Global Ras activation increased spreading and motility in neutrophils

Using our optogenetic system, we investigated the role of Ras activation on neutrophil morphology in absence of chemoattractant-mediated receptor stimulation. Within a few seconds of turning on the 488 nm light, constitutively activated HRas G12V isoform lacking its C-terminal CAAX motif, HRas G12V ΔCAAX, was recruited uniformly on the cell membrane. Immediately, an increase in F-actin-rich protrusions, marked by LifeAct, was observed and cells started moving rapidly (Figure 1A, Video S1). In a representative kymograph for these cells, narrow regions of LifeAct intensity existed in absence of blue light. However, once light was switched on, LifeAct intensity on the membrane increased substantially with a concomitant increase in cell surface area. In this HRas activated state, cells also demonstrated spreading and contraction periodically (Figure 1B). As illustrated in Figure 1C, there was an overall ∼40% increase in neutrophil surface area. Additionally, HRas activity on the membrane polarized cells and improved their migratory ability (Figure 1A). This led to a 70% and 20% increase in migration speed (Figure 1D-F) and aspect ratio (which serves as a proxy for polarity; Figure 1G), respectively, in the recruited cells. As one control, we looked at cells in the same population which showed no detectable recruitment of HRas G12V ΔCAAX to the membrane, presumably due to low expression of CIBN-CAAX. These cells, even in presence of 488 nm light, did not show any appreciable change in size of F-actin patches, protrusion shape, surface area, migration speed or polarity (Figure S2A-G). These data suggest that HRas-mediated signaling at the membrane causes the improved spreading and migration of neutrophils.

**Figure 1.**
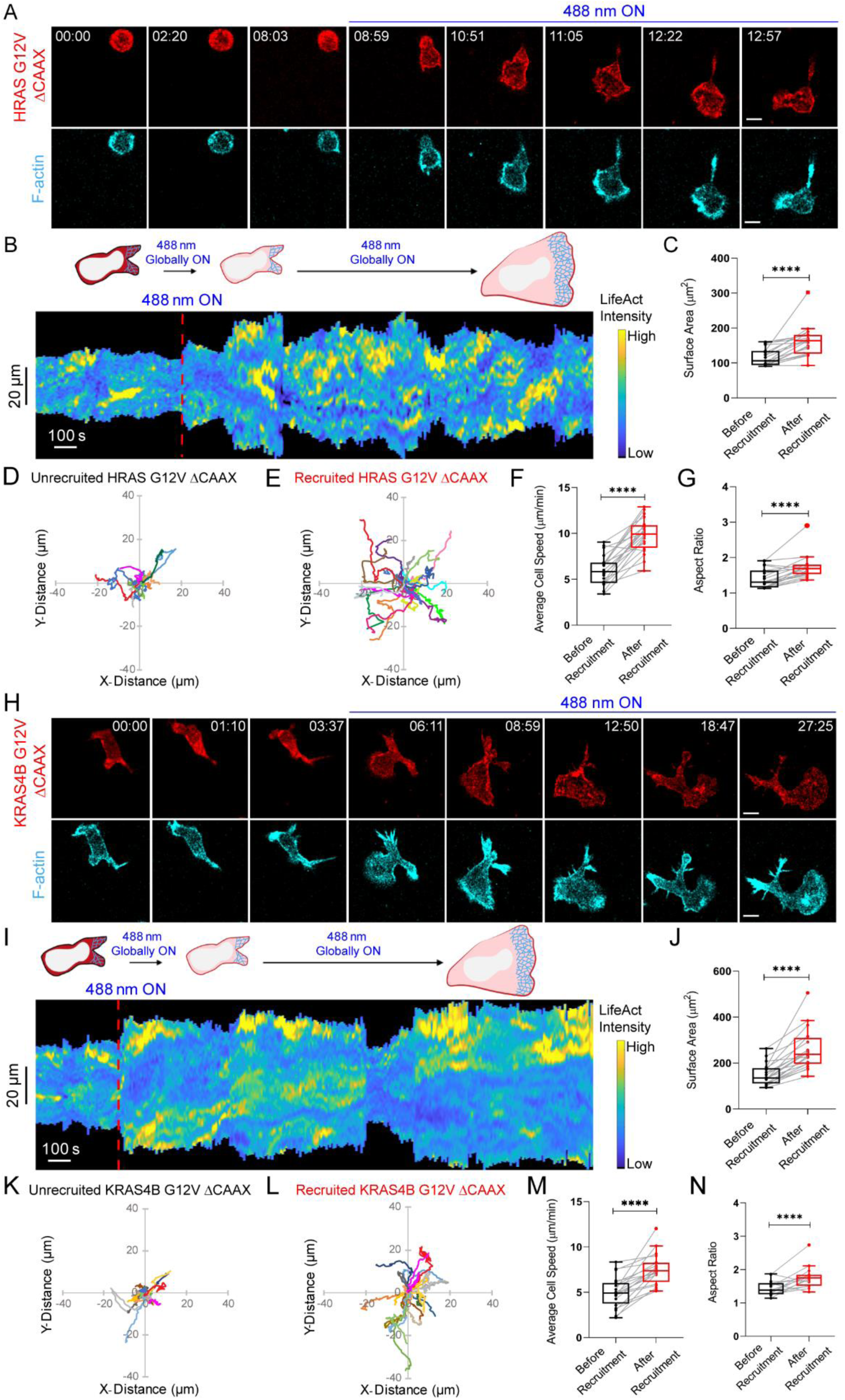
Global recruitment of constitutively active Ras isoforms improves neutrophil polarity and migration. **(A)** Time-lapse confocal images of differentiated HL-60 neutrophil expressing CRY2PHR-mcherry2-HRas G12V ΔCAAX (red; upper panel) and LifeActmiRFP703 (cyan; lower panel), before or after 488 nm laser was switched on globally. Time in min:sec format. Scale bars represent 5 µm. **(B)** Representative kymograph of cortical LifeAct intensity in HRas G12V ΔCAAX-expressing neutrophil before or after 488 nm laser was turned on. A linear color map shows that blue is the lowest LifeAct intensity whereas yellow is the highest. Duration of the kymograph is 30 mins. Cartoon depicts recruitment, F-actin polymerization or cell shape status corresponding to the kymograph. **(C)** Box-and-whisker plot of neutrophil surface area before (black) or after (red) CRY2PHR-mcherry2-HRas G12V ΔCAAX membrane recruitment. n_c_=19 from atleast 3 independent experiments; asterisks indicate significant difference, ****P ≤ 0.0001 (Wilcoxon-Mann-Whitney rank sum test). Centroid tracks of neutrophils (n_c_=19) showing random motility before **(D)** or after **(E)** HRas G12V ΔCAAX membrane recruitment. Each track lasts atleast 5 mins and was reset to same origin. Box-and-whisker plots of neutrophil speed **(F)** and aspect ratio **(G)** before (black) or after (red) HRas G12V ΔCAAX membrane recruitment. n_c_=19 from atleast 3 independent experiments; asterisks indicate significant difference, ****P ≤ 0.0001 (Wilcoxon-Mann-Whitney rank sum test). **(H)** Confocal images of differentiated HL-60 neutrophil expressing CRY2PHR-mcherry2-KRas4B G12V ΔCAAX (red; upper panel) and LifeActmiRFP703 (cyan; lower panel), before or after 488 nm laser was switched on. Time in min:sec format. Scale bars represent 5 µm. **(I)** Representative kymograph of cortical LifeAct intensity in KRas4B G12V ΔCAAX-expressing neutrophil before or after 488 nm laser was turned on globally. A linear color map shows that blue is the lowest LifeAct intensity whereas yellow is the highest. Duration of the kymograph is 35 mins. Cartoon depicts membrane recruitment, F-actin polymerization or cell shape status corresponding to the kymograph. **(J)** Box-and-whisker plot of neutrophil surface area before (black) or after (red) CRY2PHR-mcherry2-KRas4B G12V ΔCAAX membrane recruitment. n_c_=20 from atleast 3 independent experiments; asterisks indicate significant difference, ****P ≤ 0.0001 (Wilcoxon-Mann-Whitney rank sum test). Centroid tracks of neutrophils (n_c_=20) showing random motility before **(K)** or after **(L)** CRY2PHR-mcherry2-KRas4B G12V ΔCAAX membrane recruitment. Each track lasts atleast 5 mins and was reset to same origin. Box-and-whisker plots of neutrophil speed **(M)** and aspect ratio **(N)** before (black) or after (red) KRas4B G12V ΔCAAX membrane recruitment. n_c_=20 from atleast 3 independent experiments; asterisks indicate significant difference, ****P ≤ 0.0001 (Wilcoxon-Mann-Whitney rank sum test).

To test whether these cytoskeletal changes are Ras isoform-specific, we expressed a constitutively active KRas4B G12V ΔCAAX in neutrophils. Once 488 nm laser was switched on and KRas4B membrane recruitment occurred, these cells formed broad F-actin-driven lamellipodium and showed increased motility (Figure 1H, Video S2). A representative kymograph showed that, upon activating KRas4B, LifeAct intensity was tightly coordinated with cell surface area. During the spreading and contracting stages of the cell, LifeAct intensity increased and decreased accordingly (Figure 1I). In non-recruiting cells of this population, narrow regions of LifeAct intensity were present stochastically, and motility was minimal (Figure S2H and I). Overall, KRas4B activation gave rise to 70%, 50%, and 20% increase in cell surface area, migration speed, and polarity, respectively (Figure 1J-N). In the non-recruiting cells, all these migration parameters remained unchanged over time (Figure S2J-N). Altogether, HRas activity was a stronger inducer of motility whereas KRas promoted more spreading.

Although Ras activation-mediated cytoskeletal changes and motility occurred in absence of chemoattractant, we wanted to ensure that these phenotypes were caused solely by recruiting active Ras on the membrane and not basal G-protein coupled receptor signaling^86^. To test this, we inhibited heterotrimeric G-protein activity in neutrophils using a combination of Gα_i_ and Gβγ inhibitors, pertussis toxin (PTX) and gallein, respectively (Figure S3A and B)^18, 47, 83, 87–90^. Dual inhibitor treatment prevented the typical uniform chemoattractant-induced burst of F-actin polymerization all around the cell periphery and subsequent polarized migration (Figure S3B). This indicated that G-protein signaling was completely stalled in neutrophils. However, when KRas4B G12V ΔCAAX was recruited globally in these inhibited cells, they polarized and started migrating with broad fronts as observed previously (Figure S3C and Video S3; Figure 1H and Video S2). Representative kymograph showed that narrow LifeAct patches were converted to broad ones, accompanied with an increase in cell surface area once laser was switched on (Figure S3D). This suggests that spontaneous Ras activity on the membrane, independent of any receptor-mediated G-protein signaling, is sufficient to activate neutrophil motility and polarity, and Ras should be included as an intermediary coupling G-protein activity to downstream signaling (Figure S3A).

Since individual Ras isoforms could promote neutrophil migration, we wondered whether concerted Ras activation by a RasGEF would have a similar or stronger effect. We selected RasGRP4 for our study since it is expressed exclusively in myeloid cells and its dysregulation leads to the onset of diseases such as asthma, mastocytosis, and leukemia^91, 92^. Its involvement in immune cell migration has not been explored. Time-lapse imaging and kymograph analysis showed that, after a few minutes of recruiting RasGRP4 to the membrane, narrow, transient pseudopods at the cell front were converted to wide, sustained lamellipodium resulting in prolonged migration (Figure 2A-C, Video S4). Altogether, RasGRP4 membrane activation resulted in over 86% increase in neutrophil surface area, accompanied with a ∼70% and ∼25% rise in migration speed and polarity, respectively (Figure 2D-H). Neutrophils in the same population with cytosolic, non-recruiting RasGRP4 were not activated (Figure S4). Also, no response was observed when the CRY2PHR component, without being fused to either activated Ras or RasGEF, was recruited to the cell membrane (Figure S5, Video S5).

**Figure 2.**
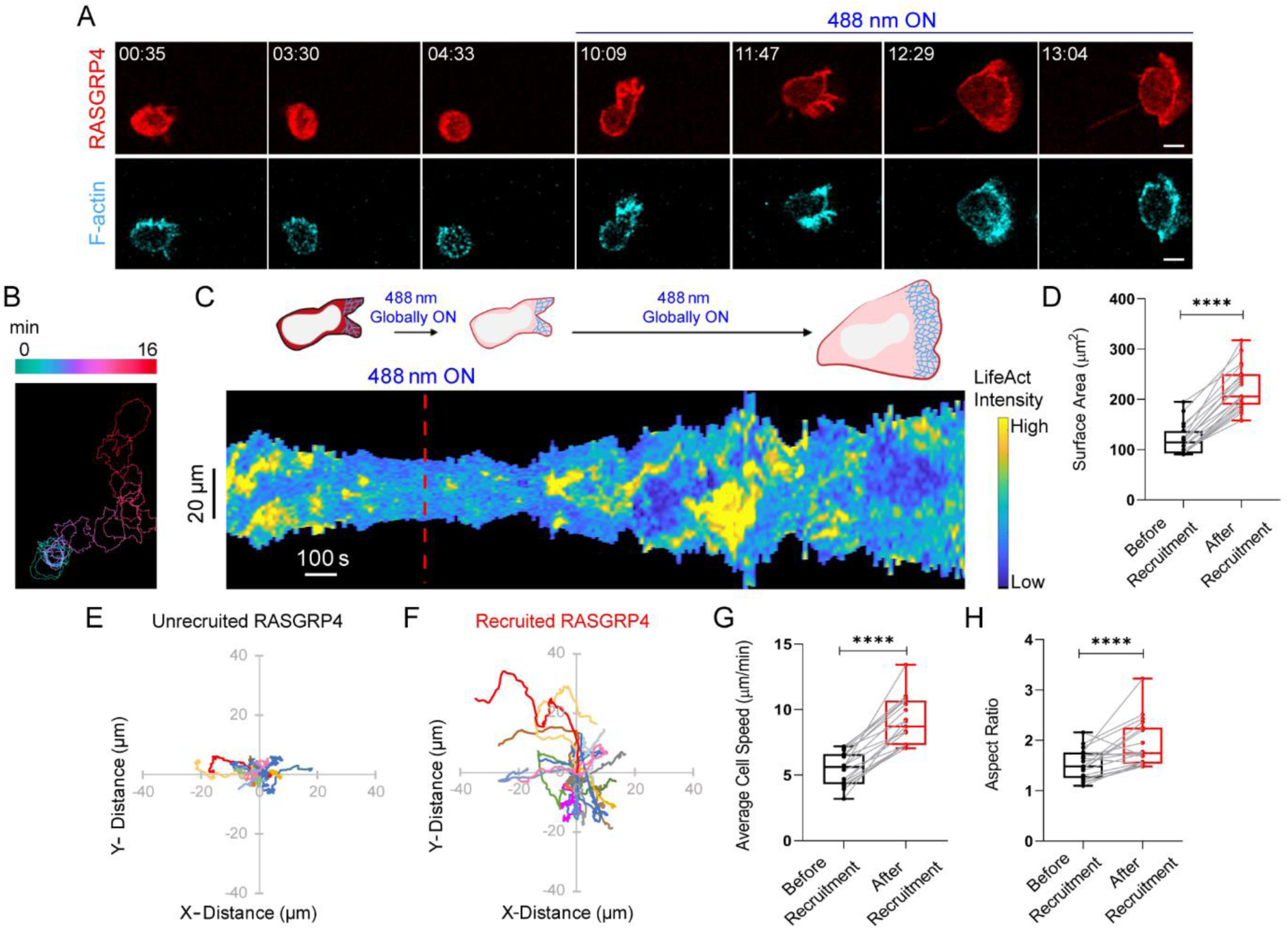
RasGEF activation on neutrophil membrane induces polarity and migration. **(A)** Time-lapse confocal images of differentiated HL-60 neutrophil expressing CRY2PHR-mcherry2-RasGRP4 (red; upper panel) and LifeActmiRFP703 (cyan; lower panel), before or after 488 nm laser was switched on globally. Time in min:sec format. Scale bars represent 5 µm. **(B)** Color-coded (at 1 min intervals) outlines of the cell shown in (A). **(C)** Representative kymograph of cortical LifeAct intensity in RasGRP4-expressing neutrophil before or after 488 nm laser was turned on. A linear color map shows that blue is the lowest LifeAct intensity whereas yellow is the highest. Duration of the kymograph is 24 mins. Cartoon depicts membrane recruitment, actin polymerization or cell shape status corresponding to the kymograph. **(D)** Box-and-whisker plot of cell surface area before (black) or after (red) RasGRP4 membrane recruitment. n_c_=19 from atleast 3 independent experiments; asterisks indicate significant difference, ****P ≤ 0.0001 (Wilcoxon-Mann-Whitney rank sum test). Centroid tracks of neutrophils (n_c_=19) showing random motility before **(E)** or after **(F)** RasGRP4 membrane recruitment. Each track lasts atleast 5 mins and was reset to same origin. Box-and-whisker plots of neutrophil speed **(G)** and aspect ratio **(H)** before (black) or after (red) RasGRP4 recruitment. n_c_=19 from atleast 3 independent experiments; asterisks indicate significant difference, ****P ≤ 0.0001 (Wilcoxon-Mann-Whitney rank sum test).

Next, we aimed to provide a mechanistic insight to actin assembly guiding Ras-driven lamellipodium formation in neutrophils. We assessed the effects of blocking the actin nucleator, Arp2/3 complex, whose activation by Scar/WAVE causes lamellipodia and phagocytic cup formation in immune cells^10, 93, 94^. After treating recruitable RasGRP4-expressing cells with Arp2/3 complex inhibitor, CK-666, they rounded up, actin polymerization on the cortex vanished, and migration was abrogated. Despite recruiting RasGRP4 on the membrane, cells did not display any F-actin-rich protrusions, cell spreading, motility, or polarity (Figure S6). Altogether, our results showed that Ras activity on the cell membrane leads to Arp2/3-mediated F-actin polymerization causing spreading and migration in mammalian cells.

### Local activation or inhibition of Ras reverses pre-existing polarity

Activating Ras over the entire cell periphery led to formation of actin-rich lamellipodia, increased polarity, and persistent migration, but could local Ras activation at the cell rear generate new protrusions and re-organize polarity? To address this question, we recruited RasGRP4 to a quiescent back region in migrating neutrophils by intermittently shining 488 nm light near it, as indicated by dashed white box (Figure 3A, Video S6). Immediately after RasGRP4 localized to the back, the cell front contracted, ruffles started diminishing, and the cell slowed down. Simultaneously, several finger-like F-actin-rich structures appeared at the site of recruitment, which gradually broadened into sustained protrusions. This forced the cell to move in the other direction by rearranging its front-rear axis (Figure 3A and E). This phenomenon was analyzed in the angular histogram which showed that the probability of new protrusion generation was highest at or near the site of RasGRP4 recruitment (Figure 3B).

**Figure 3.**
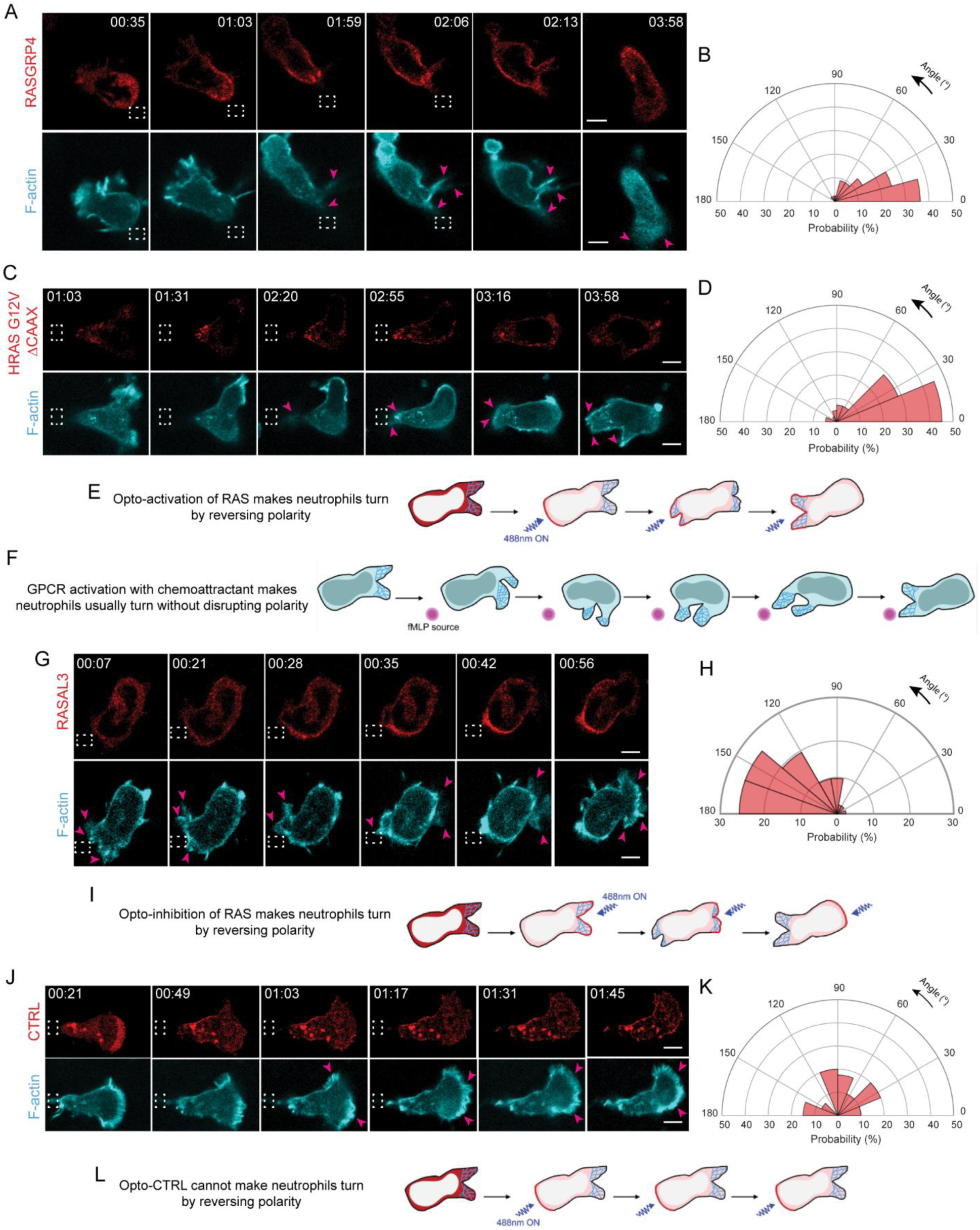
Localized Ras activation or inhibition rearranges front-rear polarity in neutrophils. **(A)** Time-lapse confocal images of differentiated HL-60 neutrophil expressing CRY2PHR-mcherry2-RasGRP4 (red; upper panel) and LifeActmiRFP703 (cyan; lower panel). RasGRP4 is recruited exclusively to the back of the cell by applying 488 nm laser near it, as shown by the dashed white box. Pink arrows highlight RasGRP4-induced new protrusion formation. Time in min:sec format. Scale bars represent 5 µm. **(B)** Polar histogram demonstrates higher probability of fresh protrusion formation near recruitment area; n_c_=16 and n_p_=44. **(C)** Time-lapse confocal images of differentiated HL-60 neutrophil expressing CRY2PHR-mcherry2-HRas G12V ΔCAAX (red; upper panel) and LifeActmiRFP703 (cyan; lower panel). HRas G12V ΔCAAX is recruited exclusively to the cell rear by shining 488 nm laser near it, as indicated by the dashed white box. Pink arrows highlight HRas G12V ΔCAAX-induced new protrusion formation. Time in min:sec format. Scale bars represent 5 µm. **(D)** Polar histogram demonstrates higher probability of fresh protrusion formation near HRas G12V ΔCAAX recruitment area; n_c_=23 and n_p_=42. **(E)** Cartoon illustrating that opto-activation of Ras at the cell back, by RasGRP4 or HRas G12V ΔCAAX recruitment, makes neutrophils turn by reversing polarity. **(F)** In response to a chemoattractant gradient at the back, cell makes a ‘U-turn’ to reorient itself towards the fMLP source without breaking its pre-existing polarity. **(G)** Time-lapse confocal images of differentiated HL-60 neutrophil expressing CRY2PHR-mcherry2-RASAL3 (red; upper panel) and LifeActmiRFP703 (cyan; lower panel). RASAL3 was recruited exclusively to front protrusions (marked by LifeAct) by applying 488 nm laser near it, as shown by the dashed white box. Pink arrows highlight disappearance of protrusions at the RASAL3-recruitment site and appearance of new ones away from it. Time in min:sec format. Scale bars represent 5 µm. **(H)** Polar histogram shows higher probability of fresh protrusion formation away from the recruitment area; n_c_=21 and n_p_=41. **(I)** Cartoon illustrating that opto-inhibition of Ras by RASAL3 recruitment at the front, makes neutrophils turn by reversing polarity. **(J)** Time-lapse confocal images of differentiated HL-60 neutrophil expressing CRY2PHR-mcherry2-CTRL (control; red; upper panel) and LifeActmiRFP703 (cyan; lower panel). CTRL is recruited exclusively to the rear of the cell by applying 488 nm laser near it, as shown by the dashed white box. Pink arrows indicate new protrusions forming at the opposite end. Time in min:sec format. Scale bars represent 5 µm. **(K)** Polar histogram demonstrates CTRL recruitment does not bias new protrusion formation; n_c_=15 and n_p_=40. **(L)** Cartoon illustrating opto-CTRL (control) recruitment at the back cannot turn neutrophils by reversing polarity.

Since RasGRP4 may activate a number of Ras proteins, we next asked whether local activation of a specific Ras isoform could elicit a similar response at the cell cortex. To examine this, we transiently recruited HRas G12V ΔCAAX to the back of migrating neutrophils. This arrested protrusion formation at the cell front, and brought about a concomitant increase in F-actin polymerization at the site of recruitment. As a result, the cell changed its direction of migration, reversing its pre-existing polarity (Figure 3C and E, Video S7). The corresponding angular histogram demonstrated that new protrusion formation could be biased towards the activated HRas G12V recruitment site (Figure 3D). When stimulated by chemoattractant from behind, neutrophils usually maintain their pre-existing polarity and make a U-turn towards the new source; they can only switch polarity if the gradient is extraordinarily steep and perfectly positioned (Figure 3F)^80^. Apparently, local activation of Ras is equivalent to a very steep chemoattractant gradient.

We next examined the effects of inhibiting Ras activity on cytoskeletal assembly and protrusions, which might be expected to stop immune cell migration. We studied a RasGAP, RASAL3, which was recently characterized for its immunomodulatory functions in neutrophils. It is possible that these effects on immune response were due to inhibition of migration, but this was not assessed^95^. For our experiment, we transiently recruited RASAL3 to the leading front by shining light periodically near the existing protrusions, as shown by the dashed white box (Figure 3G, Video S8). Upon doing so, protrusions were restricted and eventually collapsed in on themselves. Thus, a new back formed at the recruitment site, whereas a new F-actin rich front emerged on the other end of the cell. The angular histogram analysis agreed with our observation that all new protrusions formed away from the site of RASAL3 recruitment (Figure 3H). A similar observation was made when RASAL3 was locally recruited to the membrane of activated macrophages (Figure S7A, Video S9). Therefore, through its RasGAP function, RASAL3 managed to change the direction of the neutrophil by reversing pre-existing polarity (Figure 3G and I).

We next enquired whether dynamically recruiting RASAL3 over the entire cell periphery would shut down all protrusive activity in immune cells. For this experiment, we selected a chemoattractant-activated macrophage possessing F-actin driven protrusions around its periphery, as shown by pink arrows (Figure S7A, Video S9). Once we applied 488 nm light continuously all around the cell perimeter, it caused RASAL3 to recruit globally over the periphery. This resulted in cell shrinkage and protrusions vanished instantaneously (Figure S7A and C, Video S9). We could reproduce a similar phenotype in a non-polarized neutrophil having cellular protrusions around its perimeter (Figure S7B, Video S10). Upon shining 488 nm laser continuously around its periphery, RASAL3 was globally recruited causing protrusions to disappear completely. This was accompanied by a concomitant decrease in cell size (Figure S7B and C, Video S10).

Locally recruiting CRY2PHR alone did not bias new protrusion formation nor affect front-rear axis of the cell (Figure 3J-L, Video S11). This suggested that reversing polarity, either to create new cellular protrusions at the back or inhibit existing protrusions at the front, is due to localized activation or inhibition of Ras, and not due to cryptochrome recruitment or light irradiation. Taken together, all these observations suggest that transient Ras activation is sufficient and necessary to orchestrate cytoskeletal arrangement and generate protrusions.

### mTorC2 is more important than PI3K for Ras activation of polarity and migration

Our next aim was to delineate Ras downstream signaling in neutrophils. We first examined role of the PI3K pathway using a combination of optogenetic recruitment and pharmacological inhibition approaches. Although PI3Kγ is a direct effector of Ras and an important regulator of polarization and migration in many cells, the role of Ras-activated PI3K in neutrophil motility is not known^34, 42, 43, 47^. For our experiments, we selected the PI3Kγ inhibitor, AS605240, which depleted PIP3 as indicated by the signal loss of its biosensor, PH-Akt, on the neutrophil membrane (Figure S8)^96, 97^. Upon inhibitor treatment, RasGRP4-expressing cells lost polarity and migration ability (Figure 4A and B, Video S12). Kymographs showed a few spontaneous, narrow patches of LifeAct in PI3K-inhibited cells (Figure 4C). When RasGRP4 was recruited to the membrane, the PI3K-inhibited cells regained polarity, developed F-actin rich fronts, and migrated (Figure 4B, Video S12). Upon recruitment, LifeAct patches broadened and increased in number along with widening of surface area (Figure 4C). In total, Ras activation caused an increase of 50%, 38%, and 64% in surface area, polarity, and migration speed, respectively (Figure 4D-H). In non-recruiting cells where RasGRP4 recruitment was undetectable, there was no recovery from PI3K inhibition (Figure S9). We also independently checked chemoattractant-induced phosphorylation of Akt at Thr308 and Ser473 positions as an indirect readout of Ras activation in PI3K-inhibited cell population. As expected, basal phosphorylation at both crucial residues almost completely disappeared with inhibitor treatment. Upon activation, phosphorylation on Akt recovered close to untreated levels, indicating Ras activates downstream Akt in absence of PI3K activity (Figure 4I). Overall, our results suggest that while PI3K activity may be important for spontaneous protrusion formation and motility, it is not required for chemoattractant- or optically-triggered protrusive activity.

**Figure 4.**
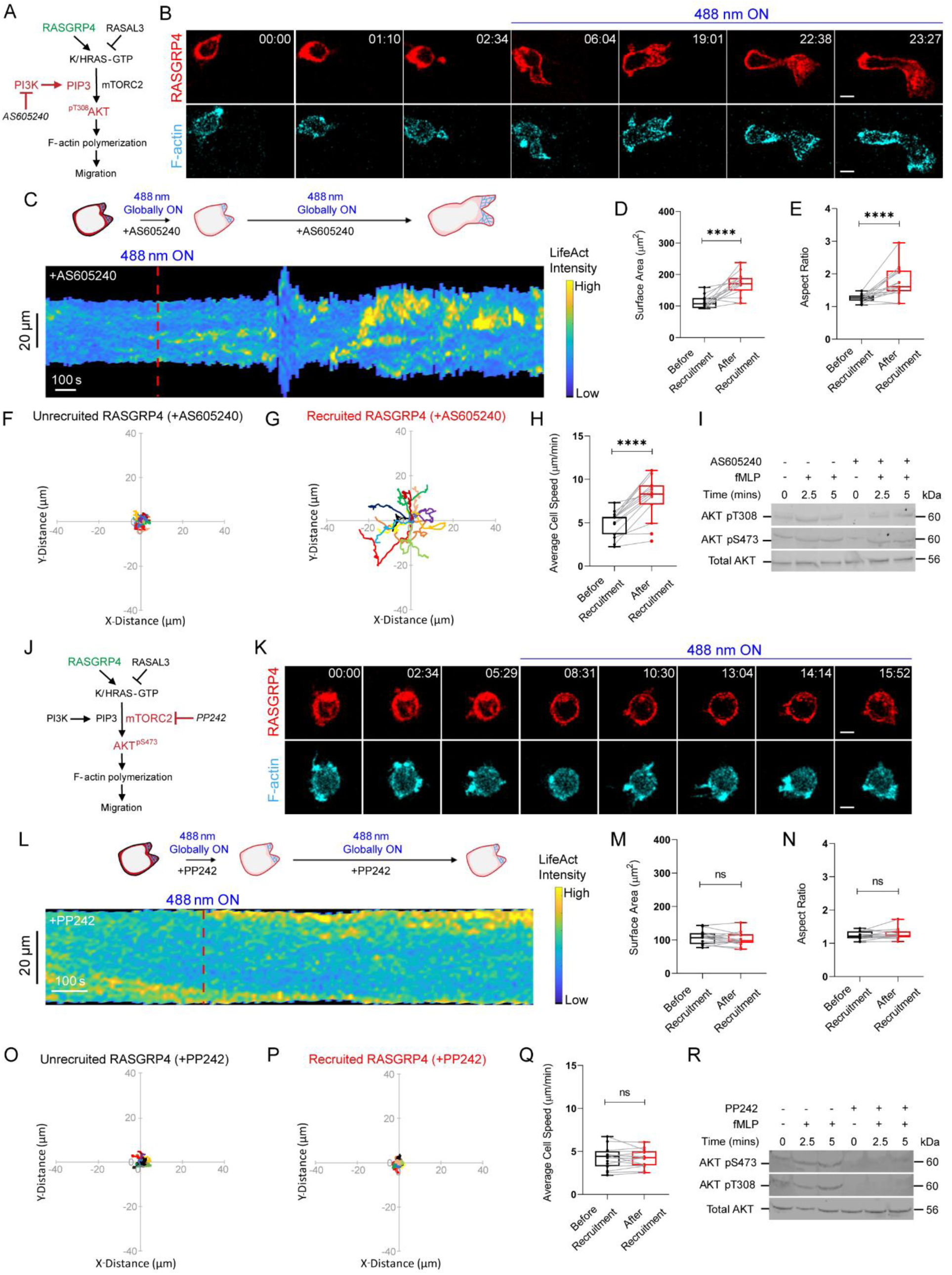
mTorC2 is more important than PI3K for Ras activation of polarity and migration. **(A)** Proposed model highlighting Ras activation, through RasGRP4 recruitment, promotes F-actin polymerization and migration by Akt phosphorylation in absence of PI3K. **(B)** Time-lapse confocal images of AS605240-treated HL-60 neutrophil expressing CRY2PHR-mcherry2-RasGRP4 (red; upper panel) and LifeActmiRFP703 (cyan; lower panel), before or after 488 nm laser was switched on globally. Time in min:sec format. Scale bars represent 5 µm. **(C)** Representative kymograph of cortical LifeAct intensity in AS605240-treated RasGRP4-expressing neutrophil before or after 488 nm laser was turned on. A linear color map shows that blue is the lowest LifeAct intensity whereas yellow is the highest. Duration of the kymograph is 24 mins. Cartoon depicts recruitment, F-actin polymerization or cell shape status corresponding to the kymograph. Box-and-whisker plot of cell surface area **(D)** or aspect ratio **(E)** before (black) or after (red) CRY2PHR-mcherry2-RasGRP4 membrane recruitment in AS605240-treated neutrophils. n_c_=15 from atleast 3 independent experiments; asterisks indicate significant difference, ****P ≤ 0.0001 (Wilcoxon-Mann-Whitney rank sum test). Centroid tracks of AS605240-treated neutrophils (n_c_=15) showing random motility before **(F)** or after **(G)** RasGRP4 membrane recruitment. Each track lasts atleast 5 mins and was reset to same origin. **(H)** Box-and-whisker plot of AS605240-treated neutrophil speed before (black) or after (red) RasGRP4 membrane recruitment. n_c_=15 from atleast 3 independent experiments; asterisks indicate significant difference, ****P ≤ 0.0001 (Wilcoxon-Mann-Whitney rank sum test). **(I)** Representative western blot demonstrating fMLP-stimulated phospho-Akt levels (Thr308 and Ser473; ∼60kDa) in untreated or AS605240-treated whole cell lysates. Total Akt (∼56kDa) was used as loading control. **(J)** Proposed model highlighting Ras activation, through RasGRP4 recruitment, cannot promote F-actin polymerization and migration by Akt phosphorylation in absence of mTorC2. **(K)** Time-lapse confocal images of PP242-treated HL-60 neutrophil expressing CRY2PHR-mcherry2-RasGRP4 (red; upper panel) and LifeActmiRFP703 (cyan; lower panel), before or after 488 nm laser was switched on globally. Time in min:sec format. Scale bars represent 5 µm. **(L)** Representative kymograph of cortical LifeAct intensity in PP242-treated RasGRP4-expressing neutrophil before or after 488 nm laser was turned on. A linear color map shows that blue is the lowest LifeAct intensity whereas yellow is the highest. Duration of the kymograph is 17 mins. Cartoon depicts recruitment, F-actin polymerization or cell shape status corresponding to the kymograph. Box-and-whisker plot of cell surface area **(M)** or aspect ratio **(N)** before (black) or after (red) CRY2PHR-mcherry2-RasGRP4 membrane recruitment in PP242-treated cells. n_c_=15 from atleast 3 independent experiments; ns denotes P>0.05 (Wilcoxon-Mann-Whitney rank sum test). Centroid tracks of PP242-treated neutrophils (n_c_=15) showing random motility before **(O)** or after **(P)** RasGRP4 membrane recruitment. Each track lasts atleast 5 mins and was reset to same origin. **(Q)** Box-and-whisker plot of PP242-treated neutrophil speed before (black) or after (red) RasGRP4 membrane recruitment. n_c_=15 from atleast 3 independent experiments; ns denotes P>0.05 (Wilcoxon-Mann-Whitney rank sum test). **(I)** Representative western blot demonstrating fMLP-stimulated phospho-Akt levels (Thr308 and Ser473; ∼60kDa) in untreated or PP242-treated whole cell lysates. Total Akt (∼56kDa) was used as loading control.

To test whether Ras regulates mTorC2 signaling in neutrophil polarization and migration, we treated RasGRP4-expressing cells with mTor inhibitor, PP242 (Figure 4J)^98^. The cells rounded up and ceased motility quickly (Figure 4K, Video S13). Upon recruiting RasGRP4, we did not see any recovery in cell shape, F-actin protrusions, polarity, surface area, or migration speed (Figure 4K-Q). The mTorC1 inhibitor, rapamycin, did not affect neutrophil shape or size, indicating that PP242-induced cytoskeletal changes were due to mTorC2 inhibition, and not mTorC1 (Figure S10). Additionally, Akt phosphorylation at both Thr308 and Ser473 failed to recover with chemoattractant-mediated activation in PP242-treated population (Figure 4R), suggesting Ras induces Akt activation via mTorC2 complex.

### Ras-mTorC2 effector, Akt, activates cellular protrusions and migration

We next asked the extent to which Akt activity directly affects cellular protrusive activity. There are two Akt isoforms in neutrophils, Akt1 and Akt2. Despite several studies in these cells, there is no clear consensus whether effect of each on motility is positive or negative^45, 46^. To resolve this dilemma, we first selected Akt1 for optical recruitment studies (Figure 5A and B). In our experiment, we transiently recruited Akt1 to the membrane at the quiescent back of a moving cell. As a result, the front protrusions started retracting, and a broad new protrusion soon emerged at the site of recruitment (Figure 5C and E, Video S14). This phenomenon was observed consistently across the population, as highlighted in the angular histogram (Figure 5D). To test if the polarity reversing ability of Akt1 was due to its kinase activity, we similarly recruited a Akt kinase-dead mutant, Akt1_T308A_, to the rear of motile neutrophils (Figure 5F and G)^99^. Akt1_T308A_ mutant could not induce fresh protrusion formation at the site of recruitment, and polarity remained unaffected (Figure 5H and J, Video S15). Angular histogram analysis validated our observations that recruitment of inactive Akt1_T308A_ did not bias new protrusion formation at the neutrophil back (Figure 5I).

**Figure 5.**
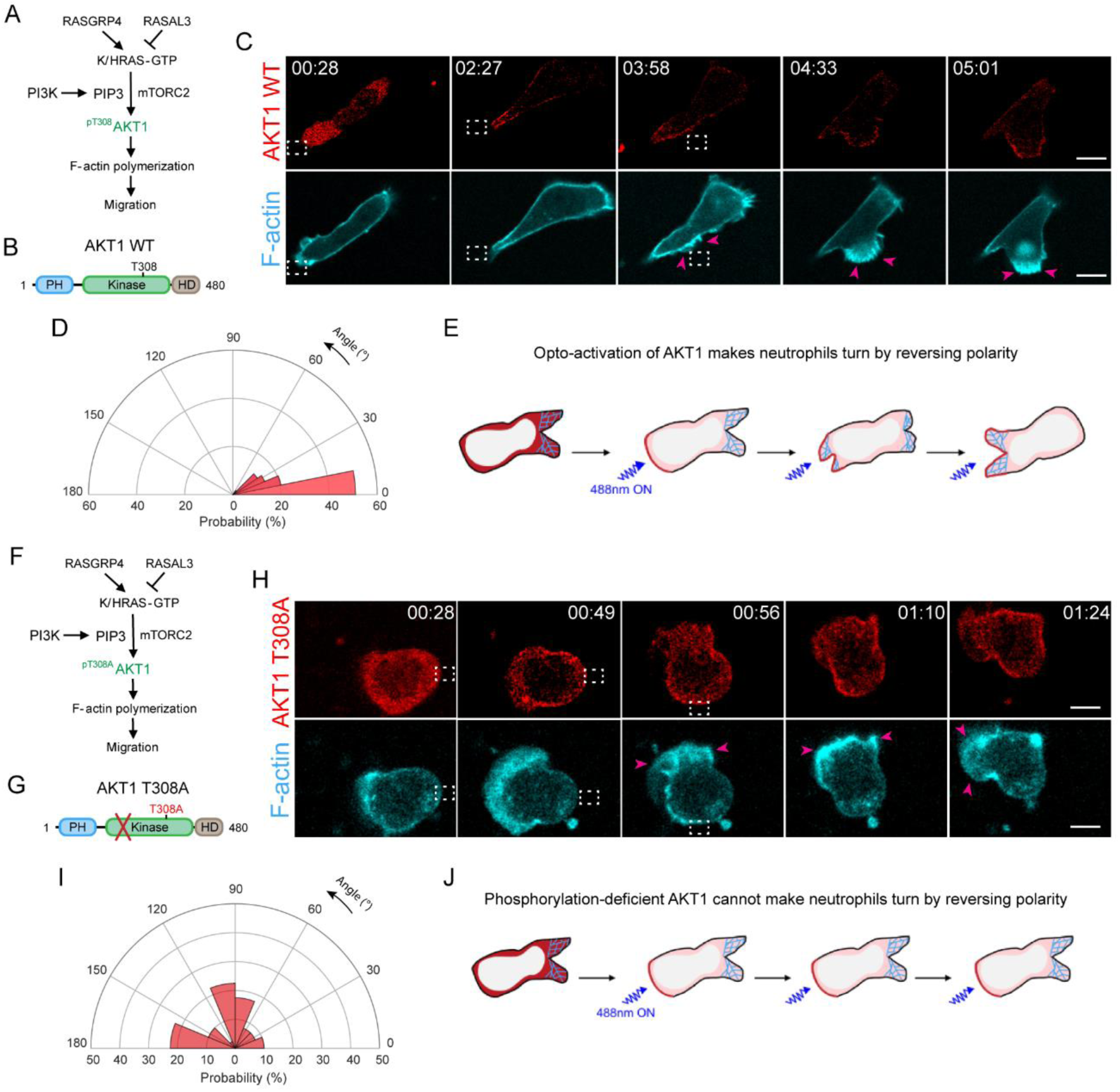
Localized Akt1 activation at the rear of neutrophils reverses front-back polarity. **(A)** Proposed model highlighting Akt1 opto-activation promotes F-actin polymerization and migration by phosphorylation at T308 position. **(B)** Cartoon of Akt1 WT protein sequence with PH, kinase, and hypervariable domains. Phosphorylation at T308 position is critical for Akt1 activity. **(C)** Time-lapse confocal images of differentiated HL-60 neutrophil expressing CRY2PHR-mcherry2-Akt1 WT (red; upper panel) and LifeActmiRFP703 (cyan; lower panel). Akt1 WT is recruited exclusively to the back of the cell by applying 488 nm laser near it, as shown by the dashed white box. Pink arrows highlight Akt1 WT-induced new protrusion formation. Time in min:sec format. Scale bars represent 5 µm. **(D)** Polar histogram demonstrates higher probability of fresh protrusion formation near recruitment area; n_c_=19 and n_p_=49. **(E)** Cartoon illustrating that opto-activation of Akt1 at the cell back makes neutrophils turn by reversing polarity. **(F)** Proposed model highlighting phospho-dead Akt1_T308A_ optical recruitment cannot promote F-actin polymerization and migration. **(G)** Cartoon of Akt1_T308A_ protein sequence with PH, kinase, and hypervariable domains. Due to T308A mutation, phosphorylation and kinase activity are absent. **(H)** Time-lapse confocal images of differentiated HL-60 neutrophil expressing CRY2PHR-mcherry2-Akt1_T308A_ (red; upper panel) and LifeActmiRFP703 (cyan; lower panel). Akt1_T308A_ is recruited exclusively to the cell rear by shining 488 nm laser near it, as indicated by the dashed white box. Pink arrows indicate new protrusions forming at the opposite end. Time in min:sec format. Scale bars represent 5 µm. **(I)** Polar histogram demonstrates Akt1_T308A_ recruitment does not bias new protrusion formation; n_c_=15 and n_p_=40. **(J)** Cartoon illustrating that optically recruiting phosphorylation-deficient, kinase-dead Akt1_T308A_ at the cell back has no effect on neutrophil polarity.

To test if the protrusion-promoting effects of Akt are isoform-specific, we recruited Akt2 over the entire cell perimeter. After a few minutes of recruitment, we noticed non-polarized, migration-incompetent cells developing more F-actin at the front, getting elongated, and migrating efficiently with broad lamellipodium (Figure S11A, Video S16). A representative kymograph showed a gradual increase in LifeAct intensity along with widening of surface area (Figure S11B). Akt2 activation brought about 1.83-, 1.47-, and 1.21-fold improvement in surface area, migration speed, and aspect ratio, respectively (Figure S11C-G). No such phenotypic change was observed when the CRY2PHR component, without being fused to Akt2, was recruited to the cell membrane (Figure S5, Video S5).

We next evaluated whether these cytoskeletal effects of Akt are conserved across cell types and species. To test this, we optically recruited PKBA (Akt1 homolog) globally on the *Dictyostelium* membrane^100, 101^. This resulted in rounded, quiescent cells to polarize and migrate efficiently (Figure S12A, Video S17). This phenomenon did not occur when a control SsrA binding partner (SsbB) lacking PKBA was globally recruited to the membrane, suggesting activation of PKBA was responsible, and not light irradiation or SspB recruitment (Figure S12B, Video S18). Thus, both local and global illumination experiments proved that Akt isoforms positively regulate F-actin polymerization, protrusion generation, and migration in an evolutionary conserved manner.

### Akt function does not require PI3K activation

It is well established that PI3K phosphorylates and activates Akt kinase to promote proliferation, survival, and metabolism, in response to extracellular signals^102^. We checked if that holds true in neutrophil polarization and migration. We treated Akt1-expressing neutrophils with PI3K inhibitor, AS605240, which removed polarity, depleted PIP3, and stopped motility completely (Figure 6A-C, Figure S8, Video S19). Interestingly, upon Akt1 activation on the membrane, cells slowly re-polarized, formed sustained lamellipodium, and migrated (Figure 6C, Video S19). Kymographs showed slightly delayed increases in F-actin polymerization and cell spreading after switching on the light (Figure 6D). Box and whisker plots showed 1.70-, 1.54-, and 1.47-fold increment in cell area, speed, and polarity, respectively (Figure 6E-I). Non-recruiting cells in the same population in which Akt1 was not translocated were unable to polarize, spread, or move (Figure S13). PI3K-inhibited neutrophils could be activated only when detectable Akt1 accumulated on the membrane. We next checked whether kinase activity of Akt1 was responsible for these cytoskeletal changes. For this, we utilized the kinase-dead mutant, Akt1_T308A_, for global recruitment (Figure 6J and K). When we turned on the blue laser on PI3K-inhibited cells, inactive Akt1 recruitment could not rescue actin polymerization, cell spread, motility or polarity (Figure 6L, Video S20). Kymograph, bar plot and cell track analyses confirmed our observations (Figure 6M-R). Altogether, these data confirmed a novel mechanism of Akt phosphorylation and activation independent of PI3K activity in neutrophils.

**Figure 6.**
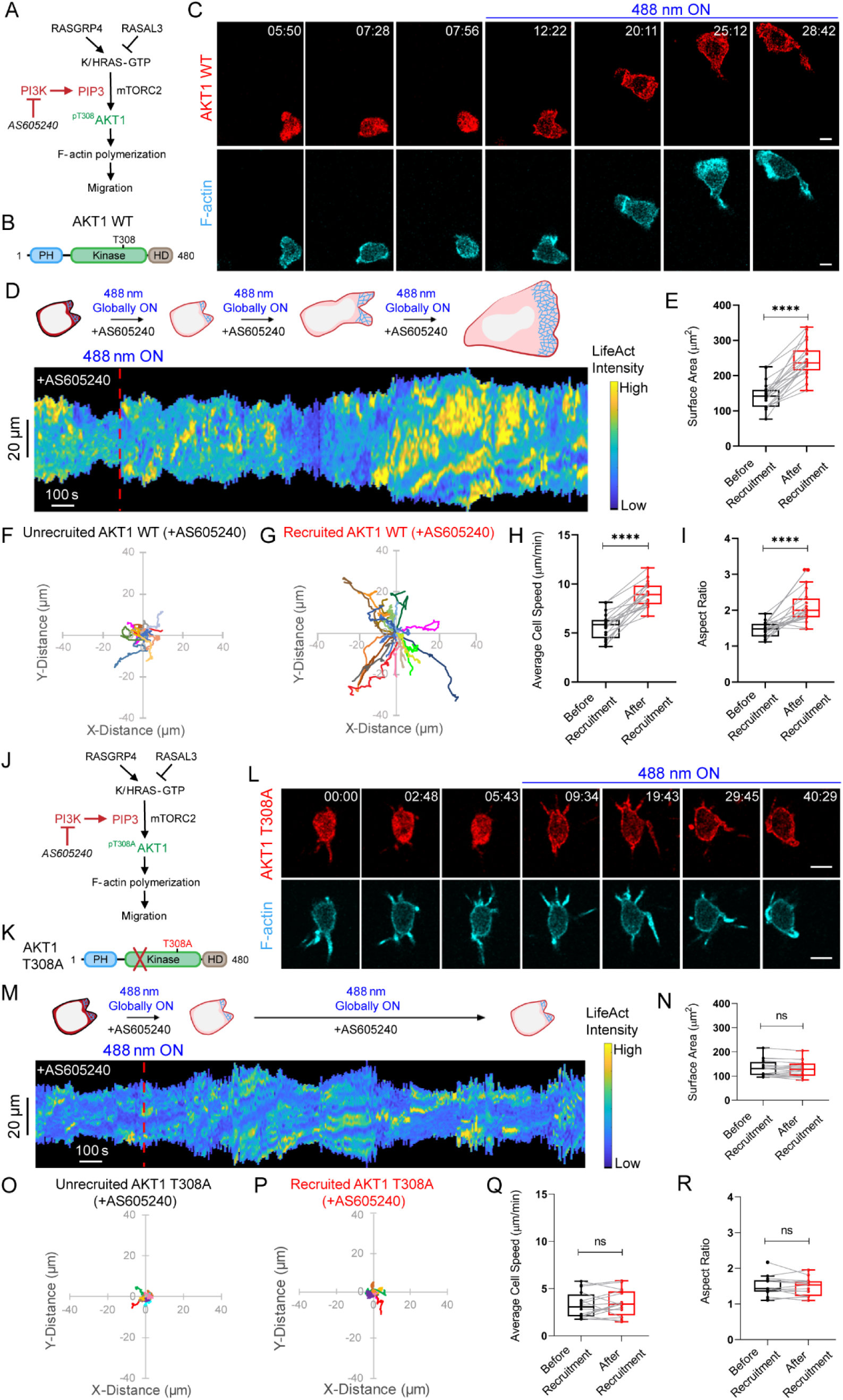
Akt function does not require PI3K activation. **(A)** Proposed model highlighting Akt1 activation promotes F-actin polymerization and migration in absence of PI3K activity. **(B)** Cartoon of Akt1 WT protein sequence with PH, kinase, and hypervariable domains. Phosphorylation at T308 position is critical for Akt1 activity. **(C)** Time-lapse confocal images of AS605240-treated HL-60 neutrophil expressing CRY2PHR-mcherry2-Akt1 WT (red; upper panel) and LifeActmiRFP703 (cyan; lower panel), before or after 488 nm laser was switched on globally. Time in min:sec format. Scale bars represent 5 µm. **(D)** Representative kymograph of cortical LifeAct intensity in AS605240-treated Akt1 WT-expressing neutrophil before or after 488 nm laser was turned on. A linear color map shows that blue is the lowest LifeAct intensity whereas yellow is the highest. Duration of the kymograph is 40 mins. Cartoon depicts recruitment, F-actin polymerization or cell shape status corresponding to the kymograph. **(E)** Box-and-whisker plot of cell surface area before (black) or after (red) Akt1 WT membrane recruitment. n_c_=20 from atleast 3 independent experiments; asterisks indicate significant difference, ****P ≤ 0.0001 (Wilcoxon-Mann-Whitney rank sum test). Centroid tracks of neutrophils (n_c_=20) showing random motility before **(F)** or after **(G)** Akt1 WT membrane recruitment. Each track lasts atleast 5 mins and was reset to same origin. Box-and-whisker plots of cell speed **(H)** and aspect ratio **(I)** before (black) or after (red) Akt1 WT recruitment. n_c_=20 from atleast 3 independent experiments; asterisks indicate significant difference, ****P ≤ 0.0001 (Wilcoxon-Mann-Whitney rank sum test). **(J)** Proposed model highlighting phospho-dead Akt1_T308A_ optical recruitment cannot promote F-actin polymerization and migration in absence of PI3K activity. **(K)** Cartoon of Akt1_T308A_ protein sequence with PH, kinase, and hypervariable domains. Due to T308A mutation, phosphorylation and kinase activity are absent. **(L)** Time-lapse confocal images of AS605240-treated HL-60 neutrophil expressing CRY2PHR-mcherry2-Akt1_T308A_ (red; upper panel) and LifeActmiRFP703 (cyan; lower panel), before or after 488 nm laser was switched on globally. Time in min:sec format. Scale bars represent 5 µm. **(M)** Representative kymograph of cortical LifeAct intensity in AS605240-treated Akt1_T308A_-expressing neutrophil before or after 488 nm laser was turned on. A linear color map shows that blue is the lowest LifeAct intensity whereas yellow is the highest. Duration of the kymograph is 46 mins. Cartoon depicts recruitment, F-actin polymerization or cell shape status corresponding to the kymograph. **(N)** Box-and-whisker plot of cell surface area before (black) or after (red) Akt1_T308A_ membrane recruitment. n_c_=16 from atleast 3 independent experiments; ns denotes P>0.05 (Wilcoxon-Mann-Whitney rank sum test). Centroid tracks of neutrophils (n_c_=16) showing random motility before **(O)** or after **(P)** Akt1_T308A_ membrane recruitment. Each track lasts atleast 5 mins and was reset to same origin. Box-and-whisker plots of cell speed **(Q)** and aspect ratio **(R)** before (black) or after (red) Akt1_T308A_ recruitment. n_c_=16 from atleast 3 independent experiments; ns denotes P>0.05 (Wilcoxon-Mann-Whitney rank sum test).

## Discussion

Our studies show that local Ras-mediated signaling cascades on the plasma membrane directly regulate immune cell polarity and migration. Our optogenetic approaches rule out secondary effects of long-term adaptation and assess effects of localized activations. In the absence of chemoattractant-mediated receptor stimulation, a global increase in Ras activity promoted F-actin-driven lamellipodium formation, stable polarization, and persistent motility in neutrophils. Moreover, strategic activation of Ras alone at inactive, back regions of the cell membrane could generate fresh protrusions thereby reversing pre-existing polarity. These effects can be brought about by local G-protein coupled receptor activation through carefully positioned, steep chemoattractant gradients, but our optogenetic approach showed that localized activity of a single component could have the same outcome. Moreover, our methods allowed us to locally inhibit Ras activity at the front, which extinguished existing protrusions and created a new cell rear, which would not be possible with a chemoattractant. The Ras-mediated cytoskeletal effects worked primarily through mTorC2 complex. Consistently, spontaneously activating the Ras/mTorC2 effector, Akt, could reverse polarity and induce actin polymerization-driven protrusions and migration even in absence of PI3Kγ signaling. Thus, direct activation of the major components of canonical growth control pathways can directly alter cytoskeletal events and immune cell migration (Figure 7).

**Figure 7.**
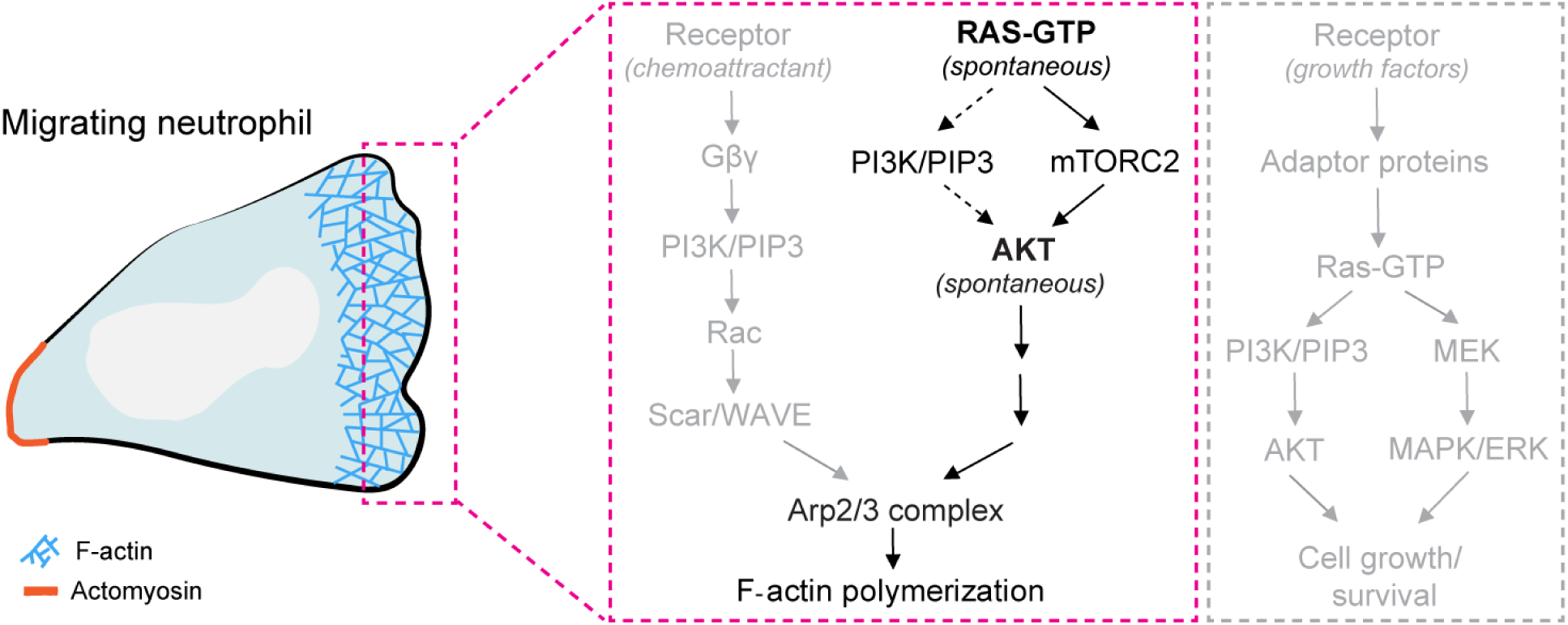
Proposed model of spontaneous Ras-Akt activation mediated actin polymerization and migration. The prevalent model in the field of immune cell migration shows that chemoattractant-mediated G-protein coupled receptor stimulation activates PI3Kγ and PIP3 accumulation which leads to downstream Rac activation. Consequently, Rac triggers it downstream effector, Scar/WAVE complex which initiates Arp2/3 complex-mediated actin polymerization and migration (as shown in grey). In our model (highlighted in black), we propose that local spontaneous Ras activity works predominantly through mTorC2 and Akt to trigger Arp2/3-based F-actin polymerization, lamellipodia formation, and directed migration in neutrophils. Local spontaneous Akt activation also promotes actin polymerization and lamellipodia-driven migration. Ras or Akt activation mediated cytoskeletal changes can occur in apparent absence of PI3K activity, suggesting it is not singularly essential for Ras-mediated polarity and migration (shown in dashed arrows). Akt activation may trigger Rac and its downstream effectors, but was not tested in our study. We report for the first time a direct role of receptor-independent, spontaneous Ras-Akt network in immune cell migration and polarity, in addition to its established role in growth factor receptor mediated signaling in mammalian cells (illustrated in the grey dashed box). In the migrating neutrophil cartoon, F-actin polymerization is shown as a meshwork of blue lines and actomyosin is denoted by a solid orange line.

Chemoattractants trigger a huge cascade of signal transduction and cytoskeletal events, yet our studies show that local activation or inactivation of single components can enhance existing polarity or reverse it and reestablish migration in a new direction. The signal transduction and cytoskeletal networks are thought to comprise of extensive crosstalk and feedback interactions^5, 67^. To examine whether all these events are activated by our perturbations are beyond the scope of this study. However, in our perturbations of Ras or Akt, we observed sustained protrusion-driven persistent migration, accompanied with changes in pre-existing polarity. It is difficult to conceive of such definite cell shape and migratory changes if global network activation or feedback between important network nodes were not triggered by our perturbations. This suggests that traditional concepts of “upstream” and “downstream” pathway components, derived from early studies, is an outdated and simplistic view. It remains to be seen whether we have chosen particularly key nodes in the networks, namely Ras and Akt, or activation of any single component will trigger activation of the entire network.

Previous studies have attributed polarity in neutrophils to various factors. A compelling argument was made for membrane tension as an inhibitor of cytoskeletal activation^103–105^. Our results are consistent with the membrane tension model since eliciting a protrusion at the back with RasGRP4 is accompanied by a collapse of the front, and conversely, causing a collapse of a front with RASAL3 leads to protrusion formation in the erstwhile back. Other experiments suggest that the cytoskeleton, but not its dynamics, is essential for polarity^85^. Our study does not test this model since our experiments always involved dynamic cytoskeletal changes. However, using our optical tools, the role of the cytoskeleton could be tested in neutrophils immobilized with a cocktail of cytoskeleton-stalling inhibitors. Would optical localization of a single component, such as Ras, recruit another component in the signaling cascade to the same place, or would the localization of the second component be also subjected to the polarity axis? Interesting questions such as these could be answered using our system.

Although Ras modulators, GEF and GAP proteins, have been investigated in growth and inflammatory functions of immune cells, their role in directed migration is largely unexplored^95, 106–109^. We selected two Ras modulators, RasGRP4 and RASAL3, for our evaluation. Using optogenetics, we could transiently activate or inhibit Ras on any region of the cell perimeter. First, upon activating Ras with RasGRP4 at the back, cells formed new protrusions at site of recruitment. Second, when Ras was inhibited by RASAL3 at the cell front, pre-existing local protrusions disappeared, and the front was converted into a new back. Third, dynamically silencing Ras all over the cell boundary shuts off all spontaneous protrusive activity. Thus, we showed for the first time that transient Ras activation underlies each protrusion and orchestrates cytoskeletal arrangement. Altogether, RasGEFs and RasGAPs, through their regulation of Ras at different locations on the cortex, can have varied effects on polarity, protrusive activity, and migration.

Some of the most notable regulators of mammalian chemotaxis are PI3K/PIP3 and mTorC2^47, 52, 53, 105, 110^. PIP3 is localized at the leading edge of migrating cells but its precise role in chemotaxis has been debated^22, 34, 36–44^. In neutrophils, recruitment of PI3K and focal production of PIP3, elicits a local protrusion^20, 47^. In cells lacking PTEN, which have elevated levels of PIP3, protrusion formation is drastically altered^111–113^. Furthermore, PIP3 was required for optimal protrusion formation elicited by optogenetic local activation of Rac^20^. In our studies, PIP3 was required for basal motility but was largely dispensable for Ras-mediated protrusion formation. On the other hand, inhibitors of mTor, but not mTorC1, completely blocked basal and RasGEF-induced motility, suggesting that mTorC2 was critical for both. In *Dictyostelium,* TorC2 is important but not essential for chemotaxis^4, 61, 114, 115^. Using optogenetic approaches to combine perturbations and expansion to many more components will help unravel the networks.

Traditionally, growth factors activate PI3K which leads to phosphorylation of activation loop of Akt in almost all cell types^43, 116^. Here, we showed that Akt activation may take place even in absence of PI3K activity. When PI3Kγ was pharmacologically inhibited in neutrophils, protrusion formation, polarity, and migration were drastically improved by optically activating Akt on the membrane. Here, Akt may either be phosphorylated by PDKs independent of PIP3, or another kinase, in absence of PI3K, may generate PIP3 for Akt activation^117, 118^. Thus, functional Akt activation can take place in absence of productive PI3K signaling.

Despite being activated through similar mechanisms, it has been reported that Akt isoforms perform different roles in growth and metabolism^119, 120^. In cell migration, it has been suggested that Akt1 and Akt2 serve opposing functions^121–125^. Even with the same isoform, opposing functions have been implied in different experiments^76, 101^. Our studies suggest that these differences might be attributable to selective genetic compensation, which can occur in knockout and overexpression studies, or to different assays in different cell types. Using our optogenetic tools to bypass such experimental artefacts, we demonstrated that, irrespective of isoform or cell type, spontaneous Akt activation can promote cell polarization and motility.

Current neutrophil-based therapeutics to treat diseases, such as COPD, cystic fibrosis, viral infections, and cancers, target only neutrophil cell surface chemokine receptors^126^. Aberrant chemokine signaling in neutrophil directed migration cause these diseases. Since chemokine receptor stimulation activates multiple downstream signaling pathways, pharmacological inhibition of these receptors has several off-target effects. Our optogenetic study in the model leukemic neutrophils showed that targeting activities of individual downstream components of these signaling cascades can have strong effects on cell motility and polarity. We provided proof-of-concept that targeting specific signaling molecules affect neutrophil behavior, but with greater control. Although neutrophil’s complex and intimate involvement in many diseases make them ideal therapeutic targets, measures are required to delimitate their harmful side-effects from beneficial responses. Single component regulation of neutrophil physiology, as shown in our study, will go a long way to serve this purpose.

## Acknowledgements

We thank all members of the Peter Devreotes, Pablo Iglesias, Douglas Robinson, and Miho Iijima laboratories (School of Medicine and Whiting School of Engineering, JHU) for helpful discussions and providing resources. We are grateful to Pablo Iglesias (Whiting School of Engineering, JHU) for providing kymograph codes. We thank Huiwang Zhan (Devreotes Lab) for assisting with a few experiments. We thank Orion Weiner (UCSF) and N. Gautam (WUSTL) for providing HL-60 and RAW 264.7 cell lines, respectively. We acknowledge Sean Collins (UC Davis) and Akihiro Kusumi (OIST, Japan) for providing plasmids. We are grateful to Stephen Gould and Shang-Jui Tsai (School of Medicine, JHU) for help with instrumentation. We thank Xiaoling Zhang and Ross Research Flow Cytometry Core, JHU for cell sorting services. We acknowledge Addgene and dictyBase for providing plasmids. This work was supported by NIH grant R35 GM118177 (to PND), DARPA HR0011-16-C-0139 (to PAI and PND), AFOSR MURI FA95501610052 (to PND), as well as NIH grant S10 OD016374 (to S Kuo of the JHU Microscope Facility).

## Author Contributions

DSP and PND conceived and designed the project, with inputs from TB; DSP developed optogenetic constructs, engineered neutrophil stable cell lines, designed and executed experiments in neutrophils with contribution from TB; DSP and TB designed and carried out macrophage experiments; DSP, TB, and YL designed *Dictyostelium* experiments, and YL and TB performed them; DSP performed majority of data analyses with input from TB and PND; TB generated kymographs and executed MATLAB analyses; JB generated constructs; DSP wrote final version of the manuscript based on contributions from PND, TB, YL, and JB; PND supervised the study.

## Materials and Methods

### Reagents and inhibitors

Fibronectin (Sigma-Aldrich; F4759-2MG) was dissolved in 2 ml sterile water, followed by dilution in 8 ml PBS to make a stock solution of 200 µg/ml. N-Formyl-Met-Leu-Phe (fMLP, Sigma-Aldrich; 47729) was dissolved in DMSO (Sigma Aldrich; D2650) to make stock solution of 10 mM. FKP-(D-Cha)-Cha-r (Anaspec; 65121) was dissolved in PBS to prepare a 2.5 mM stock solution. AS605240 (Sigma-Aldrich; A0233) or PP242 (EMD Millipore; 475988) was dissolved in DMSO to make a stock solution of 20 mM. Pertussis toxin (PTX, Sigma-Aldrich; 516560-50UG) was reconstituted in 500 µl sterile water to prepare a 100 µg/ml stock. Gallein (Sigma-Aldrich; 371708-5MG) or CK-666 (EMD Millipore; 182515) was dissolved in DMSO to make 20 mM or 50 mM stock solution, respectively. Janelia Fluor 646 HaloTag (Promega Corporation; GA1120) was dissolved in DMSO to make a 200 μM stock. 2.5 mg/ml puromycin (Sigma-Aldrich; P8833) or 10 mg/ml blasticidine S (Sigma-Aldrich; 15205) stock was prepared in sterile water. Hygromycin B (Thermo Fisher Scientific; 10687010) or G418 sulphate (Thermo Fisher Scientific; 10131035) was available as ready-made 50 mg/ml stock solution. All stocks were aliquoted and stored at −20^0^C. According to experimental requirements, further dilutions were made in PBS or culture medium before adding to cells.

### Cell culture and differentiation

Human neutrophil-like HL-60 cell line was a kind gift from Dr. Orion Weiner (UCSF). Cells were cultured in RPMI medium 1640 containing L-glutamine and 25 mM HEPES (Gibco; 22400-089). This medium was supplemented with 15% heat-inactivated fetal bovine serum (FBS; Thermo Fisher Scientific; 16140071) and 1% penicillin-streptomycin (Thermo Fisher Scientific; 15140122). For subculturing, cells were split at a density of 0.15 million cells/ml and passaged every 3 days. Stable lines were maintained similarly, in presence of selection antibiotics. For differentiating, 1.3% DMSO was added to cells at a density of 0.15 million cells/ml, followed by incubation for 5-7 days, before experimentation^127^. This myeloid leukemia cell line, upon differentiation, serves as an effective model to investigate human neutrophils^59, 128^. Stable lines of HL-60 cells were similarly differentiated, and selection antibiotics were removed for experimentation.

Mouse monocyte/macrophage RAW 264.7 cell line was obtained from Dr. N. Gautam (Washington University School of Medicine). Cell line was maintained in DMEM medium containing 4.5 g/L glucose, sodium pyruvate, and sodium bicarbonate (Sigma-Aldrich; D6429). Culture medium was supplemented with 10% FBS and 1% penicillin-streptomycin. At ∼90% confluency, cells were harvested by scraping and subcultured at a split ratio of 1:4^129^.

Human embryonic kidney HEK293T cell line was grown in DMEM medium containing 4.5 g/L glucose and sodium pyruvate (Gibco; 10569-010). Medium was supplemented with 10% FBS and 1% penicillin-streptomycin. Cells were maintained according to ATCC guidelines.

All mammalian cells were maintained under humidified conditions at 37^0^C and 5% CO_2_, and experiments were done with cells at low passage.

Wild type *Dictyostelium discoideum* cells of the axenic strain AX2 were used in this study. Cells were subcultured, either in suspension or on tissue culture dishes, in HL-5 medium at 22^0^C. Stable cell lines, generated by electroporation, were grown in presence of hygromycin B and G418 sulphate. All experiments were carried out within 2 months of thawing the cells from frozen stocks.

### Cloning

All DNA oligonucleotides were ordered from Sigma-Aldrich. For lentiviral constructs, CIBN-CAAX (Addgene #79574) or LifeActmiRFP703 (Addgene #79993) ORF was subcloned into pLJM1-eGFP plasmid (Addgene #19319), in place of the eGFP gene, using primer sets P1/P2 or P3/P4 respectively (Table S1). We procured PiggyBac™ transposon system from Dr. Sean Collins (UC Davis) which consists of a transposon and a transposase expression plasmid^130^. CRY2PHR-mcherry2 gene (Addgene #26866) was first sub-cloned into the transposon plasmid using primers P5/P6. Next, at the C-terminal of CRY2PHR-mcherry2 gene in the transposon plasmid, we introduced HRas G12V ΔCAAX (Addgene #18666), KRas4B G12V ΔCAAX (Addgene #9052), RasGRP4 (Addgene #70533), RASAL3 (Addgene #70521), Akt1 (Addgene #86631), Akt1_T308A_ (Addgene #49189), or Akt2 (Addgene #86593) gene using primer sets P7/P8, P9/P10, P11/P12, P13/P14, P15/P16, P17/P18, or P19/P20, respectively (Table S1). pFUW2-RFP-PH-Akt construct was developed previously in our lab^131^.

For non-viral constructs, RASAL3 (Addgene #70521) was subcloned into pCRY2PHR– mCherryN1 plasmid (Addgene #26866), at the C-terminal of CRY2PHR-mcherry2 gene, using primers P21/22 (Table S1). pEGFPN1-Lifeact-7Alinker-Halo7 plasmid was a kind gift from Dr. Akihiro Kusumi (Okinawa Institute of Science and Technology Graduate University).

For *Dictyostelium* constructs, Venus-iLID-CAAX (Addgene #60411) or tgRFPt-SSPB R73Q (Addgene #60416) gene was subcloned in pDM358 (dictyBase #534) or pCV5 (dictyBase #23) plasmid using AgeI/SbfI or AgeI/BamHI restriction digestion, respectively. Next, at the C-terminal of tgRFPt-SSPB R73Q gene in pCV5, we introduced PKBA gene using primer sets P23/24 (Table S1).

All constructs were verified by diagnostic restriction digestion and Sanger sequencing (JHMI Synthesis and Sequencing Facility).

### Stable cell line construction

Stable expression lines in HL-60 cells were generated by a combination of 3^rd^-generation lentiviral- and PiggyBac™ transposon-integration based approaches^18, 132^. Virus was prepared in HEK293T cells grown to ∼80% confluency in 10 cm cell culture dish (Greiner Bio-One; 664160). For each reaction, a mixture of 3.32 µg pMD2.G (Addgene plasmid #12259), 2 µg pMDLg/pRRE (Addgene plasmid #12251), 4.64 µg pRSV-Rev (Addgene plasmid #12253), and 10 µg pLJM1 construct with gene of interest (CIBN-CAAX or LifeActmiRFP703) or pFUW2-RFP-PH-Akt construct were transfected using Lipofectamine 3000® as per manufacturer’s instructions (Invitrogen; L3000-008). After 96 hours, virus containing culture medium was harvested at 3000 rpm for 20 mins at 4°C. In a 6-well plate (Greiner Bio-One; 657160), entire viral medium was added to 4×10^6^ HL-60 cells (seeded at a density of 0.25×10^6^ cells/mL) in presence of 10 µg/mL polybrene (Sigma; TR1003). After 24 hours, viral medium was removed, and cells were introduced to a mix of fresh and conditioned (mixed) culture medium. For selecting LifeActmiRFP703 or RFP-PH-Akt expressors, infected cells were sorted after 5 days, and subsequently, grown to confluency. For selecting CIBN-CAAX-expressors, infected cells were allowed to recover for 24 hours, and subsequently, incubated with 1 µg/mL puromycin in 24-well cell culture plate (Greiner Bio-One; 662160) for 4-5 days. Resistant cells were grown to confluency in puromycin.

For transposon integration in HL-60 cell line, 5 µg transposon plasmid, containing CRY2PHR-mcherry2 fused to a gene of interest, was co-electroporated with an equal amount of transposase expression plasmid into two million cells using Neon™ transfection kit (Invitrogen; MPK10025B). Cells and DNA mix were resuspended in buffer ‘R’ before electroporation in 100 µl pipettes at 1350 volts for 35 ms in Neon™ electroporation system (Invitrogen; MPK5000). Cells were resuspended in mixed culture medium in a 6-well plate and allowed to recover for 24 hours. Post-recovery, transfected cells were selected in presence of 10 µg/mL blasticidine S for 5-6 days. Once blasticidine S was removed, resistant cells were transferred to 48-well cell culture plate (Sarstedt; 83.3923), and further grown over 3-4 weeks into stable cell lines. Stable cells were maintained throughout in Blasticidine S.

Stable lines in *Dictyostelium* cells were generated by electroporation as described previously^115^. Briefly, 5×10^6^ cells were harvested, washed and resuspended in 100 μl chilled H-50 buffer. Next, 2 μg each of Venus-iLID-CAAX/pDM358 and tgRFPt-SSPB R73Q-CTRL/pCV5 (control) or tgRFPt-SSPB R73Q-PKBA/pCV5 plasmids were mixed with cell suspension, and transferred to chilled 0.1 cm-gap cuvette (Bio-Rad, 1652089). Transfection took place over two rounds of electroporation at 850 V and 25 μF with an interval of 5 secs (Bio-Rad Gene Pulser Xcell Electroporation Systems). Electroporated cells were incubated on ice for 10 mins, and subsequently transferred to HL-5 culture medium, supplemented with heat-killed *Klebsiella aerogenes* (lab stock), in 10 cm cell culture dish. On the following day, 50 μg/ml hygromycin B and 20 μg/ml G418 sulphate were added to cells and selected over 3-4 weeks.

### Transient transfection

RAW 264.7 macrophage-like cells were transiently transfected by nucleofection using Amaxa cell line kit V (Lonza; VACA-1003) as previously described^115, 129^. Briefly, 3×10^6^ cells were harvested and added to 100 µl supplemented Nucleofector Solution V containing 0.7 µg CIBN-CAAX (Addgene #79574), 1.1 µg CRY2PHRmcherry2-RASAL3, and 0.7 µg pEGFPN1-Lifeact-7Alinker-Halo7. Cell and DNA were mixed gently, transferred to a Lonza cuvette and electroporated with Amaxa Nucleofector II device using program ‘D-32’. After a single pulse, cells were transferred carefully to 0.5 ml pre-warmed culture medium and incubated at 37^0^C and 5% CO_2_ for 10 mins. Subsequently, 2×10^5^ cells were transferred to each well of 8-well chambered coverglass (LAB-TEK; 155409) and left in the incubator for 1 hour. Next, 0.5 ml warm culture medium was added to each sample and incubated for 4 hours before imaging.

### Microscopy and FACS

Two microscopes were used for time-lapse imaging: 1) Zeiss LSM780-FCS single-point, laser scanning confocal microscope (Zeiss Axio Observer with 780-Quasar confocal module) and 2) Zeiss LSM800 GaAsP single-point, laser scanning confocal microscope with a wide-field camera. In these microscopes, argon laser (488 nm excitation) was used for GFP or Venus visualization, solid-state laser (561 nm excitation) was used for mcherry2 or RFP, and diode laser (633 nm excitation) was used for miRFP703 or Janelia Fluor 646 HaloTag. 40X/1.30 Plan-Neofluar oil objective was used, along with digital zoom. Microscopes were equipped with temperature-controlled chamber held at 5% CO_2_ and 37^0^C for imaging mammalian cells. Zeiss 780 and 800 were operated by ZEN Black and ZEN Blue software, respectively.

For high-speed sorting, we used two instruments: 1) BD FACSAria IIu cell sorter and 2) SH800S cell sorter (Sony). We used 561 nm excitation laser to sort for RFP expression, and 633 nm excitation to sort miRFP703 expressors. Briefly, cells were harvested, resuspended in sorting buffer (1x PBS, Ca^2+^/Mg^2+^ free; 0.9% heat-inactivated FBS; 2% penicillin-streptomycin) at a density of 15×10^6^ cells/ml, and sorted using 100 μm microfluidic sorting chip. High expressors (top 1%-10%) were collected in fresh culture medium (containing 2% penicillin-streptomycin) and grown to confluency.

### Optogenetics

All optogenetic experiments with differentiated HL-60 cells were done in absence of chemoattractant, fMLP. On day of the experiment, differentiated neutrophils were allowed to attach on chambered coverglass, coated with fibronectin at a density of ∼35 µg/cm^2^, for 40 mins. Next, depending on experimental requirements, attached cells were treated with inhibitors (20 µM AS605240, 20 µM PP242, 20 µM gallein, or 100 µM CK-666) for 10 mins before imaging. For pertussis toxin (PTX) treatment, differentiated cells were incubated with 1 µg/ml PTX for 20 hours before imaging. For global recruitment experiments, 488 nm excitation laser was switched on after imaging for at least 300 secs. Photoactivation and image acquisition were done once every 7 secs for single plane imaging. Laser intensity during image capture was at a low level (laser power of ∼0.05 mW at the objective) so that protein recruitment over the entire cell periphery was maintained throughout without inducing light damage. For local recruitment studies, a small region of interest was drawn (shown as dashed white box in images or solid yellow box in videos), which was irradiated with 488 nm laser, at low power, in multiple iteration.

We discovered that pre-treatment of differentiated, migration-competent HL-60 cells with heat-killed *Klebsiella aerogenes* vastly improved efficiency of CRY2-CIBN optogenetic system. Briefly, we treated 10^7^ differentiated cells, grown on 10 cm cell culture dish, with 13 µg/ml heat killed *Klebsiella aerogenes* (lab stock) for 7 hours. Dead bacteria were rinsed off along with non-adherent neutrophils on fibronectin-coated chambered coverglass, and all imaging was completed within 5 hours.

Transiently transfected RAW 264.7 cells were transferred to 450 µL pre-warmed HBSS buffer supplemented with 1 g/L glucose, and subsequently stained with 5 nM Janelia Fluor 646 HaloTag for 15 mins. During imaging, cells were activated with 10 μM C5a-receptor agonist FKP-(D-Cha)-Cha-r. Local recruitment studies were performed similarly to HL-60 cells.

Optogenetic experiments with *Dictyostelium* cells were done in absence of any chemoattractant. Cells were allowed to attach on chambered coverglass for 30 mins before imaging. For recruitment experiments, 488 nm excitation laser was switched on after imaging for at least 250 secs. Photoactivation and image acquisition were done once every 5-10 secs for single plane imaging. Very low laser intensity (10-20% of what was used for mammalian optogenetics) was sufficient to stably recruit SspB over the cell perimeter without inducing light damage.

### Akt phosphorylation assay and immunoblotting

This assay was performed as described previously with minor modifications^59^. Briefly, differentiated HL-60 cells, pre-treated with heat-killed bacteria, were harvested, resuspended in starvation medium (culture medium without FBS) at a density of 2×10^7^ cells/mL, and kept on ice for 10 mins. During this time, cells were treated with 20 µM AS605240 or PP242. Next, while shaking at 200 rpm, 10 µM fMLP was added to untreated or inhibitor-treated cells. At indicated timepoints, 125 µl of cell suspension was collected from untreated or treated samples, mixed with SDS sample buffer, and boiled for 5 mins.

Samples were subjected to SDS-PAGE (4-15% polyacrylamide; Bio-Rad). Protein equivalent to at least 2.5×10^5^ cells was loaded per well for each sample. Akt phosphorylation at Thr308 or Ser473 (∼60 kDa) was detected with rabbit anti-phospho-Akt_T308_ (Cell Signaling; 13038) or anti-phospho-Akt_S473_ (Cell Signaling; 4060) antibody, respectively. Total Akt (∼56 kDa) was detected with rabbit anti-pan Akt antibody (Cell Signaling; 9272). All primary antibodies were used at a dilution of 1:1000. After overnight primary antibody incubation at 4^0^C, blots were probed with goat anti-rabbit IRDye 680RD-cojugated secondary antibody (1:10,000 dilution; Li-Cor; 925-68071) for 1 hour. Near-infrared signal from the blots was detected via Odyssey CLx imaging system (Li-Cor).

### Image Analysis

Images were analyzed on Fiji/ImageJ 1.52i (NIH) and MATLAB 2019a (MathWorks, MA, USA) as described previously^115^. Results were plotted using GraphPad Prism 8 (GraphPad software, CA, USA) and Microsoft Excel (Microsoft, WA, USA).

#### Linescan intensity profile

Linescans were generated in Fiji/ImageJ 1.52i software. A straight line segment (width of 12 pixels) was drawn on the red channel, using the line tool option, across the cell. Using the “Plot Profile” option, we obtained the average intensity value along that line for the red channel. Values were normalized and graphed in Microsoft Excel.

#### Kymographs

Cell segmentation was done against the background using a custom code written in MATLAB 2019b, after standard image processing steps were carried out. Membrane kymographs were generated from segmented cells as described earlier^133^. We used a linear color map for normalized intensities, where blue indicated lowest intensity and yellow denoted highest.

#### Cell migration analysis

Analysis was performed by segmenting neutrophil cells in Fiji/ImageJ 1.52i software. For this, image stack was thresholded using the ‘Threshold’ option. ‘Calculate threshold for each image’ box was unchecked, and range was not reset. Next, cell masks were created by size-based thresholding using the ‘Analyzed particles’ option. To optimize binarized masks, ‘Fill holes’, ‘Dilate’, and ‘Erode’ were done several times. For creating temporal color-coded cell outlines, ‘Outline’ was applied on binarized masks, followed by ‘Temporal-Color Code’ option. Next, ‘Centroid’ and ‘Shape descriptors’ boxes were checked in ‘Set Measurements’ option under ‘Analyze’ tab. This provided us with values for centroid coordinates and aspect ratio. Mean and SEM from replicates of aspect ratio values were determined and plotted in GraphPad Prism 8. The starting point for centroid values was set to zero for each track, and these new coordinates were plotted in Microsoft Excel to generate migration tracks. Before- and after-recruitment tracks for a cell have the same color-code to aid comparison. Velocity was calculated by computing displacement between two consecutive frames. Displacement was then divided by time interval to obtain speed for each cell. These speed values were then time-averaged over all frames to produce data points for cell speed which were plotted as box-and-whisker graphs in GraphPad Prism 8.

#### Local protrusion formation analysis

The region of recruitment in the red channel was marked using the ‘segmented line’ tool in ImageJ software. Next, ‘Fit spline’ and ‘Straighten’, two custom-written macros, were sequentially used to determine the midpoint of the recruited region. The centroid was estimated with help of another ImageJ macro. Holding centroid as vertex, the angle between the midpoint of recruitment region and new protrusion was determined using the ‘angle’ tool. In MATLAB, these values were plotted with help of ‘polarhistogram’ command. Minimum number of bins for each plot were determined by Sturges’ formula. For each histogram, atleast 40 fresh protrusions were considered.

### Statistical Analysis

Statistical analyses were performed by paired or unpaired 2-tailed non-parametric tests on GraphPad Prism 8 and Microsoft Excel. All results are expressed as mean ± SD from at least 3 independent experiments. ns denotes P>0.05, * denotes P ≤ 0.05, ** denotes P ≤ 0.01, *** denotes P ≤ 0.001, **** denotes P ≤ 0.0001.

## Data availability

All data are provided in the main or supplementary text. Request for additional information on this work are to be made to the corresponding author.

## SUPPLEMENTARY INFORMATION

### Supplementary Figures

**Figure S1.**
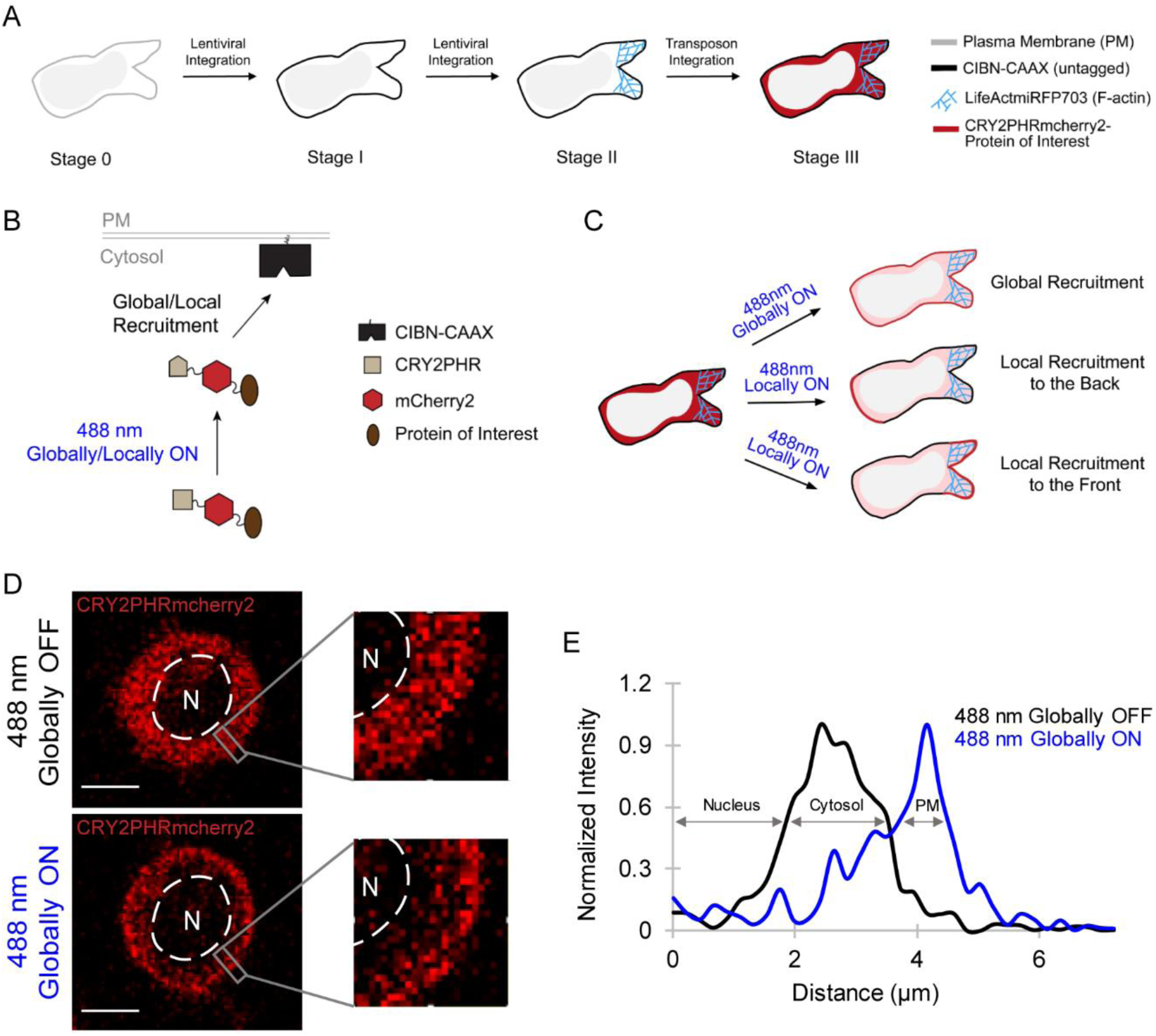
Construction and validation of CRY2-CIBN optogenetic system in neutrophils. **(A)** Cartoon showing stepwise introduction of CRY2-CIBN optogenetics in HL-60 cells. In wild type HL-60 cells (Stage 0), CIBN-CAAX membrane anchor (Stage I) and F-actin polymerization biosensor, LifeActmiRFP703 (Stage II), were introduced sequentially by lentiviral integration. These cells were used to express cytosolic CRY2PHR-mcherry2 fused with protein of interest by transposon-based integration (Stage III). **(B)** Schematic demonstrating recruitment of cytosolic CRY2PHR-mcherry2 fused with protein of interest to the membrane CIBN-CAAX by local or global illumination with 488 nm laser. **(C)** Depending on the region of the membrane where 488 nm laser was applied, CRY2PHR-mcherry2-protein of interest fusion was recruited all over the cell periphery (global recruitment), or specifically to the back or front of the cell. **(D)** Representative cell showing CRY2PHR-mcherry2 in the cytosol when 488 nm laser was off (top panel). Once light was switched on, CRY2PHR-mcherry2 translocated evenly to the periphery (bottom panel). The circular dashed line region with ‘N’ denotes the large nucleus in these cells. Images are representative of many cells from atleast three independent experiments. Scale bars represent 5 µm. **(E)** Linescan across the cytosol and membrane of the cell (denoted by white box in D), before (black) or after (blue) 488 nm laser was switched on globally. There is a distinct shift in CRY2PHR-mcherry2 intensity peak from cytosol to plasma membrane (PM) when laser was on.

**Figure S2.**
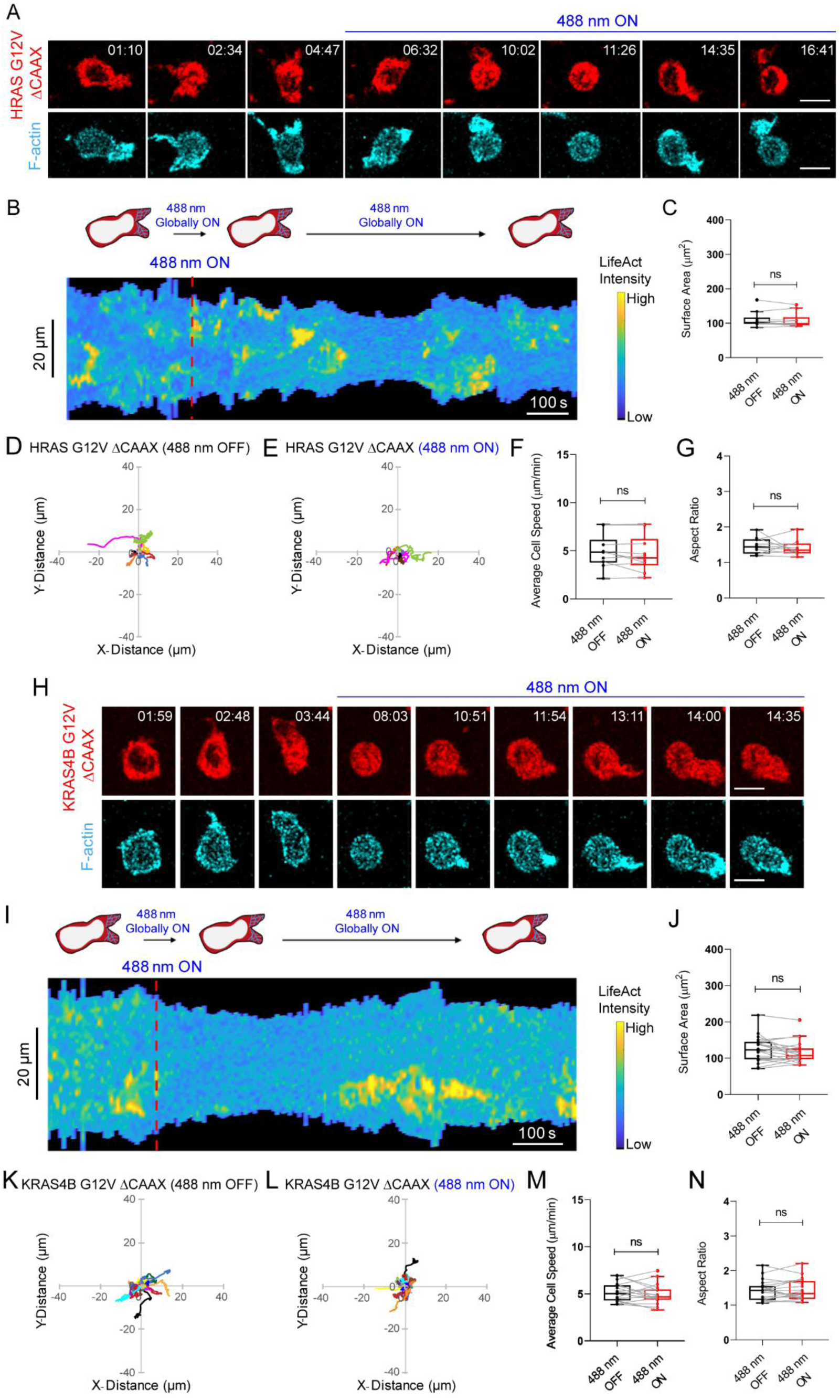
Cytosolic form of constitutively active Ras isoforms does not affect neutrophil polarity and migration. **(A)** Time-lapse confocal images of differentiated HL-60 neutrophil expressing CRY2PHR-mcherry2-HRas G12V ΔCAAX (red; upper panel) and LifeActmiRFP703 (cyan; lower panel), before or after 488 nm laser was switched on globally. No appreciable HRas G12V ΔCAAX recruitment was observed presumably due to low expression of CIBN-CAAX membrane anchor. Time in min:sec format. Scale bars represent 5 µm. **(B)** Representative kymograph of cortical LifeAct intensity in HRas G12V ΔCAAX-expressing neutrophil before or after 488 nm laser was turned on. A linear color map shows that blue is the lowest LifeAct intensity whereas yellow is the highest. Duration of the kymograph is 17 mins. Cartoon depicts recruitment, F-actin polymerization or cell shape status corresponding to the kymograph. **(C)** Box-and-whisker plot of neutrophil surface area before (black) or after (red) 488 nm laser was turned on. n_c_=10 from atleast 3 independent experiments; ns denotes P>0.05 (Wilcoxon-Mann-Whitney rank sum test). Centroid tracks of neutrophils (n_c_=10) showing random motility before **(D)** or after **(E)** 488 nm laser was turned on. Each track lasts atleast 5 mins and was reset to same origin. Box-and-whisker plots of neutrophil speed **(F)** and aspect ratio **(G)** before (black) or after (red) 488 nm laser was switched on. n_c_=10 from atleast 3 independent experiments; ns denotes P>0.05 (Wilcoxon-Mann-Whitney rank sum test). **(H)** Time-lapse confocal images of differentiated neutrophil expressing CRY2PHR-mcherry2-KRas4B G12V ΔCAAX (red; upper panel) and LifeActmiRFP703 (cyan; lower panel), before or after 488 nm laser was switched on globally. No appreciable KRas4B G12V ΔCAAX recruitment was observed presumably due to low expression of CIBN-CAAX membrane anchor. Time in min:sec format. Scale bars represent 5 µm. **(I)** Representative kymograph of cortical LifeAct intensity in KRas4B G12V ΔCAAX -expressing neutrophil before or after 488 nm laser was turned on. A linear color map shows that blue is the lowest LifeAct intensity whereas yellow is the highest. Duration of the kymograph is 15 mins. Cartoon depicts recruitment, F-actin polymerization or cell shape status corresponding to the kymograph. **(J)** Box- and-whisker plot of neutrophil surface area before (black) or after (red) 488 nm laser was turned on. n_c_=20 from atleast 3 independent experiments; ns denotes P>0.05 (Wilcoxon-Mann-Whitney rank sum test). Centroid tracks of neutrophils (n_c_=20) showing random motility before **(K)** or after **(L)** 488 nm laser was turned on. Each track lasts atleast 5 mins and was reset to same origin. Box- and-whisker plots of neutrophil speed **(M)** and aspect ratio **(N)** before (black) or after (red) 488 nm laser was switched on. n_c_=20 from atleast 3 independent experiments; ns denotes P>0.05 (Wilcoxon-Mann-Whitney rank sum test).

**Figure S3.**
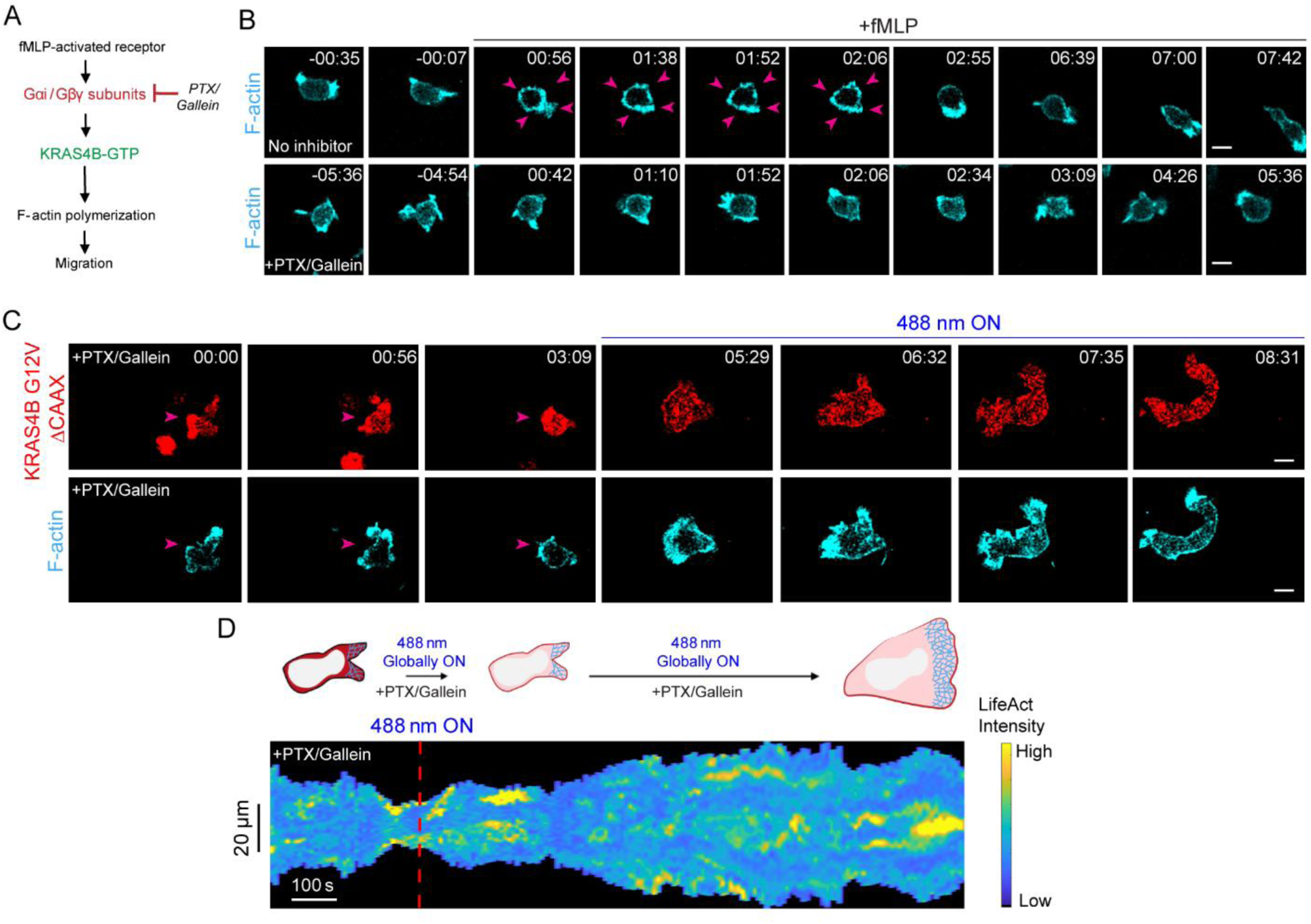
Spontaneous Kras4B activation can induce neutrophil polarization and motility in absence of G-protein signaling. **(A)** Proposed model highlighting KRas4B G12V ΔCAAX activity, in absence of G-protein signaling, promotes F-actin polymerization and migration. A combination of Gαi and Gβγ inhibitors, pertussis toxin (PTX) and gallein respectively, were used to test this. **(B)** Time-lapse confocal images of untreated (upper panel) or PTX/gallein-treated (lower panel) HL-60 neutrophil expressing LifeActmiRFP703 (cyan), before or after a uniform stimulus of 100 nM fMLP was applied. Within a minute of fMLP addition, we observed a global burst of F-actin polymerization, as indicated with increased LifeActmiRFP703 on the membrane (shown with pink arrows), in untreated cells. This did not occur in cells pre-treated with Gαi/ Gβγ inhibitors. Images are representative of many cells from atleast three independent experiments. Time in min:sec format. Scale bars represent 5 µm. **(C)** Time-lapse confocal images of PTX/gallein-treated HL-60 neutrophil expressing CRY2PHR-mcherry2-KRas4B G12V ΔCAAX (red; upper panel) and LifeActmiRFP703 (cyan; lower panel), before or after 488 nm laser was turned on globally. Pink arrows denote representative cell. Images are representative of many cells from atleast three independent experiments. Time in min:sec format. Scale bars represent 5 µm. **(D)** Representative kymograph of cortical LifeAct intensity in PTX/gallein-treated KRas4B G12V ΔCAAX-expressing neutrophil before or after 488 nm laser was turned on. A linear color map shows that blue is the lowest LifeAct intensity whereas yellow is the highest. Duration of the kymograph is 10 mins. Cartoon depicts membrane recruitment, actin polymerization or cell shape status corresponding to the kymograph.

**Figure S4.**
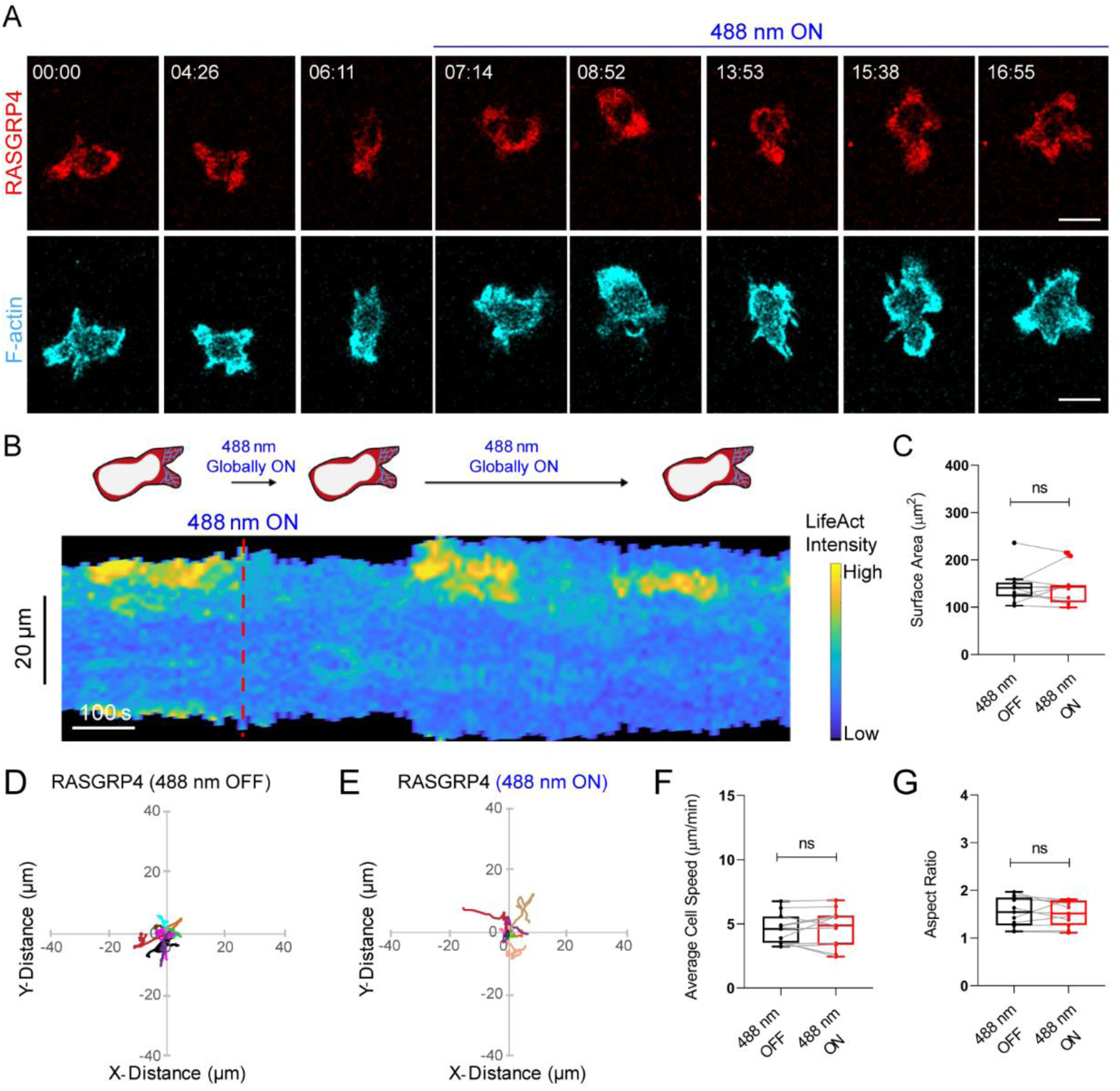
Cytosolic RasGRP4 does not improve polarity and migration. **(A)** Time-lapse confocal images of differentiated HL-60 neutrophil expressing CRY2PHR-mcherry2-RasGRP4 (red; upper panel) and LifeActmiRFP703 (cyan; lower panel), before or after 488 nm laser was switched on globally. No appreciable RasGRP4 recruitment was observed presumably due to low expression of CIBN-CAAX membrane anchor. Time in min:sec format. Scale bars represent 5 µm. **(B)** Representative kymograph of cortical LifeAct intensity in RasGRP4-expressing neutrophil before or after 488 nm laser was turned on. A linear color map shows that blue is the lowest LifeAct intensity whereas yellow is the highest. Duration of the kymograph is 17 mins. Cartoon depicts recruitment, F-actin polymerization or cell shape status corresponding to the kymograph. **(C)** Box- and-whisker plot of neutrophil surface area before (black) or after (red) 488 nm laser was turned on. n_c_=11 from atleast 3 independent experiments; ns denotes P>0.05 (Wilcoxon-Mann-Whitney rank sum test). Centroid tracks of neutrophils (n_c_=11) showing random motility before **(D)** or after **(E)** 488 nm laser was turned on. Each track lasts atleast 5 mins and was reset to same origin. Box- and-whisker plots of neutrophil speed **(F)** and aspect ratio **(G)** before (black) or after (red) 488 nm laser was switched on. n_c_=11 from atleast 3 independent experiments; ns denotes P>0.05 (Wilcoxon-Mann-Whitney rank sum test).

**Figure S5.**
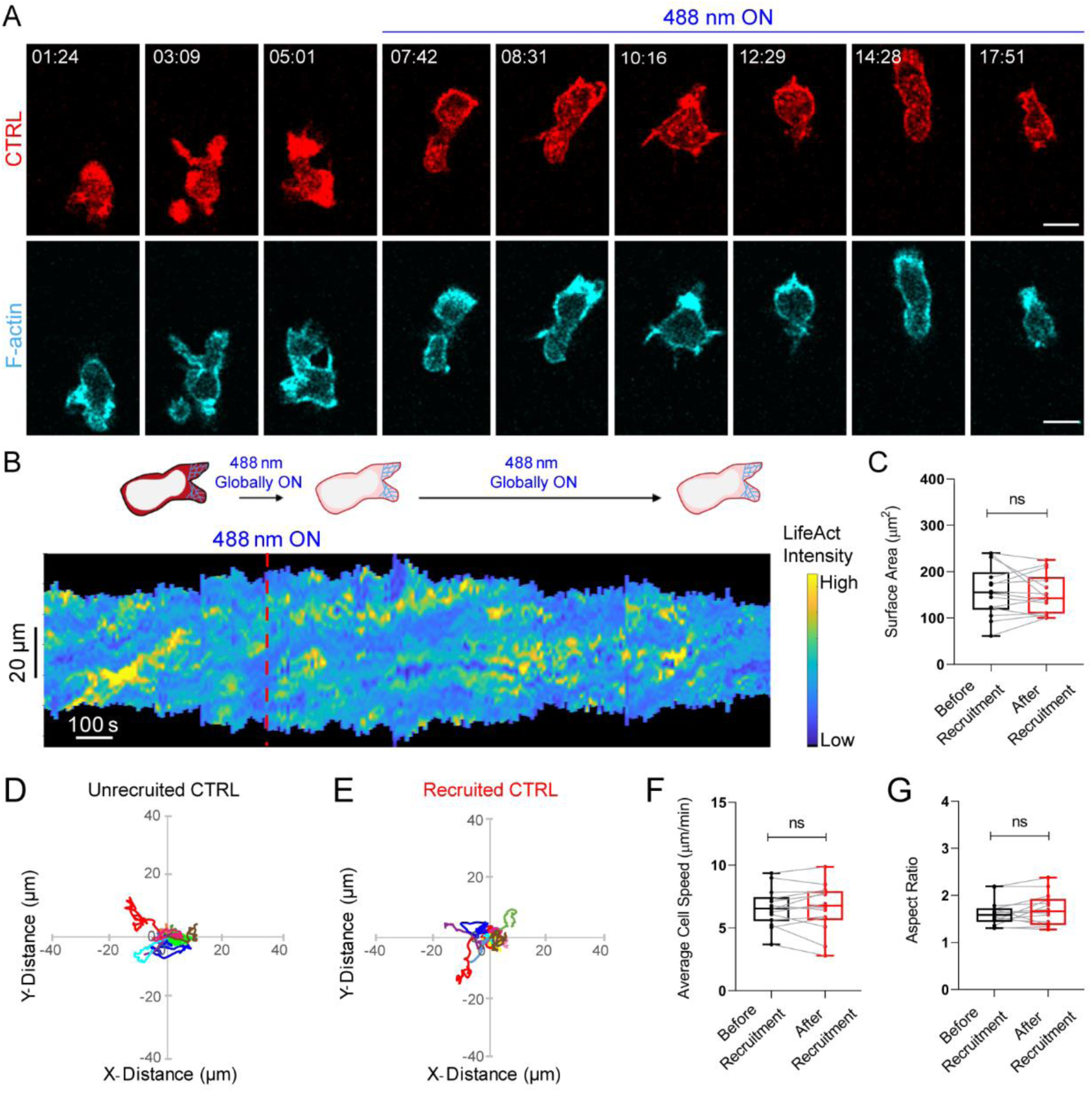
CRY2PHR recruitment on neutrophil membrane has no effect on polarity and migration. **(A)** Time-lapse confocal images of differentiated HL-60 neutrophil expressing CRY2PHR-mcherry2-CTRL (control; red; upper panel) and LifeActmiRFP703 (cyan; lower panel), before or after 488 nm laser was switched on globally. Time in min:sec format. Scale bars represent 5 µm. **(B)** Representative kymograph of cortical LifeAct intensity in CTRL-expressing neutrophil before or after 488 nm laser was turned on. A linear color map shows that blue is the lowest LifeAct intensity whereas yellow is the highest. Duration of the kymograph is 18 mins. Cartoon depicts membrane recruitment, actin polymerization or cell shape status corresponding to the kymograph. **(C)** Box-and-whisker plot of cell surface area before (black) or after (red) CTRL membrane recruitment. n_c_=14 from atleast 3 independent experiments; ns denotes P>0.05 (Wilcoxon-Mann-Whitney rank sum test). Centroid tracks of neutrophils (n_c_=14) showing random motility before **(D)** or after **(E)** CTRL membrane recruitment. Each track lasts atleast 5 mins and was reset to same origin. Box-and-whisker plots of neutrophil speed **(F)** and aspect ratio **(G)** before (black) or after (red) CTRL recruitment. n_c_=14 from atleast 3 independent experiments; ns denotes P>0.05 (Wilcoxon-Mann-Whitney rank sum test).

**Figure S6.**
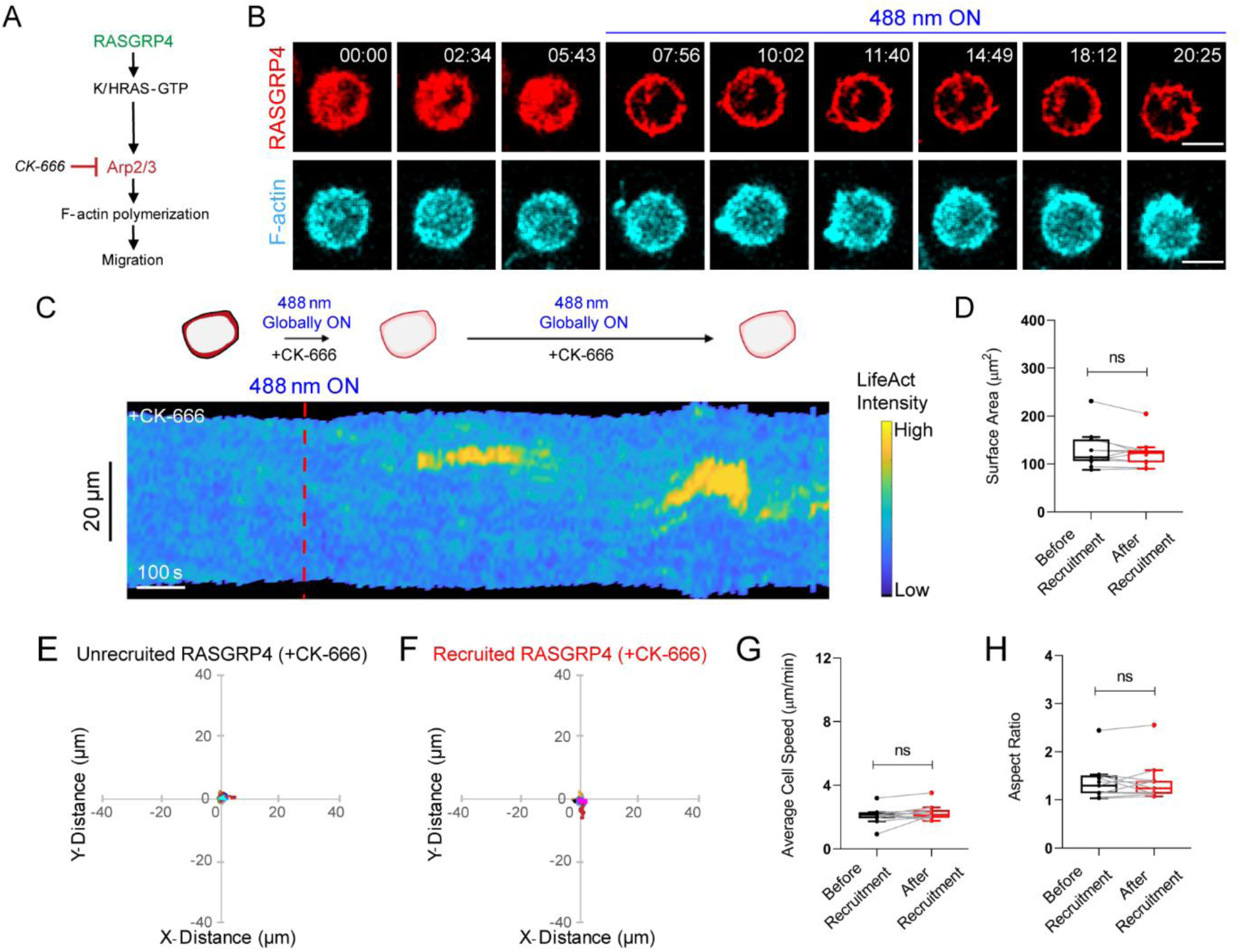
RasGRP4-mediated neutrophil activation works through Arp2/3 complex. **(A)** Proposed model highlighting RasGRP4 activation promotes F-actin polymerization and migration through Arp2/3 complex. CK-666 was the inhibitor used to test this. **(B)** Time-lapse confocal images of CK-666-treated HL-60 neutrophil expressing CRY2PHR-mcherry2-RasGRP4 (red; upper panel) and LifeActmiRFP703 (cyan; lower panel), before or after 488 nm laser was turned on globally. Time in min:sec format. Scale bars represent 5 µm. **(C)** Representative kymograph of cortical LifeAct intensity in CK-666-treated RasGRP4-expressing neutrophil before or after 488 nm laser was turned on. A linear color map shows that blue is the lowest LifeAct intensity whereas yellow is the highest. Duration of the kymograph is 21 mins. Cartoon depicts membrane recruitment, actin polymerization or cell shape status corresponding to the kymograph. **(D)** Box- and-whisker plot of cell surface area before (black) or after (red) RasGRP4 membrane recruitment. n_c_=11 from atleast 3 independent experiments; ns denotes P>0.05 (Wilcoxon-Mann-Whitney rank sum test). Centroid tracks of CK-666-treated neutrophils (n_c_=11) showing random motility before **(E)** or after **(F)** RasGRP4 membrane recruitment. Each track lasts atleast 5 mins and was reset to same origin. Box-and-whisker plots of speed **(G)** and aspect ratio **(H)** before (black) or after (red) RasGRP4 recruitment in CK-666-treated neutrophils. n_c_=11 from atleast 3 independent experiments; ns denotes P>0.05 (Wilcoxon-Mann-Whitney rank sum test).

**Figure S7.**
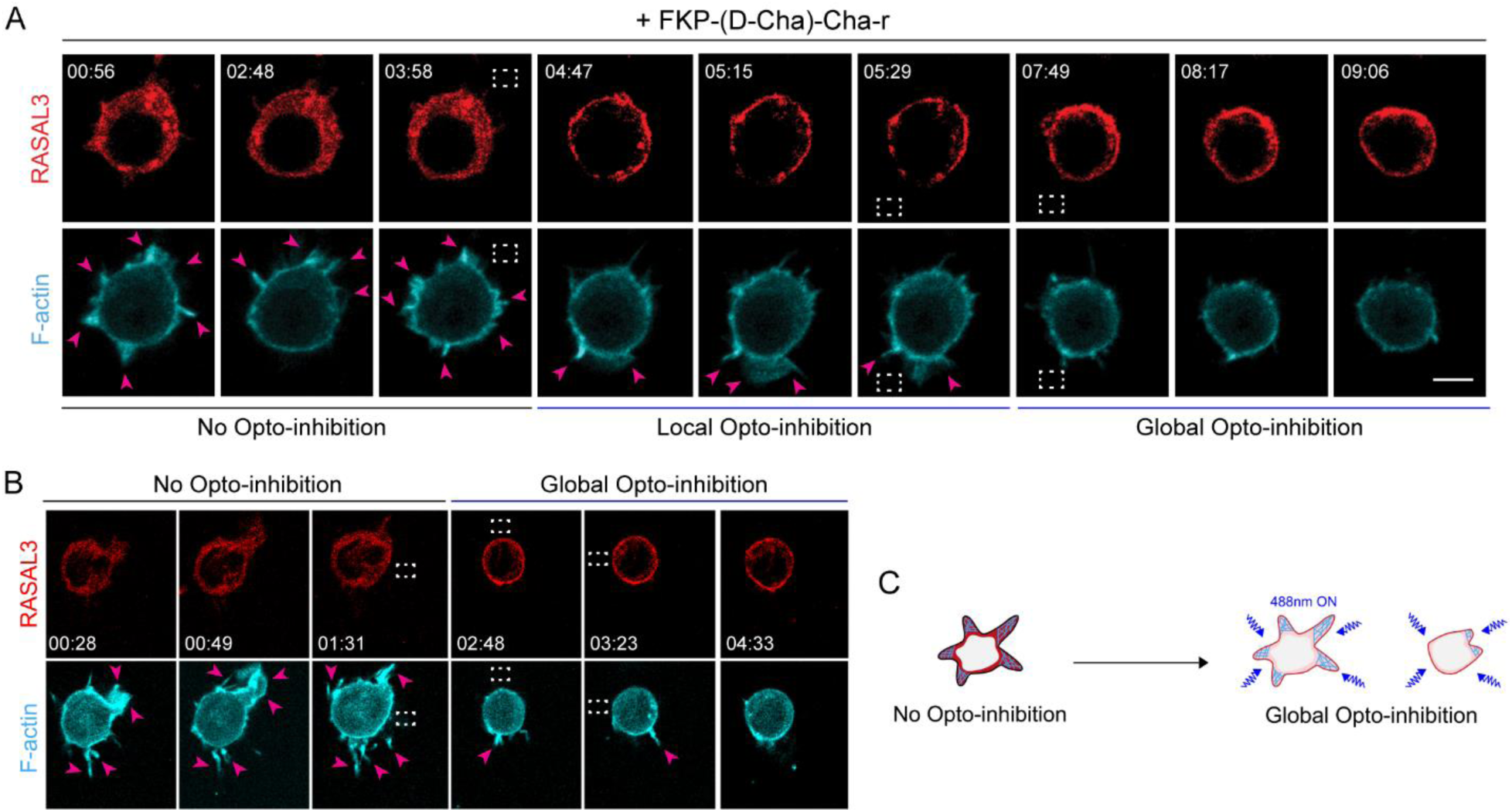
Ras inhibition shuts down protrusive activity in macrophages and neutrophils. **(A)** Time-lapse confocal images of FKP-(D-Cha)-Cha-r-treated RAW 264.7 macrophage expressing CRY2PHR-mcherry2-RASAL3 (red; upper panel) and LifeAct-Halo (cyan; lower panel). Activated macrophage had multiple F-actin-rich protrusions (marked by LifeAct) around its perimeter, as shown by pink arrows (No Opto-inhibition). Next, RASAL3 was recruited to protrusions by applying 488 nm laser near it, as shown by the dashed white box. Protrusions disappeared at site of recruitment and only formed at the cortex where RASAL3 was mostly absent (Local Opto-inhibition). Pink arrows highlight these protrusions. Finally, RASAL3 was recruited over the entire periphery, after 488 nm light was applied all around the cell. It shrank the cell and LifeAct-containing protrusions soon disappeared (Global Opto-inhibition). Images are representative of many cells from atleast three independent experiments. Time in min:sec format. Scale bars represent 5 µm. **(B)** Time-lapse confocal images of HL-60 neutrophil expressing CRY2PHR-mcherry2-RASAL3 (red; upper panel) and LifeActmiRFP703 (cyan; lower panel). Unpolarized, non-migratory neutrophil had multiple F-actin-rich protrusions around its perimeter, as shown by pink arrows (No Opto-inhibition). RASAL3 recruitment over the entire periphery, after 488 nm light was applied all around the cell, caused the cell to shrink and LifeAct-containing protrusions soon disappeared (Global Opto-inhibition). Pink arrows highlight cellular protrusions. Region of blue light illumination is shown by the dashed white box. Images are representative of many cells from atleast three independent experiments. Time in min:sec format. Scale bars represent 5 µm. **(C)** Cartoon illustrating RASAL3-mediated phenomenon seen in (A, ‘Global Opto-inhibition’) and (B).

**Figure S8.**
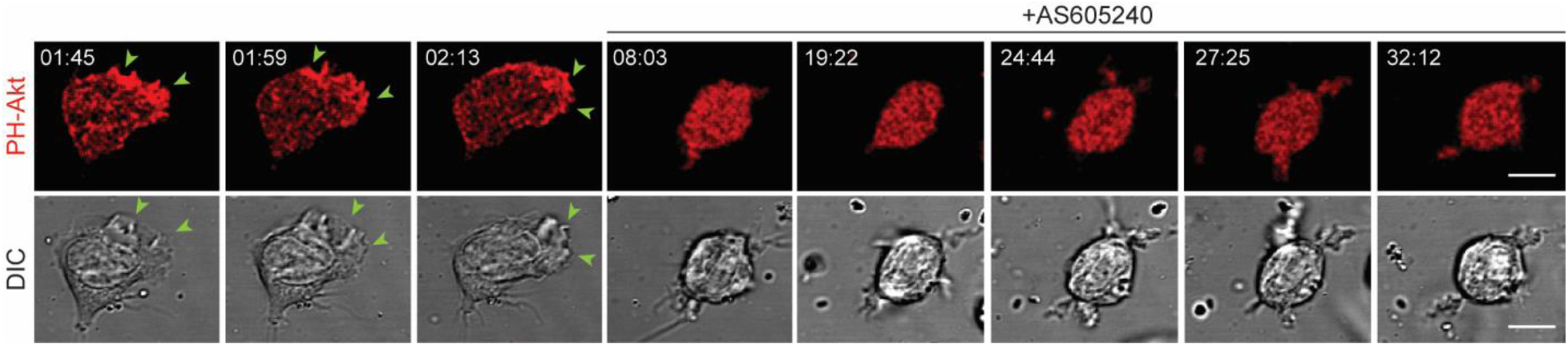
AS605240 treatment depletes PIP3 in neutrophils. Time-lapse confocal images of AS605240-treated HL-60 neutrophil expressing RFP-PH-Akt (red). Green arrows show PH-Akt localization or PIP3 accumulation on protrusions (DIC). Images are representative of many cells from atleast three independent experiments. Time in min:sec format. Scale bars represent 5 µm.

**Figure S9.**
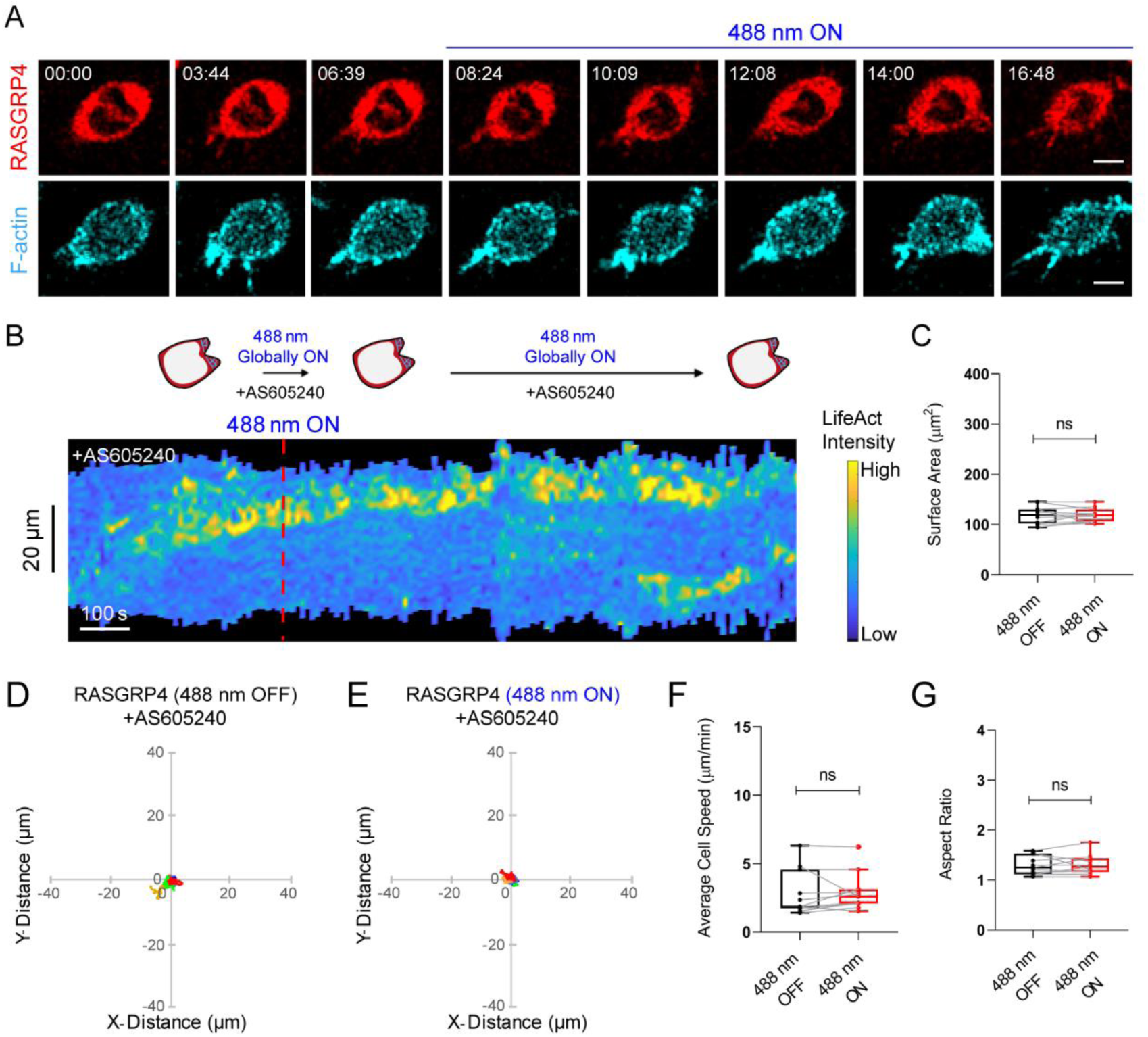
Cytosolic RasGRP4 does not overcome PI3K inhibition. **(A)** Time-lapse confocal images of AS605240-treated HL-60 neutrophil expressing CRY2PHR-mcherry2-RasGRP4 (red; upper panel) and LifeActmiRFP703 (cyan; lower panel), before or after 488 nm laser was switched on globally. No appreciable RasGRP4 recruitment was observed presumably due to low expression of CIBN-CAAX membrane anchor. Time in min:sec format. Scale bars represent 5 µm. **(B)** Representative kymograph of cortical LifeAct intensity in AS605240-treated RasGRP4-expressing neutrophil before or after 488 nm laser was turned on. A linear color map shows that blue is the lowest LifeAct intensity whereas yellow is the highest. Duration of the kymograph is 17 mins. Cartoon depicts recruitment, F-actin polymerization or cell shape status corresponding to the kymograph. **(C)** Box-and-whisker plot of neutrophil surface area before (black) or after (red) 488 nm laser was turned on. n_c_=11 from atleast 3 independent experiments; ns denotes P>0.05 (Wilcoxon-Mann-Whitney rank sum test). Centroid tracks of AS605240-treated neutrophils (n_c_=11) showing random motility before **(D)** or after **(E)** 488 nm laser was turned on. Each track lasts atleast 5 mins and was reset to same origin. Box-and-whisker plots of neutrophil speed **(F)** and aspect ratio **(G)** before (black) or after (red) 488 nm laser was switched on. n_c_=11 from atleast 3 independent experiments; ns denotes P>0.05 (Wilcoxon-Mann-Whitney rank sum test).

**Figure S10.**
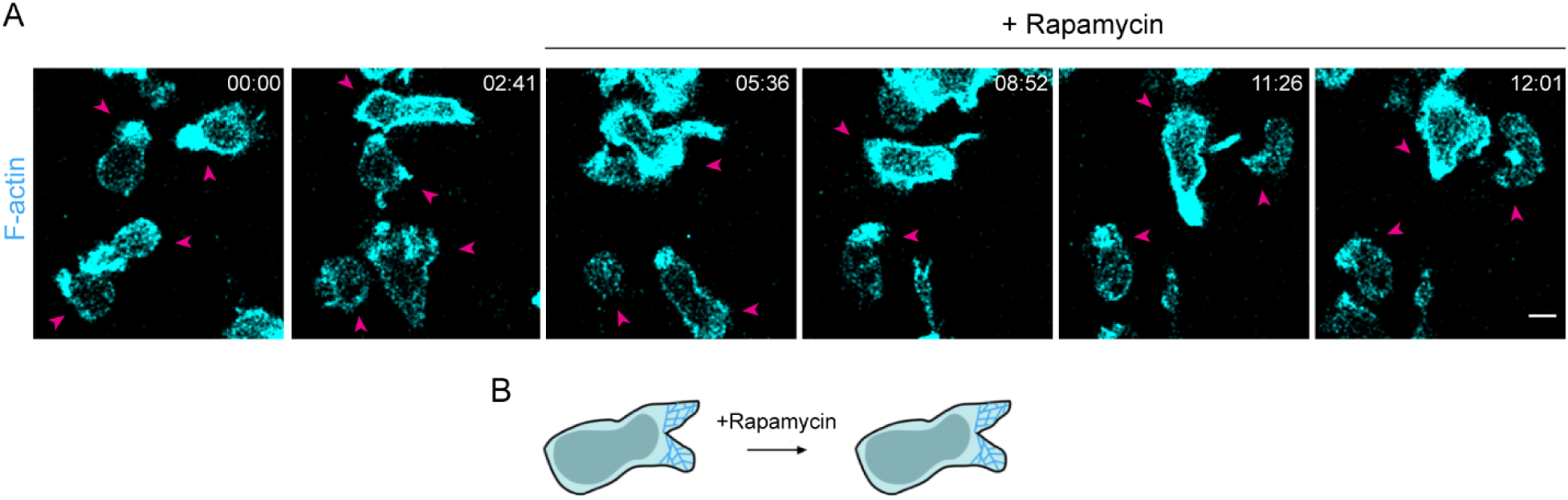
Rapamycin inhibition of mTorC1 does not cause cytoskeletal defects. **(A)** Time-lapse confocal images of rapamycin-treated HL-60 neutrophils expressing LifeActmiRFP703 (cyan). Pink arrows show representative cells which have not lost polarity or show any deformity in shape or size after rapamycin treatment. Images are representative of many cells from atleast three independent experiments. Time in min:sec format. Scale bars represent 5 µm. **(B)** Cartoon depicting rapamycin treatment does not affect neutrophil shape or F-actin polymerization.

**Figure S11.**
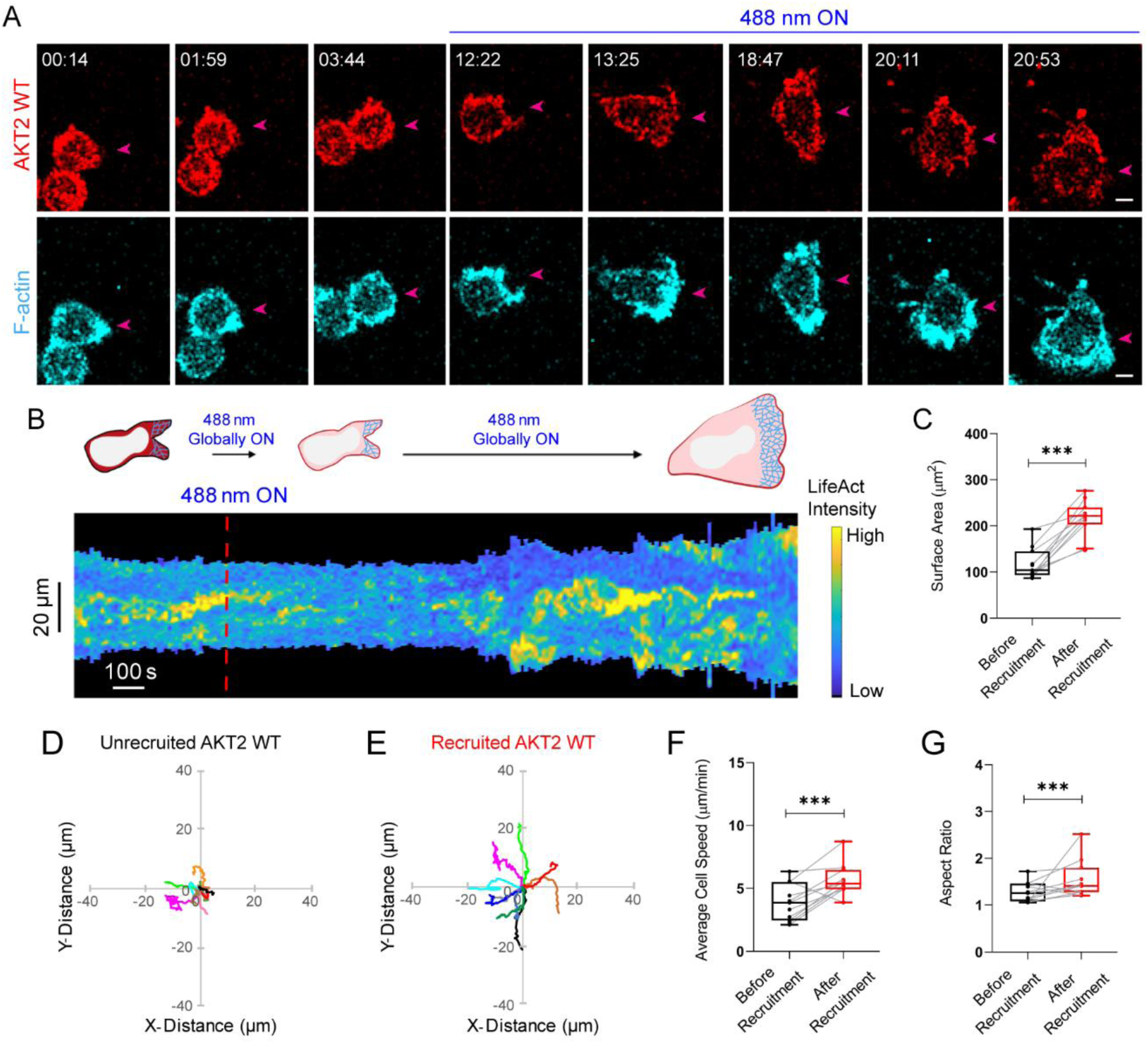
Global recruitment of Akt2 improves neutrophil polarity and migration. **(A)** Time-lapse confocal images of differentiated HL-60 neutrophil expressing CRY2PHR-mcherry2-AKT2 WT (red; upper panel) and LifeActmiRFP703 (cyan; lower panel), before or after 488 nm laser was switched on globally. Pink arrows denote representative cell. Time in min:sec format. Scale bars represent 5 µm. **(B)** Representative kymograph of cortical LifeAct intensity in Akt2 WT-expressing neutrophil before or after 488 nm laser was turned on. A linear color map shows that blue is the lowest LifeAct intensity whereas yellow is the highest. Duration of the kymograph is 23 mins. Cartoon depicts recruitment, F-actin polymerization or cell shape status corresponding to the kymograph. **(C)** Box-and-whisker plot of neutrophil surface area before (black) or after (red) CRY2PHR-mcherry2-Akt2 WT membrane recruitment. n_c_=11 from atleast 3 independent experiments; asterisks indicate significant difference, ***P ≤ 0.001 (Wilcoxon-Mann-Whitney rank sum test). Centroid tracks of neutrophils (n_c_=11) showing random motility before **(D)** or after **(E)** Akt2 WT membrane recruitment. Each track lasts atleast 5 mins and was reset to same origin. Box-and-whisker plots of neutrophil speed **(F)** and aspect ratio **(G)** before (black) or after (red) Akt2 WT membrane recruitment. n_c_=11 from atleast 3 independent experiments; asterisks indicate significant difference, ***P ≤ 0.001 (Wilcoxon-Mann-Whitney rank sum test).

**Figure S12.**
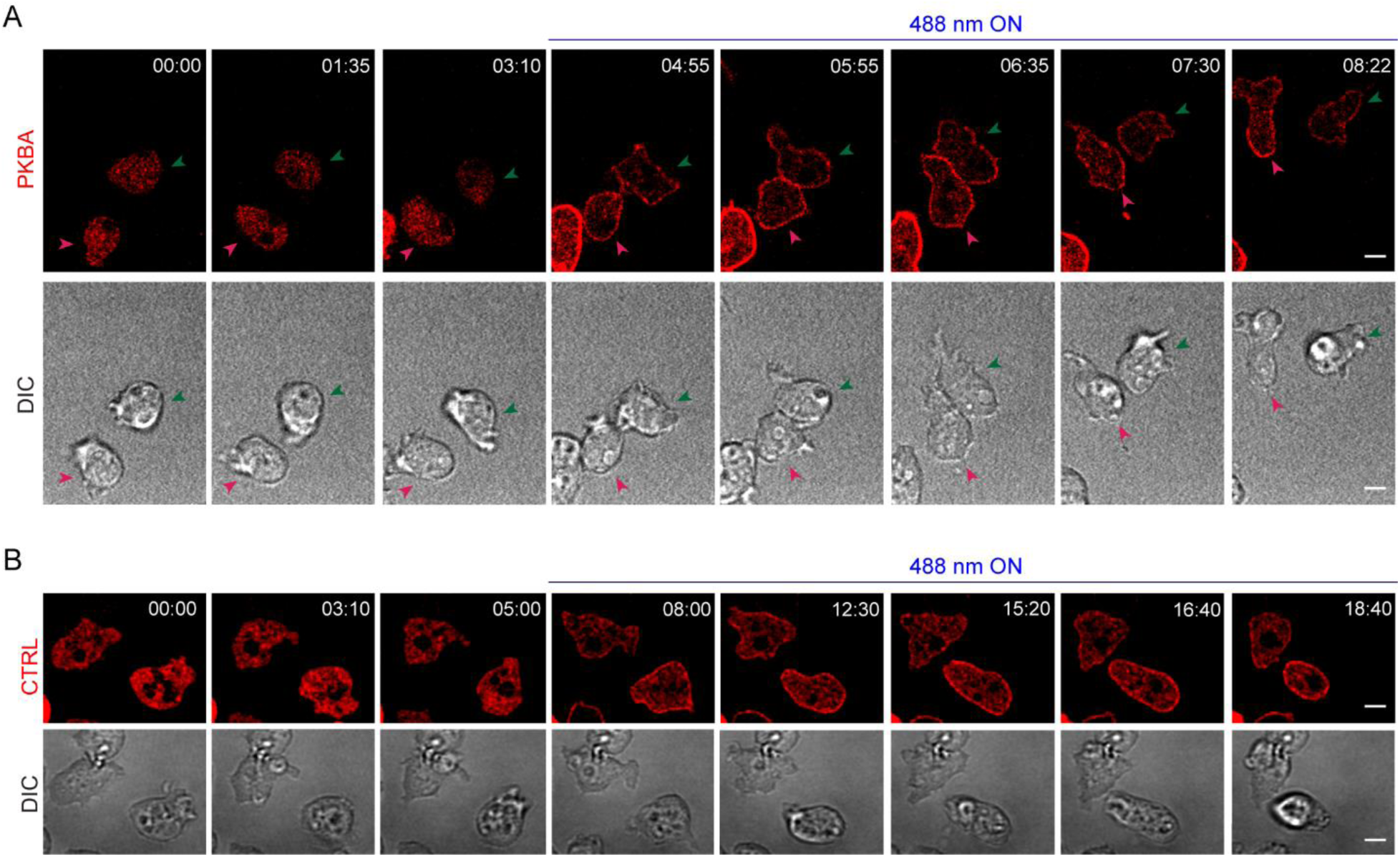
Global recruitment of PKBA polarizes *Dictyostelium* migration. **(A)** Time-lapse confocal images of vegetative *Dictyostelium* expressing tgRFPt-SSPB R73Q-PKBA (red; upper panel), before or after 488 nm laser was switched on globally. Within minutes of PKBA recruitment, previously non-polarized cells formed sustained protrusions and started migrating (DIC; lower panel). Pink and green arrows are provided to demarcate the two cells throughout the images. These are representative of many cells from atleast three independent experiments. Time in min:sec format. Scale bars represent 5 µm. **(B)** Time-lapse confocal images of vegetative *Dictyostelium* expressing tgRFPt-SSPB R73Q-CTRL (control, red; upper panel), before or after 488 nm laser was switched on globally. Non-polarized cells do not polarize or migrate once CTRL is recruited to the membrane (DIC; lower panel). These images are representative of many cells from atleast three independent experiments. Time in min:sec format. Scale bars represent 5 µm.

**Figure S13.**
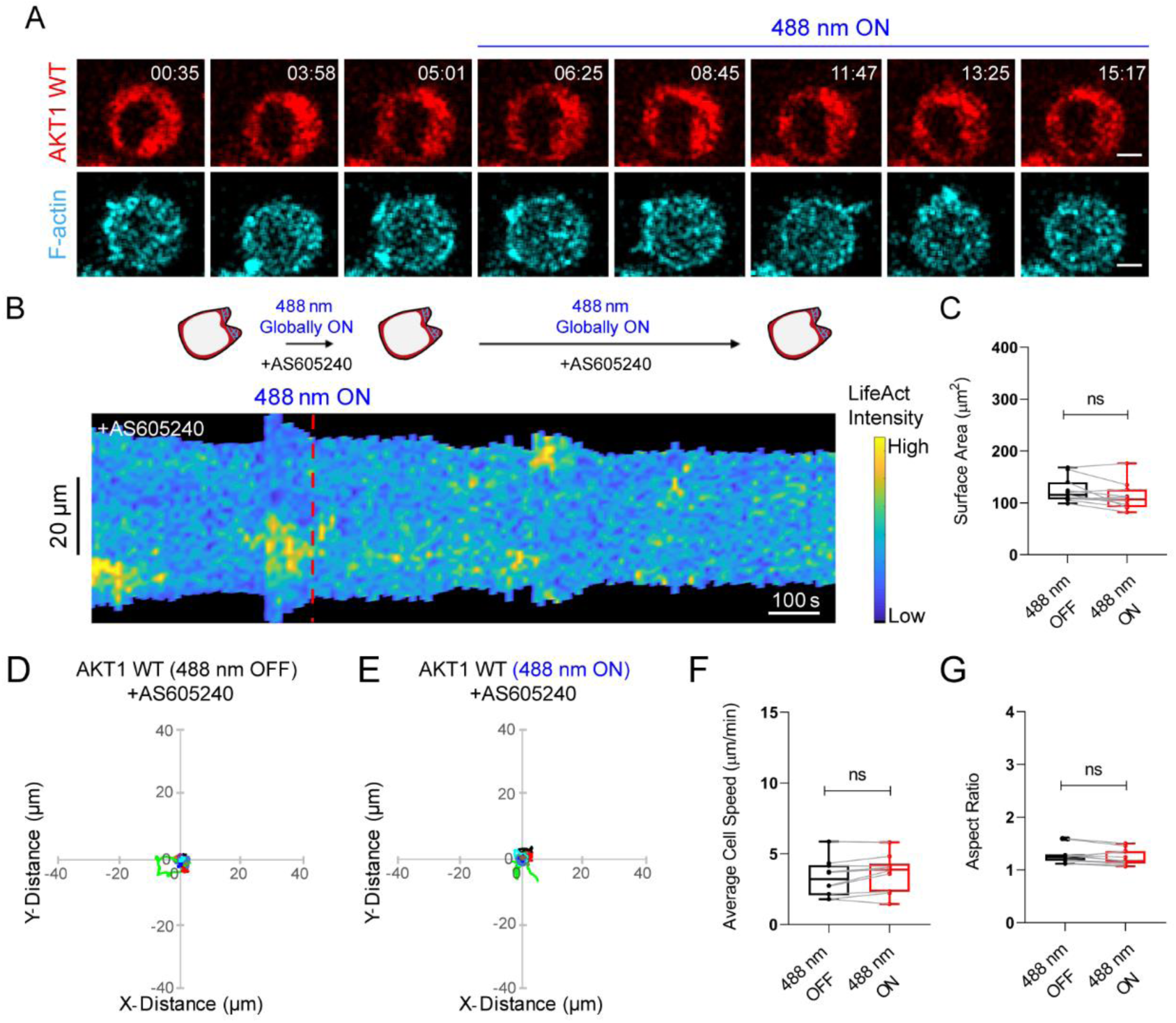
Cytosolic Akt1 does not overcome PI3K inhibition. **(A)** Time-lapse confocal images of AS605240-treated HL-60 neutrophil expressing CRY2PHR-mcherry2-Akt1 WT (red; upper panel) and LifeActmiRFP703 (cyan; lower panel), before or after 488 nm laser was switched on globally. No appreciable Akt1 WT recruitment was observed presumably due to low expression of CIBN-CAAX membrane anchor. Time in min:sec format. Scale bars represent 5 µm. **(B)** Representative kymograph of cortical LifeAct intensity in AS605240-treated Akt1 WT-expressing neutrophil before or after 488 nm laser was turned on. A linear color map shows that blue is the lowest LifeAct intensity whereas yellow is the highest. Duration of the kymograph is 16 mins. Cartoon depicts recruitment, F-actin polymerization or cell shape status corresponding to the kymograph. **(C)** Box-and-whisker plot of neutrophil surface area before (black) or after (red) 488 nm laser was turned on. n_c_=10 from atleast 3 independent experiments; ns denotes P>0.05 (Wilcoxon-Mann-Whitney rank sum test). Centroid tracks of AS605240-treated neutrophils (n_c_=10) showing random motility before **(D)** or after **(E)** 488 nm laser was turned on. Each track lasts atleast 5 mins and was reset to same origin. Box-and-whisker plots of neutrophil speed **(F)** and aspect ratio **(G)** before (black) or after (red) 488 nm laser was switched on. n_c_=10 from atleast 3 independent experiments; ns denotes P>0.05 (Wilcoxon-Mann-Whitney rank sum test).

### Supplementary Tables

**Table S1.**
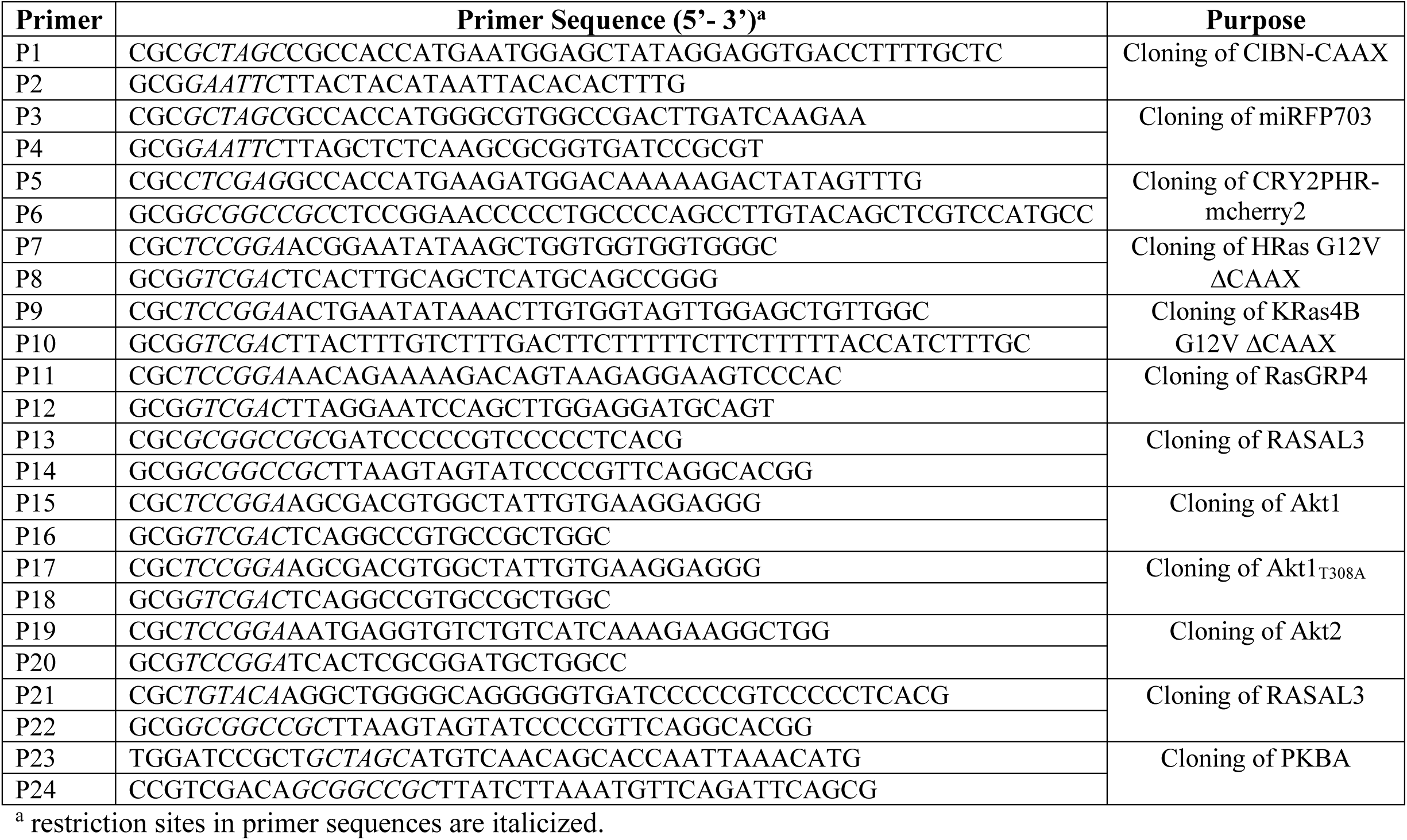
Oligonucleotides used in this paper.

### Supplementary Movie Legends

**Video S1. Global recruitment of constitutively active HRas generates lamellipodium and improves polarity, spreading, and migration in neutrophils.** Time-lapse confocal microscopy of differentiated HL-60 neutrophil expressing CRY2PHR-mcherry2-HRas G12V ΔCAAX (red; left panel) and LifeActmiRFP703 (cyan; right panel), before or after 488 nm laser was switched on globally. The membrane anchor, untagged CIBN-CAAX, was expressed. Pink arrows in both panels denote cell of interest. Top left corner shows time in min:sec format. To initiate recruitment (red; left panel), the laser was switched on at ‘08:10’ once ‘488 nm ON’ appears at the top of the video. Neutrophils were not exposed to any chemoattractant during the course of this experiment. Scale bar represents 5 µm.

**Video S2. Global recruitment of constitutively active KRas4B increases neutrophil polarity, spreading, and migration.** Time-lapse confocal microscopy of differentiated HL-60 neutrophil expressing CRY2PHR-mcherry2-KRas4B G12V ΔCAAX (red; left panel) and LifeActmiRFP703 (cyan; right panel), before or after 488 nm laser was switched on globally. The membrane anchor, untagged CIBN-CAAX, was expressed. Top left corner shows time in min:sec format. To initiate recruitment (red; left panel), the laser was switched on at ‘05:01’ once ‘488 nm ON’ appears at the top of the video. Neutrophils were not exposed to any chemoattractant during the course of this experiment. Scale bar represents 5 µm.

**Video S3. Global recruitment of constitutively active KRas4B improves neutrophil polarity, spreading, and migration in absence of receptor activated G-protein signaling.** Time-lapse confocal microscopy of pertussis toxin- and gallein-treated differentiated HL-60 neutrophil expressing CRY2PHR-mcherry2-KRas4B G12V ΔCAAX (red; left panel) and LifeActmiRFP703 (cyan; right panel), before or after 488 nm laser was switched on globally. The membrane anchor, untagged CIBN-CAAX, was expressed. Neutrophils were pre-treated with pertussis toxin (PTX, Gα_i_ inhibitor) and gallein (Gβγ inhibitor) for 20 hrs and 10 mins, respectively, before imaging, and kept in presence of both inhibitors throughout imaging. Pink arrows in both panels denote representative cell. Top left corner shows time in min:sec format. To initiate recruitment (red; left panel), the laser was switched on at ‘03:23’ once ‘488 nm ON’ appears at the top of the video. Neutrophils were not exposed to any chemoattractant during the course of this experiment. Scale bar represents 5 µm.

**Video S4. Global recruitment of RasGRP4 on neutrophil membrane induces polarity, spreading, and migration.** Time-lapse confocal microscopy of differentiated HL-60 neutrophil expressing CRY2PHR-mcherry2-RasGRP4 (red; left panel) and LifeActmiRFP703 (cyan; right panel), before or after 488 nm laser was switched on globally. The membrane anchor, untagged CIBN-CAAX, was expressed. Top left corner shows time in min:sec format. To initiate recruitment (red; left panel), the laser was switched on at ‘04:47’ once ‘488 nm ON’ appears at the top of the video. Neutrophils were not exposed to any chemoattractant during the course of this experiment. Scale bar represents 5 µm.

**Video S5. Global recruitment of CRY2PHR on neutrophil membrane has no effect on polarity, spreading, and migration.** Time-lapse confocal microscopy of differentiated HL-60 neutrophil expressing CRY2PHR-mcherry2-CTRL(control, red; left panel) and LifeActmiRFP703 (cyan; right panel), before or after 488 nm laser was switched on globally. The membrane anchor, untagged CIBN-CAAX, was expressed. Top left corner shows time in min:sec format. To initiate recruitment (red; left panel), the laser was switched on at ‘05:08’ once ‘488 nm ON’ appears at the top of the video. Neutrophils were not exposed to any chemoattractant during the course of this experiment. Scale bar represents 5 µm.

**Video S6. Local recruitment of RasGRP4 to the cell rear generates protrusions and reverses pre-existing polarity in neutrophils.** Time-lapse confocal microscopy of differentiated HL-60 neutrophil expressing CRY2PHR-mcherry2-RasGRP4 (red; left panel) and LifeActmiRFP703 (cyan; right panel). The membrane anchor, untagged CIBN-CAAX, was expressed. RasGRP4 was recruited exclusively to the back of the cell by transiently applying 488 nm laser near it, when the solid yellow box appears in the red panel. Nascent protrusions originating from site of RasGRP4 recruitment is marked by solid yellow box in the blue panel. Top right corner shows time in min:sec format. Neutrophils were not exposed to any chemoattractant during the course of this experiment. Scale bar represents 5 µm.

**Video S7. Local recruitment of constitutively active HRas to the cell rear creates protrusions and rearranges front-back axis of neutrophils.** Time-lapse confocal microscopy of differentiated HL-60 neutrophil expressing CRY2PHR-mcherry2-HRas G12V ΔCAAX (red; left panel) and LifeActmiRFP703 (cyan; right panel). The membrane anchor, untagged CIBN-CAAX, was expressed. HRas G12V ΔCAAX was recruited exclusively to the back of the cell by transiently applying 488 nm laser near it, when the solid yellow box appears in the red panel. Nascent protrusions originating from site of HRas G12V ΔCAAX recruitment is marked by solid yellow box in the blue panel. Top left corner shows time in min:sec format. Neutrophils were not exposed to any chemoattractant during the course of this experiment. Scale bar represents 5 µm.

**Video S8. Local recruitment of RASAL3 to the cell front inhibits mature protrusions and creates a new back in neutrophils.** Time-lapse confocal microscopy of differentiated HL-60 neutrophil expressing CRY2PHR-mcherry2-RASAL3 (red; left panel) and LifeActmiRFP703 (cyan; right panel). The membrane anchor, untagged CIBN-CAAX, was expressed. RASAL3 was recruited exclusively to the front of the cell by transiently applying 488 nm laser near it, when the solid yellow box appears in the red panel. Disappearance of protrusions at the site of RASAL3 recruitment is marked by solid yellow box in the blue panel. Top left corner shows time in min:sec format. Neutrophils were not exposed to any chemoattractant during the course of this experiment. Scale bar represents 5 µm.

**Video S9. RASAL3 recruitment inhibits protrusive activity in activated macrophages.** Time-lapse confocal microscopy of FKP-(D-Cha)-Cha-r-treated RAW264.7 macrophage expressing CRY2PHR-mcherry2-RASAL3 (red; left panel) and LifeAct-Halo (cyan; right panel). LifeAct-Halo was visualized by standard Janelia Fluor 646 HaloTag labeling protocol. The membrane anchor, untagged CIBN-CAAX, was expressed. Imaging was started within a minute of adding the chemoattractant, FKP-(D-Cha)-Cha-r, to macrophages. RASAL3 was recruited to protrusions by transiently applying 488 nm laser near it, when the solid yellow box appears in the red panel. Disappearance of protrusions at the site of RASAL3 recruitment is marked by solid yellow box in the blue panel. Top left corner shows time in min:sec format. Scale bar represents 5 µm.

**Video S10. RASAL3 recruitment silences protrusions in neutrophils.** Time-lapse confocal microscopy of differentiated HL-60 neutrophil expressing CRY2PHR-mcherry2-RASAL3 (red; left panel) and LifeActmiRFP703 (cyan; right panel). The membrane anchor, untagged CIBN-CAAX, was expressed. RASAL3 is recruited to protrusions by transiently applying 488 nm laser around the cell perimeter, when the solid yellow box appears in the red panel. Disappearance of protrusions at the site of RASAL3 recruitment is marked by solid yellow box in the blue panel. Neutrophils were not exposed to any chemoattractant during the course of this experiment. Top left corner shows time in min:sec format. Scale bar represents 5 µm.

**Video S11. Local recruitment of CRY2PHR to the cell rear does not reverse pre-existing polarity in neutrophils.** Time-lapse confocal microscopy of differentiated HL-60 neutrophil expressing CRY2PHR-mcherry2-CTRL (control, red; left panel) and LifeActmiRFP703 (cyan; right panel). The membrane anchor, untagged CIBN-CAAX, was expressed. CTRL was recruited exclusively to the back of the cell by transiently applying 488 nm laser near it, when the solid yellow box appears in the red panel. No formation of new protrusions at the site of CTRL recruitment is marked by solid yellow box in the blue panel. Top left corner shows time in min:sec format. Neutrophils were not exposed to any chemoattractant during the course of this experiment. Scale bar represents 5 µm.

**Video S12. Global recruitment of RasGRP4 improves neutrophil polarity, spreading, and migration in absence of PI3K activity.** Time-lapse confocal microscopy of AS605240-treated differentiated HL-60 neutrophil expressing CRY2PHR-mcherry2-RasGRP4 (red; left panel) and LifeActmiRFP703 (cyan; right panel), before or after 488 nm laser was switched on globally. The membrane anchor, untagged CIBN-CAAX, was expressed. Pink arrows in both panels denote cell of interest. Neutrophils were pre-treated with AS605240 (PI3K inhibitor) for 10 mins before imaging, and kept in presence of the inhibitor throughout imaging as denoted by ‘+AS605240’ on top of the video. Top left corner shows time in min:sec format. To initiate recruitment (red; left panel), the laser was switched on at ‘03:09’ once ‘488 nm ON’ appears at the top of the video. Neutrophils were not exposed to any chemoattractant during the course of this experiment. Scale bar represents 5 µm.

**Video S13. Global recruitment of RasGRP4 cannot activate neutrophil polarity, spreading, and migration in absence of mTorC2 activity.** Time-lapse confocal microscopy of PP242-treated differentiated HL-60 neutrophil expressing CRY2PHR-mcherry2-RasGRP4 (red; left panel) and LifeActmiRFP703 (cyan; right panel), before or after 488 nm laser was switched on globally. The membrane anchor, untagged CIBN-CAAX, was expressed. Neutrophils were pre-treated with PP242 (mTor inhibitor) for 10 mins before imaging, and kept in presence of the inhibitor throughout imaging. Top left corner shows time in min:sec format. To initiate recruitment (red; left panel), the laser was switched on at ‘06:11’ once ‘488 nm ON’ appears at the top of the video. Neutrophils were not exposed to any chemoattractant during the course of this experiment. Scale bar represents 5 µm.

**Video S14. Local recruitment of Akt1 to the cell rear generates protrusions and reverses pre-existing polarity in neutrophils.** Time-lapse confocal microscopy of differentiated HL-60 neutrophil expressing CRY2PHR-mcherry2-AKT1 WT (red; left panel) and LifeActmiRFP703 (cyan; right panel). The membrane anchor, untagged CIBN-CAAX, was expressed. AKT1 WT was recruited exclusively to the back of the cell by transiently applying 488 nm laser near it, when the solid yellow box appears in the red panel. Nascent protrusions originating from site of AKT1 WT recruitment is marked by solid yellow box in the blue panel. Top left corner shows time in min:sec format. Neutrophils were not exposed to any chemoattractant during the course of this experiment. Scale bar represents 5 µm.

**Video S15. Local recruitment of inactive Akt1_T308A_ to the cell rear does not reverse pre-existing polarity in neutrophils.** Time-lapse confocal microscopy of differentiated HL-60 neutrophil expressing CRY2PHR-mcherry2-AKT1 T308A (control, red; left panel) and LifeActmiRFP703 (cyan; right panel). The membrane anchor, untagged CIBN-CAAX, was expressed. AKT1 T308A was recruited exclusively to the back of the cell by transiently applying 488 nm laser near it, when the solid yellow box appears in the red panel. No formation of new protrusions at the site of AKT1 T308A recruitment is marked by solid yellow box in the blue panel. Top left corner shows time in min:sec format. Neutrophils were not exposed to any chemoattractant during the course of this experiment. Scale bar represents 5 µm.

**Video S16. Global recruitment of Akt2 improves neutrophil polarity, spreading, and migration.** Time-lapse confocal microscopy of differentiated HL-60 neutrophil expressing CRY2PHR-mcherry2-AKT2 WT (red; left panel) and LifeActmiRFP703 (cyan; right panel), before or after 488 nm laser was switched on globally. The membrane anchor, untagged CIBN-CAAX, was expressed. Pink arrows in both panels denote cell of interest. Top left corner shows time in min:sec format. To initiate recruitment (red; left panel), the laser was switched on at ‘04:26’ once ‘488 nm ON’ appears at the top of the video. Neutrophils were not exposed to any chemoattractant during the course of this experiment. Scale bar represents 5 µm.

**Video S17. Global recruitment of PKBA polarizes *Dictyostelium* and improves migration.** Time-lapse confocal microscopy of vegetative *Dictyostelium* expressing tgRFPt-SSPB R73Q-PKBA (red; left panel), before or after 488 nm laser was switched on globally. Right panel is DIC. The membrane anchor, Venus-iLID-CAAX, was expressed but not shown here. Pink and green arrows in both panels denote cells of interest. Top left corner shows time in min:sec format. To initiate recruitment (red; left panel), the laser was switched on at ‘04:05’ once ‘488 nm ON’ appears at the top of the video. *Dictyostelium* cells were not exposed to any chemoattractant during the course of this experiment. Scale bar represents 5 µm.

**Video S18. Global recruitment of SspB does not polarize *Dictyostelium* or improve migration.** Time-lapse confocal microscopy of vegetative *Dictyostelium* expressing tgRFPt-SSPB R73Q-CTRL (control, red; left panel), before or after 488 nm laser was switched on globally. Right panel is DIC. The membrane anchor, Venus-iLID-CAAX, was expressed but not shown here. Top left corner shows time in min:sec format. To initiate recruitment (red; left panel), the laser was switched on at ‘05:10’ once ‘488 nm ON’ appears at the top of the video. *Dictyostelium* cells were not exposed to any chemoattractant during the course of this experiment. Scale bar represents 5 µm.

**Video S19. Global recruitment of Akt1 activates neutrophil polarity, spreading, and migration in absence of PI3K activity.** Time-lapse confocal microscopy of AS605240-treated differentiated HL-60 neutrophil expressing CRY2PHR-mcherry2-AKT1 WT (red; left panel) and LifeActmiRFP703 (cyan; right panel), before or after 488 nm laser was switched on globally. The membrane anchor, untagged CIBN-CAAX, was expressed. Pink arrows in both panels denote cell of interest. Neutrophils were pre-treated with AS605240 (PI3K inhibitor) for 10 mins before imaging, and kept in presence of the inhibitor throughout imaging. Top left corner shows time in min:sec format. To initiate recruitment (red; left panel), the laser was switched on at ‘08:24’ once ‘488 nm ON’ appears at the top of the video. Neutrophils were not exposed to any chemoattractant during the course of this experiment. Scale bar represents 5 µm.

**Video S20. Global recruitment of inactive Akt1_T308A_ cannot overcome PI3K inhibition.** Time-lapse confocal microscopy of AS605240-treated differentiated HL-60 neutrophil expressing CRY2PHR-mcherry2-AKT1 T308A (red; left panel) and LifeActmiRFP703 (cyan; right panel), before or after 488 nm laser was switched on globally. The membrane anchor, untagged CIBN-CAAX, was expressed. Neutrophils were pre-treated with AS605240 (PI3K inhibitor) for 10 mins before imaging, and kept in presence of the inhibitor throughout imaging. Top left corner shows time in min:sec format. To initiate recruitment (red; left panel), the laser was switched on at ‘05:50’ once ‘488 nm ON’ appears at the top of the video. Neutrophils were not exposed to any chemoattractant during the course of this experiment. Scale bar represents 5 µm.

## References

1. Luster, A.D., Alon, R., and von Andrian, U.H. (2005). Immune cell migration in inflammation: present and future therapeutic targets. Nat Immunol 6, 1182–1190. 10.1038/ni1275.

2. Mantovani, A., Cassatella, M.A., Costantini, C., and Jaillon, S. (2011). Neutrophils in the activation and regulation of innate and adaptive immunity. Nat Rev Immunol 11, 519–531. 10.1038/nri3024.

3. Nourshargh, S., and Alon, R. (2014). Leukocyte migration into inflamed tissues. Immunity 41, 694–707. 10.1016/j.immuni.2014.10.008.

4. Artemenko, Y., Axiotakis, L., Jr., Borleis, J., Iglesias, P.A., and Devreotes, P.N. (2016). Chemical and mechanical stimuli act on common signal transduction and cytoskeletal networks. Proc Natl Acad Sci U S A 113, E7500–E7509. 10.1073/pnas.1608767113.

5. Devreotes, P.N., Bhattacharya, S., Edwards, M., Iglesias, P.A., Lampert, T., and Miao, Y. (2017). Excitable Signal Transduction Networks in Directed Cell Migration. Annu Rev Cell Dev Biol 33, 103–125. 10.1146/annurev-cellbio-100616-060739.

6. Sasaki, A.T., Janetopoulos, C., Lee, S., Charest, P.G., Takeda, K., Sundheimer, L.W., Meili, R., Devreotes, P.N., and Firtel, R.A. (2007). G protein-independent Ras/PI3K/F-actin circuit regulates basic cell motility. J Cell Biol 178, 185–191. 10.1083/jcb.200611138.

7. Tsai, T.Y., Collins, S.R., Chan, C.K., Hadjitheodorou, A., Lam, P.Y., Lou, S.S., Yang, H.W., Jorgensen, J., Ellett, F., Irimia, D., et al. (2019). Efficient Front-Rear Coupling in Neutrophil Chemotaxis by Dynamic Myosin II Localization. Dev Cell 49, 189–205 e186. 10.1016/j.devcel.2019.03.025.

8. Wang, Y., Ku, C.J., Zhang, E.R., Artyukhin, A.B., Weiner, O.D., Wu, L.F., and Altschuler, S.J. (2013). Identifying network motifs that buffer front-to-back signaling in polarized neutrophils. Cell Rep 3, 1607–1616. 10.1016/j.celrep.2013.04.009.

9. Weiner, O.D., Neilsen, P.O., Prestwich, G.D., Kirschner, M.W., Cantley, L.C., and Bourne, H.R. (2002). A PtdInsP(3)- and Rho GTPase-mediated positive feedback loop regulates neutrophil polarity. Nat Cell Biol 4, 509–513. 10.1038/ncb811.

10. Weiner, O.D., Servant, G., Welch, M.D., Mitchison, T.J., Sedat, J.W., and Bourne, H.R. (1999). Spatial control of actin polymerization during neutrophil chemotaxis. Nat Cell Biol 1, 75–81. 10.1038/10042.

11. Yang, X., Dormann, D., Munsterberg, A.E., and Weijer, C.J. (2002). Cell movement patterns during gastrulation in the chick are controlled by positive and negative chemotaxis mediated by FGF4 and FGF8. Dev Cell 3, 425–437. 10.1016/s1534-5807(02)00256-3.

12. Janssen, E., and Geha, R.S. (2019). Primary immunodeficiencies caused by mutations in actin regulatory proteins. Immunol Rev 287, 121–134. 10.1111/imr.12716.

13. SenGupta, S., Parent, C.A., and Bear, J.E. (2021). The principles of directed cell migration. Nat Rev Mol Cell Biol 22, 529–547. 10.1038/s41580-021-00366-6.

14. Kolaczkowska, E., and Kubes, P. (2013). Neutrophil recruitment and function in health and inflammation. Nat Rev Immunol 13, 159–175. 10.1038/nri3399.

15. Kruger, P., Saffarzadeh, M., Weber, A.N., Rieber, N., Radsak, M., von Bernuth, H., Benarafa, C., Roos, D., Skokowa, J., and Hartl, D. (2015). Neutrophils: Between host defence, immune modulation, and tissue injury. PLoS Pathog 11, e1004651. 10.1371/journal.ppat.1004651.

16. Artemenko, Y., Lampert, T.J., and Devreotes, P.N. (2014). Moving towards a paradigm: common mechanisms of chemotactic signaling in Dictyostelium and mammalian leukocytes. Cell Mol Life Sci 71, 3711–3747. 10.1007/s00018-014-1638-8.

17. Bagorda, A., and Parent, C.A. (2008). Eukaryotic chemotaxis at a glance. J Cell Sci 121, 2621–2624. 10.1242/jcs.018077.

18. Bell, G.R.R., Rincon, E., Akdogan, E., and Collins, S.R. (2021). Optogenetic control of receptors reveals distinct roles for actin- and Cdc42-dependent negative signals in chemotactic signal processing. Nat Commun 12, 6148. 10.1038/s41467-021-26371-z.

19. Benard, V., Bohl, B.P., and Bokoch, G.M. (1999). Characterization of rac and cdc42 activation in chemoattractant-stimulated human neutrophils using a novel assay for active GTPases. J Biol Chem 274, 13198–13204. 10.1074/jbc.274.19.13198.

20. Graziano, B.R., Gong, D., Anderson, K.E., Pipathsouk, A., Goldberg, A.R., and Weiner, O.D. (2017). A module for Rac temporal signal integration revealed with optogenetics. J Cell Biol 216, 2515–2531. 10.1083/jcb.201604113.

21. Graziano, B.R., Town, J.P., Sitarska, E., Nagy, T.L., Fosnaric, M., Penic, S., Iglic, A., Kralj-Iglic, V., Gov, N.S., Diz-Munoz, A., and Weiner, O.D. (2019). Cell confinement reveals a branched-actin independent circuit for neutrophil polarity. PLoS Biol 17, e3000457. 10.1371/journal.pbio.3000457.

22. Hannigan, M., Zhan, L., Li, Z., Ai, Y., Wu, D., and Huang, C.K. (2002). Neutrophils lacking phosphoinositide 3-kinase gamma show loss of directionality during N-formyl-Met-Leu-Phe-induced chemotaxis. Proc Natl Acad Sci U S A 99, 3603–3608. 10.1073/pnas.052010699.

23. Inoue, T., and Meyer, T. (2008). Synthetic activation of endogenous PI3K and Rac identifies an AND-gate switch for cell polarization and migration. PLoS One 3, e3068. 10.1371/journal.pone.0003068.

24. Karunarathne, W.K., Giri, L., Kalyanaraman, V., and Gautam, N. (2013). Optically triggering spatiotemporally confined GPCR activity in a cell and programming neurite initiation and extension. Proc Natl Acad Sci U S A 110, E1565–1574. 10.1073/pnas.1220697110.

25. O’Neill, P.R., Castillo-Badillo, J.A., Meshik, X., Kalyanaraman, V., Melgarejo, K., and Gautam, N. (2018). Membrane Flow Drives an Adhesion-Independent Amoeboid Cell Migration Mode. Dev Cell 46, 9–22 e24. 10.1016/j.devcel.2018.05.029.

26. O’Neill, P.R., and Gautam, N. (2014). Subcellular optogenetic inhibition of G proteins generates signaling gradients and cell migration. Mol Biol Cell 25, 2305–2314. 10.1091/mbc.E14-04-0870.

27. O’Neill, P.R., Kalyanaraman, V., and Gautam, N. (2016). Subcellular optogenetic activation of Cdc42 controls local and distal signaling to drive immune cell migration. Mol Biol Cell 27, 1442–1450. 10.1091/mbc.E15-12-0832.

28. Srinivasan, S., Wang, F., Glavas, S., Ott, A., Hofmann, F., Aktories, K., Kalman, D., and Bourne, H.R. (2003). Rac and Cdc42 play distinct roles in regulating PI(3,4,5)P3 and polarity during neutrophil chemotaxis. J Cell Biol 160, 375–385. 10.1083/jcb.200208179.

29. Stephens, L., Milne, L., and Hawkins, P. (2008). Moving towards a better understanding of chemotaxis. Curr Biol 18, R485–494. 10.1016/j.cub.2008.04.048.

30. Van Keymeulen, A., Wong, K., Knight, Z.A., Govaerts, C., Hahn, K.M., Shokat, K.M., and Bourne, H.R. (2006). To stabilize neutrophil polarity, PIP3 and Cdc42 augment RhoA activity at the back as well as signals at the front. J Cell Biol 174, 437–445. 10.1083/jcb.200604113.

31. Weiner, O.D., Marganski, W.A., Wu, L.F., Altschuler, S.J., and Kirschner, M.W. (2007). An actin-based wave generator organizes cell motility. PLoS Biol 5, e221. 10.1371/journal.pbio.0050221.

32. Weiner, O.D., Rentel, M.C., Ott, A., Brown, G.E., Jedrychowski, M., Yaffe, M.B., Gygi, S.P., Cantley, L.C., Bourne, H.R., and Kirschner, M.W. (2006). Hem-1 complexes are essential for Rac activation, actin polymerization, and myosin regulation during neutrophil chemotaxis. PLoS Biol 4, e38. 10.1371/journal.pbio.0040038.

33. Yang, H.W., Collins, S.R., and Meyer, T. (2016). Locally excitable Cdc42 signals steer cells during chemotaxis. Nat Cell Biol 18, 191–201. 10.1038/ncb3292.

34. Suire, S., Condliffe, A.M., Ferguson, G.J., Ellson, C.D., Guillou, H., Davidson, K., Welch, H., Coadwell, J., Turner, M., Chilvers, E.R., et al. (2006). Gbetagammas and the Ras binding domain of p110gamma are both important regulators of PI(3)Kgamma signalling in neutrophils. Nat Cell Biol 8, 1303–1309. 10.1038/ncb1494.

35. Zigmond, S.H. (1974). Mechanisms of sensing chemical gradients by polymorphonuclear leukocytes. Nature 249, 450–452. 10.1038/249450a0.

36. Thevathasan, J.V., Tan, E., Zheng, H., Lin, Y.C., Li, Y., Inoue, T., and Fivaz, M. (2013). The small GTPase HRas shapes local PI3K signals through positive feedback and regulates persistent membrane extension in migrating fibroblasts. Mol Biol Cell 24, 2228–2237. 10.1091/mbc.E12-12-0905.

37. al-Aoukaty, A., Rolstad, B., and Maghazachi, A.A. (1999). Recruitment of pleckstrin and phosphoinositide 3-kinase gamma into the cell membranes, and their association with G beta gamma after activation of NK cells with chemokines. J Immunol 162, 3249–3255.

38. Hirsch, E., Katanaev, V.L., Garlanda, C., Azzolino, O., Pirola, L., Silengo, L., Sozzani, S., Mantovani, A., Altruda, F., and Wymann, M.P. (2000). Central role for G protein-coupled phosphoinositide 3-kinase gamma in inflammation. Science 287, 1049–1053. 10.1126/science.287.5455.1049.

39. Knall, C., Worthen, G.S., and Johnson, G.L. (1997). Interleukin 8-stimulated phosphatidylinositol-3-kinase activity regulates the migration of human neutrophils independent of extracellular signal-regulated kinase and p38 mitogen-activated protein kinases. Proc Natl Acad Sci U S A 94, 3052–3057. 10.1073/pnas.94.7.3052.

40. Li, Z., Jiang, H., Xie, W., Zhang, Z., Smrcka, A.V., and Wu, D. (2000). Roles of PLC-beta2 and -beta3 and PI3Kgamma in chemoattractant-mediated signal transduction. Science 287, 1046–1049. 10.1126/science.287.5455.1046.

41. Nishio, M., Watanabe, K., Sasaki, J., Taya, C., Takasuga, S., Iizuka, R., Balla, T., Yamazaki, M., Watanabe, H., Itoh, R., et al. (2007). Control of cell polarity and motility by the PtdIns(3,4,5)P3 phosphatase SHIP1. Nat Cell Biol 9, 36–44. 10.1038/ncb1515.

42. Pacold, M.E., Suire, S., Perisic, O., Lara-Gonzalez, S., Davis, C.T., Walker, E.H., Hawkins, P.T., Stephens, L., Eccleston, J.F., and Williams, R.L. (2000). Crystal structure and functional analysis of Ras binding to its effector phosphoinositide 3-kinase gamma. Cell 103, 931–943. 10.1016/s0092-8674(00)00196-3.

43. Sasaki, T., Irie-Sasaki, J., Jones, R.G., Oliveira-dos-Santos, A.J., Stanford, W.L., Bolon, B., Wakeham, A., Itie, A., Bouchard, D., Kozieradzki, I., et al. (2000). Function of PI3Kgamma in thymocyte development, T cell activation, and neutrophil migration. Science 287, 1040–1046. 10.1126/science.287.5455.1040.

44. Vicente-Manzanares, M., Rey, M., Jones, D.R., Sancho, D., Mellado, M., Rodriguez-Frade, J.M., del Pozo, M.A., Yanez-Mo, M., de Ana, A.M., Martinez, A.C., et al. (1999). Involvement of phosphatidylinositol 3-kinase in stromal cell-derived factor-1 alpha-induced lymphocyte polarization and chemotaxis. J Immunol 163, 4001–4012.

45. Chen, J., Tang, H., Hay, N., Xu, J., and Ye, R.D. (2010). Akt isoforms differentially regulate neutrophil functions. Blood 115, 4237–4246. 10.1182/blood-2009-11-255323.

46. Liu, G., Bi, Y., Wang, R., Shen, B., Zhang, Y., Yang, H., Wang, X., Liu, H., Lu, Y., and Han, F. (2013). Kinase AKT1 negatively controls neutrophil recruitment and function in mice. J Immunol 191, 2680–2690. 10.4049/jimmunol.1300736.

47. Servant, G., Weiner, O.D., Herzmark, P., Balla, T., Sedat, J.W., and Bourne, H.R. (2000). Polarization of chemoattractant receptor signaling during neutrophil chemotaxis. Science 287, 1037–1040. 10.1126/science.287.5455.1037.

48. Yagi, M., Kantarci, A., Iwata, T., Omori, K., Ayilavarapu, S., Ito, K., Hasturk, H., and Van Dyke, T.E. (2009). PDK1 regulates chemotaxis in human neutrophils. J Dent Res 88, 1119–1124. 10.1177/0022034509349402.

49. Worthen, G.S., Avdi, N., Buhl, A.M., Suzuki, N., and Johnson, G.L. (1994). FMLP activates Ras and Raf in human neutrophils. Potential role in activation of MAP kinase. J Clin Invest 94, 815–823. 10.1172/JCI117401.

50. Suire, S., Lecureuil, C., Anderson, K.E., Damoulakis, G., Niewczas, I., Davidson, K., Guillou, H., Pan, D., Jonathan, C., Phillip, T.H., and Stephens, L. (2012). GPCR activation of Ras and PI3Kc in neutrophils depends on PLCb2/b3 and the RasGEF RasGRP4. EMBO J 31, 3118–3129. 10.1038/emboj.2012.167.

51. Baffi, T.R., Lorden, G., Wozniak, J.M., Feichtner, A., Yeung, W., Kornev, A.P., King, C.C., Del Rio, J.C., Limaye, A.J., Bogomolovas, J., et al. (2021). mTORC2 controls the activity of PKC and Akt by phosphorylating a conserved TOR interaction motif. Sci Signal 14. 10.1126/scisignal.abe4509.

52. He, Y., Li, D., Cook, S.L., Yoon, M.S., Kapoor, A., Rao, C.V., Kenis, P.J., Chen, J., and Wang, F. (2013). Mammalian target of rapamycin and Rictor control neutrophil chemotaxis by regulating Rac/Cdc42 activity and the actin cytoskeleton. Mol Biol Cell 24, 3369–3380. 10.1091/mbc.E13-07-0405.

53. Liu, L., Das, S., Losert, W., and Parent, C.A. (2010). mTORC2 regulates neutrophil chemotaxis in a cAMP- and RhoA-dependent fashion. Dev Cell 19, 845–857. 10.1016/j.devcel.2010.11.004.

54. Liu, L., Gritz, D., and Parent, C.A. (2014). PKCbetaII acts downstream of chemoattractant receptors and mTORC2 to regulate cAMP production and myosin II activity in neutrophils. Mol Biol Cell 25, 1446–1457. 10.1091/mbc.E14-01-0037.

55. Liu, L., and Parent, C.A. (2011). Review series: TOR kinase complexes and cell migration. J Cell Biol 194, 815–824. 10.1083/jcb.201102090.

56. Pylayeva-Gupta, Y., Grabocka, E., and Bar-Sagi, D. (2011). RAS oncogenes: weaving a tumorigenic web. Nat Rev Cancer 11, 761–774. 10.1038/nrc3106.

57. Senoo, H., Murata, D., Wai, M., Arai, K., Iwata, W., Sesaki, H., and Iijima, M. (2021). KARATE: PKA-induced KRAS4B-RHOA-mTORC2 supercomplex phosphorylates AKT in insulin signaling and glucose homeostasis. Mol Cell 81, 4622–4634 e4628. 10.1016/j.molcel.2021.09.001.

58. Zheng, L., Eckerdal, J., Dimitrijevic, I., and Andersson, T. (1997). Chemotactic peptide-induced activation of Ras in human neutrophils is associated with inhibition of p120-GAP activity. J Biol Chem 272, 23448–23454. 10.1074/jbc.272.37.23448.

59. Xu, X., Wen, X., Moosa, A., Bhimani, S., and Jin, T. (2021). Ras inhibitor CAPRI enables neutrophil-like cells to chemotax through a higher-concentration range of gradients. Proc Natl Acad Sci U S A 118. 10.1073/pnas.2002162118.

60. Arai, Y., Shibata, T., Matsuoka, S., Sato, M.J., Yanagida, T., and Ueda, M. (2010). Self-organization of the phosphatidylinositol lipids signaling system for random cell migration. Proc Natl Acad Sci U S A 107, 12399–12404. 10.1073/pnas.0908278107.

61. Cai, H., Das, S., Kamimura, Y., Long, Y., Parent, C.A., and Devreotes, P.N. (2010). Ras-mediated activation of the TORC2-PKB pathway is critical for chemotaxis. J Cell Biol 190, 233–245. 10.1083/jcb.201001129.

62. Fukushima, S., Matsuoka, S., and Ueda, M. (2019). Excitable dynamics of Ras triggers spontaneous symmetry breaking of PIP3 signaling in motile cells. J Cell Sci 132. 10.1242/jcs.224121.

63. Kae, H., Lim, C.J., Spiegelman, G.B., and Weeks, G. (2004). Chemoattractant-induced Ras activation during Dictyostelium aggregation. EMBO Rep 5, 602–606. 10.1038/sj.embor.7400151.

64. Kortholt, A., Kataria, R., Keizer-Gunnink, I., Van Egmond, W.N., Khanna, A., and Van Haastert, P.J. (2011). Dictyostelium chemotaxis: essential Ras activation and accessory signalling pathways for amplification. EMBO Rep 12, 1273–1279. 10.1038/embor.2011.210.

65. Li, X., Edwards, M., Swaney, K.F., Singh, N., Bhattacharya, S., Borleis, J., Long, Y., Iglesias, P.A., Chen, J., and Devreotes, P.N. (2018). Mutually inhibitory Ras-PI(3,4)P2 feedback loops mediate cell migration. Proc Natl Acad Sci U S A 115, E9125–E9134. 10.1073/pnas.1809039115.

66. Li, X., Miao, Y., Pal, D.S., and Devreotes, P.N. (2020). Excitable networks controlling cell migration during development and disease. Semin Cell Dev Biol 100, 133–142. 10.1016/j.semcdb.2019.11.001.

67. Pal, D.S., Li, X., Banerjee, T., Miao, Y., and Devreotes, P.N. (2019). The excitable signal transduction networks: movers and shapers of eukaryotic cell migration. Int J Dev Biol 63, 407–416. 10.1387/ijdb.190265pd.

68. Sasaki, A.T., Chun, C., Takeda, K., and Firtel, R.A. (2004). Localized Ras signaling at the leading edge regulates PI3K, cell polarity, and directional cell movement. J Cell Biol 167, 505–518. 10.1083/jcb.200406177.

69. Wu, L., Valkema, R., Van Haastert, P.J., and Devreotes, P.N. (1995). The G protein beta subunit is essential for multiple responses to chemoattractants in Dictyostelium. J Cell Biol 129, 1667–1675. 10.1083/jcb.129.6.1667.

70. Bolourani, P., Spiegelman, G., and Weeks, G. (2010). Determinants of RasC specificity during Dictyostelium aggregation. J Biol Chem 285, 41374–41379. 10.1074/jbc.M110.181115.

71. Chubb, J.R., Wilkins, A., Thomas, G.M., and Insall, R.H. (2000). The Dictyostelium RasS protein is required for macropinocytosis, phagocytosis and the control of cell movement. J Cell Sci 113 (Pt 4), 709–719. 10.1242/jcs.113.4.709.

72. Khosla, M., Spiegelman, G.B., Insall, R., and Weeks, G. (2000). Functional overlap of the dictyostelium RasG, RasD and RasB proteins. J Cell Sci 113 (Pt 8), 1427–1434. 10.1242/jcs.113.8.1427.

73. El-Brolosy, M.A., and Stainier, D.Y.R. (2017). Genetic compensation: A phenomenon in search of mechanisms. PLoS Genet 13, e1006780. 10.1371/journal.pgen.1006780.

74. Rossi, A., Kontarakis, Z., Gerri, C., Nolte, H., Holper, S., Kruger, M., and Stainier, D.Y. (2015). Genetic compensation induced by deleterious mutations but not gene knockdowns. Nature 524, 230–233. 10.1038/nature14580.

75. Zhang, S., Charest, P.G., and Firtel, R.A. (2008). Spatiotemporal regulation of Ras activity provides directional sensing. Curr Biol 18, 1587–1593. 10.1016/j.cub.2008.08.069.

76. Miao, Y., Bhattacharya, S., Banerjee, T., Abubaker-Sharif, B., Long, Y., Inoue, T., Iglesias, P.A., and Devreotes, P.N. (2019). Wave patterns organize cellular protrusions and control cortical dynamics. Mol Syst Biol 15, e8585. 10.15252/msb.20188585.

77. Miao, Y., Bhattacharya, S., Edwards, M., Cai, H., Inoue, T., Iglesias, P.A., and Devreotes, P.N. (2017). Altering the threshold of an excitable signal transduction network changes cell migratory modes. Nat Cell Biol 19, 329–340. 10.1038/ncb3495.

78. Ross, B., Mehta, S., and Zhang, J. (2016). Molecular tools for acute spatiotemporal manipulation of signal transduction. Curr Opin Chem Biol 34, 135–142. 10.1016/j.cbpa.2016.08.012.

79. Arrieumerlou, C., and Meyer, T. (2005). A local coupling model and compass parameter for eukaryotic chemotaxis. Dev Cell 8, 215–227. 10.1016/j.devcel.2004.12.007.

80. Gerisch, G., and Keller, H.U. (1981). Chemotactic reorientation of granulocytes stimulated with micropipettes containing fMet-Leu-Phe. J Cell Sci 52, 1–10. 10.1242/jcs.52.1.1.

81. Hadjitheodorou, A., Bell, G.R.R., Ellett, F., Shastry, S., Irimia, D., Collins, S.R., and Theriot, J.A. (2021). Directional reorientation of migrating neutrophils is limited by suppression of receptor input signaling at the cell rear through myosin II activity. Nat Commun 12, 6619. 10.1038/s41467-021-26622-z.

82. Olguin-Olguin, A., Aalto, A., Maugis, B., Boquet-Pujadas, A., Hoffmann, D., Ermlich, L., Betz, T., Gov, N.S., Reichman-Fried, M., and Raz, E. (2021). Chemokine-biased robust self-organizing polarization of migrating cells in vivo. Proc Natl Acad Sci U S A 118. 10.1073/pnas.2018480118.

83. Surve, C.R., To, J.Y., Malik, S., Kim, M., and Smrcka, A.V. (2016). Dynamic regulation of neutrophil polarity and migration by the heterotrimeric G protein subunits Galphai-GTP and Gbetagamma. Sci Signal 9, ra22. 10.1126/scisignal.aad8163.

84. Tang, M., Wang, M., Shi, C., Iglesias, P.A., Devreotes, P.N., and Huang, C.H. (2014). Evolutionarily conserved coupling of adaptive and excitable networks mediates eukaryotic chemotaxis. Nat Commun 5, 5175. 10.1038/ncomms6175.

85. Wang, M.J., Artemenko, Y., Cai, W.J., Iglesias, P.A., and Devreotes, P.N. (2014). The directional response of chemotactic cells depends on a balance between cytoskeletal architecture and the external gradient. Cell Rep 9, 1110–1121. 10.1016/j.celrep.2014.09.047.

86. Milligan, G. (2003). Constitutive activity and inverse agonists of G protein-coupled receptors: a current perspective. Mol Pharmacol 64, 1271–1276. 10.1124/mol.64.6.1271.

87. Kirimanjeswara, G.S., Agosto, L.M., Kennett, M.J., Bjornstad, O.N., and Harvill, E.T. (2005). Pertussis toxin inhibits neutrophil recruitment to delay antibody-mediated clearance of Bordetella pertussis. J Clin Invest 115, 3594–3601. 10.1172/JCI24609.

88. Lehmann, D.M., Seneviratne, A.M., and Smrcka, A.V. (2008). Small molecule disruption of G protein beta gamma subunit signaling inhibits neutrophil chemotaxis and inflammation. Mol Pharmacol 73, 410–418. 10.1124/mol.107.041780.

89. Shan, D., Chen, L., Wang, D., Tan, Y.C., Gu, J.L., and Huang, X.Y. (2006). The G protein G alpha(13) is required for growth factor-induced cell migration. Dev Cell 10, 707–718. 10.1016/j.devcel.2006.03.014.

90. Surve, C.R., Lehmann, D., and Smrcka, A.V. (2014). A chemical biology approach demonstrates G protein betagamma subunits are sufficient to mediate directional neutrophil chemotaxis. J Biol Chem 289, 17791–17801. 10.1074/jbc.M114.576827.

91. Reuther, G.W., Lambert, Q.T., Rebhun, J.F., Caligiuri, M.A., Quilliam, L.A., and Der, C.J. (2002). RasGRP4 is a novel Ras activator isolated from acute myeloid leukemia. J Biol Chem 277, 30508–30514. 10.1074/jbc.M111330200.

92. Yang, Y., Li, L., Wong, G.W., Krilis, S.A., Madhusudhan, M.S., Sali, A., and Stevens, R.L. (2002). RasGRP4, a new mast cell-restricted Ras guanine nucleotide-releasing protein with calcium- and diacylglycerol-binding motifs. Identification of defective variants of this signaling protein in asthma, mastocytosis, and mast cell leukemia patients and demonstration of the importance of RasGRP4 in mast cell development and function. J Biol Chem 277, 25756–25774. 10.1074/jbc.M202575200.

93. Rottner, K., and Schaks, M. (2019). Assembling actin filaments for protrusion. Curr Opin Cell Biol 56, 53–63. 10.1016/j.ceb.2018.09.004.

94. Hetrick, B., Han, M.S., Helgeson, L.A., and Nolen, B.J. (2013). Small molecules CK-666 and CK-869 inhibit actin-related protein 2/3 complex by blocking an activating conformational change. Chem Biol 20, 701–712. 10.1016/j.chembiol.2013.03.019.

95. Saito, S., Cao, D.Y., Victor, A.R., Peng, Z., Wu, H.Y., and Okwan-Duodu, D. (2021). RASAL3 Is a Putative RasGAP Modulating Inflammatory Response by Neutrophils. Front Immunol 12, 744300. 10.3389/fimmu.2021.744300.

96. Azzi, J., Moore, R.F., Elyaman, W., Mounayar, M., El Haddad, N., Yang, S., Jurewicz, M., Takakura, A., Petrelli, A., Fiorina, P., et al. (2012). The novel therapeutic effect of phosphoinositide 3-kinase-gamma inhibitor AS605240 in autoimmune diabetes. Diabetes 61, 1509–1518. 10.2337/db11-0134.

97. Camps, M., Ruckle, T., Ji, H., Ardissone, V., Rintelen, F., Shaw, J., Ferrandi, C., Chabert, C., Gillieron, C., Francon, B., et al. (2005). Blockade of PI3Kgamma suppresses joint inflammation and damage in mouse models of rheumatoid arthritis. Nat Med 11, 936–943. 10.1038/nm1284.

98. Hoang, B., Frost, P., Shi, Y., Belanger, E., Benavides, A., Pezeshkpour, G., Cappia, S., Guglielmelli, T., Gera, J., and Lichtenstein, A. (2010). Targeting TORC2 in multiple myeloma with a new mTOR kinase inhibitor. Blood 116, 4560–4568. 10.1182/blood-2010-05-285726.

99. Aoki, M., Batista, O., Bellacosa, A., Tsichlis, P., and Vogt, P.K. (1998). The akt kinase: molecular determinants of oncogenicity. Proc Natl Acad Sci U S A 95, 14950–14955. 10.1073/pnas.95.25.14950.

100. Meili, R., Ellsworth, C., and Firtel, R.A. (2000). A novel Akt/PKB-related kinase is essential for morphogenesis in Dictyostelium. Curr Biol 10, 708–717. 10.1016/s0960-9822(00)00536-4.

101. Meili, R., Ellsworth, C., Lee, S., Reddy, T.B., Ma, H., and Firtel, R.A. (1999). Chemoattractant-mediated transient activation and membrane localization of Akt/PKB is required for efficient chemotaxis to cAMP in Dictyostelium. EMBO J 18, 2092–2105. 10.1093/emboj/18.8.2092.

102. Hoxhaj, G., and Manning, B.D. (2020). The PI3K-AKT network at the interface of oncogenic signalling and cancer metabolism. Nat Rev Cancer 20, 74–88. 10.1038/s41568-019-0216-7.

103. Houk, A.R., Jilkine, A., Mejean, C.O., Boltyanskiy, R., Dufresne, E.R., Angenent, S.B., Altschuler, S.J., Wu, L.F., and Weiner, O.D. (2012). Membrane tension maintains cell polarity by confining signals to the leading edge during neutrophil migration. Cell 148, 175–188. 10.1016/j.cell.2011.10.050.

104. Diz-Munoz, A., Fletcher, D.A., and Weiner, O.D. (2013). Use the force: membrane tension as an organizer of cell shape and motility. Trends Cell Biol 23, 47–53. 10.1016/j.tcb.2012.09.006.

105. Diz-Munoz, A., Thurley, K., Chintamen, S., Altschuler, S.J., Wu, L.F., Fletcher, D.A., and Weiner, O.D. (2016). Membrane Tension Acts Through PLD2 and mTORC2 to Limit Actin Network Assembly During Neutrophil Migration. PLoS Biol 14, e1002474. 10.1371/journal.pbio.1002474.

106. Adachi, R., Krilis, S.A., Nigrovic, P.A., Hamilton, M.J., Chung, K., Thakurdas, S.M., Boyce, J.A., Anderson, P., and Stevens, R.L. (2012). Ras guanine nucleotide-releasing protein-4 (RasGRP4) involvement in experimental arthritis and colitis. J Biol Chem 287, 20047–20055. 10.1074/jbc.M112.360388.

107. Jun, J.E., Rubio, I., and Roose, J.P. (2013). Regulation of ras exchange factors and cellular localization of ras activation by lipid messengers in T cells. Front Immunol 4, 239. 10.3389/fimmu.2013.00239.

108. Saito, S., Kawamura, T., Higuchi, M., Kobayashi, T., Yoshita-Takahashi, M., Yamazaki, M., Abe, M., Sakimura, K., Kanda, Y., Kawamura, H., et al. (2015). RASAL3, a novel hematopoietic RasGAP protein, regulates the number and functions of NKT cells. Eur J Immunol 45, 1512–1523. 10.1002/eji.201444977.

109. Zhu, M., Fuller, D.M., and Zhang, W. (2012). The role of Ras guanine nucleotide releasing protein 4 in Fc epsilonRI-mediated signaling, mast cell function, and T cell development. J Biol Chem 287, 8135–8143. 10.1074/jbc.M111.320580.

110. Ferguson, G.J., Milne, L., Kulkarni, S., Sasaki, T., Walker, S., Andrews, S., Crabbe, T., Finan, P., Jones, G., Jackson, S., et al. (2007). PI(3)Kgamma has an important context-dependent role in neutrophil chemokinesis. Nat Cell Biol 9, 86–91. 10.1038/ncb1517.

111. Funamoto, S., Meili, R., Lee, S., Parry, L., and Firtel, R.A. (2002). Spatial and temporal regulation of 3-phosphoinositides by PI 3-kinase and PTEN mediates chemotaxis. Cell 109, 611–623. 10.1016/s0092-8674(02)00755-9.

112. Iijima, M., and Devreotes, P. (2002). Tumor suppressor PTEN mediates sensing of chemoattractant gradients. Cell 109, 599–610. 10.1016/s0092-8674(02)00745-6.

113. Wessels, D., Lusche, D.F., Kuhl, S., Heid, P., and Soll, D.R. (2007). PTEN plays a role in the suppression of lateral pseudopod formation during Dictyostelium motility and chemotaxis. J Cell Sci 120, 2517–2531. 10.1242/jcs.010876.

114. Veltman, D.M., Keizer-Gunnik, I., and Van Haastert, P.J. (2008). Four key signaling pathways mediating chemotaxis in Dictyostelium discoideum. J Cell Biol 180, 747–753. 10.1083/jcb.200709180.

115. Banerjee, T., Biswas, D., Pal, D.S., Miao, Y., Iglesias, P.A., and Devreotes, P.N. (2022). Spatiotemporal dynamics of membrane surface charge regulates cell polarity and migration. Nat Cell Biol 24, 1499–1515. 10.1038/s41556-022-00997-7.

116. Lien, E.C., Dibble, C.C., and Toker, A. (2017). PI3K signaling in cancer: beyond AKT. Curr Opin Cell Biol 45, 62–71. 10.1016/j.ceb.2017.02.007.

117. Kamimura, Y., and Devreotes, P.N. (2010). Phosphoinositide-dependent protein kinase (PDK) activity regulates phosphatidylinositol 3,4,5-trisphosphate-dependent and - independent protein kinase B activation and chemotaxis. J Biol Chem 285, 7938–7946. 10.1074/jbc.M109.089235.

118. Guha, P., Reilly, L., Semenza, E.R., Abramson, E., Mishra, S., Sei, Y., Wank, S.A., Donowitz, M., and Snyder, S.H. (2020). Loss of PI3-kinase activity of inositol polyphosphate multikinase impairs PDK1-mediated AKT activation, cell migration and intestinal homeostasis. bioRxiv, 2020.2012.2018.423145. 10.1101/2020.12.18.423145.

119. Chen, W.S., Xu, P.Z., Gottlob, K., Chen, M.L., Sokol, K., Shiyanova, T., Roninson, I., Weng, W., Suzuki, R., Tobe, K., et al. (2001). Growth retardation and increased apoptosis in mice with homozygous disruption of the Akt1 gene. Genes Dev 15, 2203–2208. 10.1101/gad.913901.

120. Garofalo, R.S., Orena, S.J., Rafidi, K., Torchia, A.J., Stock, J.L., Hildebrandt, A.L., Coskran, T., Black, S.C., Brees, D.J., Wicks, J.R., et al. (2003). Severe diabetes, age-dependent loss of adipose tissue, and mild growth deficiency in mice lacking Akt2/PKB beta. J Clin Invest 112, 197–208. 10.1172/JCI16885.

121. Ackah, E., Yu, J., Zoellner, S., Iwakiri, Y., Skurk, C., Shibata, R., Ouchi, N., Easton, R.M., Galasso, G., Birnbaum, M.J., et al. (2005). Akt1/protein kinase Balpha is critical for ischemic and VEGF-mediated angiogenesis. J Clin Invest 115, 2119–2127. 10.1172/JCI24726.

122. Arboleda, M.J., Lyons, J.F., Kabbinavar, F.F., Bray, M.R., Snow, B.E., Ayala, R., Danino, M., Karlan, B.Y., and Slamon, D.J. (2003). Overexpression of AKT2/protein kinase Bbeta leads to up-regulation of beta1 integrins, increased invasion, and metastasis of human breast and ovarian cancer cells. Cancer Res 63, 196–206.

123. Higuchi, M., Masuyama, N., Fukui, Y., Suzuki, A., and Gotoh, Y. (2001). Akt mediates Rac/Cdc42-regulated cell motility in growth factor-stimulated cells and in invasive PTEN knockout cells. Curr Biol 11, 1958–1962. 10.1016/s0960-9822(01)00599-1.

124. Irie, H.Y., Pearline, R.V., Grueneberg, D., Hsia, M., Ravichandran, P., Kothari, N., Natesan, S., and Brugge, J.S. (2005). Distinct roles of Akt1 and Akt2 in regulating cell migration and epithelial-mesenchymal transition. J Cell Biol 171, 1023–1034. 10.1083/jcb.200505087.

125. Yoeli-Lerner, M., Yiu, G.K., Rabinovitz, I., Erhardt, P., Jauliac, S., and Toker, A. (2005). Akt blocks breast cancer cell motility and invasion through the transcription factor NFAT. Mol Cell 20, 539–550. 10.1016/j.molcel.2005.10.033.

126. de Oliveira, S., Rosowski, E.E., and Huttenlocher, A. (2016). Neutrophil migration in infection and wound repair: going forward in reverse. Nat Rev Immunol 16, 378–391. 10.1038/nri.2016.49.

127. Millius, A., and Weiner, O.D. (2010). Manipulation of neutrophil-like HL-60 cells for the study of directed cell migration. Methods Mol Biol 591, 147–158. 10.1007/978-1-60761-404-3_9.

128. Rincon, E., Rocha-Gregg, B.L., and Collins, S.R. (2018). A map of gene expression in neutrophil-like cell lines. BMC Genomics 19, 573. 10.1186/s12864-018-4957-6.

129. Meshik, X., O’Neill, P.R., and Gautam, N. (2018). Optogenetic Control of Cell Migration. Methods Mol Biol 1749, 313–324. 10.1007/978-1-4939-7701-7_22.

130. Yusa, K., Rad, R., Takeda, J., and Bradley, A. (2009). Generation of transgene-free induced pluripotent mouse stem cells by the piggyBac transposon. Nat Methods 6, 363–369. 10.1038/nmeth.1323.

131. Zhan, H., Bhattacharya, S., Cai, H., Iglesias, P.A., Huang, C.H., and Devreotes, P.N. (2020). An Excitable Ras/PI3K/ERK Signaling Network Controls Migration and Oncogenic Transformation in Epithelial Cells. Dev Cell 54, 608–623 e605. 10.1016/j.devcel.2020.08.001.

132. Li, X., Pal, D.S., Biswas, D., Iglesias, P.A., and Devreotes, P.N. (2021). Reverse fountain flow of phosphatidylinositol-3,4-bisphosphate polarizes migrating cells. EMBO J 40, e105094. 10.15252/embj.2020105094.

133. Chronopoulos, A., Thorpe, S.D., Cortes, E., Lachowski, D., Rice, A.J., Mykuliak, V.V., Rog, T., Lee, D.A., Hytonen, V.P., and Del Rio Hernandez, A.E. (2020). Syndecan-4 tunes cell mechanics by activating the kindlin-integrin-RhoA pathway. Nat Mater 19, 669–678. 10.1038/s41563-019-0567-1.

